# Overall patient’s survival of glioblastoma associated to molecular markers: a pan-proteomic prospective study

**DOI:** 10.1101/2020.11.24.397117

**Authors:** Lauranne Drelich, Marie Duhamel, Maxence Wisztorski, Soulaimane Aboulouard, Jean-Pascal Gimeno, Pierre-Damien Caux, Nina Ogrinc, Patrick Devos, Tristan Cardon, Michael Weller, Fabienne Escande, Fahed Zairi, Claude-Alain Maurage, Isabelle Fournier, Emilie Le Rhun, Michel Salzet

## Abstract

Molecular heterogeneities are a key feature of glioblastoma (GBM) pathology impeding patient’s stratification and leading to high discrepancies between patients mean survivals. Here, we established a molecular classification of GBM tumors using a pan-proteomic analysis. Then, we identified, from our proteomic data, 2 clusters of biomarkers associated with good or bad patient survival from 46 IDH wild-type GBMs. Three molecular groups have been identified and associated with systemic biology analyses. Group A tumors exhibit neurogenesis characteristics and tumorigenesis. Group B shows a strong immune cell signature and express poor prognosis markers while group C tumors are characterized by an anti-viral signature and tumor growth proteins. 124 proteins were found statistically different based on patient’s survival times, of which 10 are issued from alternative AltORF or non-coding RNA. After statistical analysis, a panel of markers associated to higher survival (PPP1R12A, RPS14, HSPD1 and LASP1) and another panel associated to lower survival (ALCAM, ANXA11, MAOB, IP_652563 and IGHM) has been validated by immunofluorescence. Taken together, our data will guide GBM prognosis and help to improve the current GBM classification by stratifying the patients and may open new opportunities for therapeutic development.

**Significance:** Glioblastoma are very heterogeneous tumors with median survivals usually inferior to 20 months. We conducted a pan-proteomics analysis of glioblastoma (GBM) in order to stratify GBM based on the molecular contained. Forty-six GBM cases were classified into three groups where proteins are involved in specific pathways *i.e.* the first group has a neurogenesis signature and is associated with a better prognosis while the second group of patients has an immune profile with a bad prognosis. The third group is more associated to tumorigenesis. We correlated these results with the TCGA data. Finally, we have identified 28 new prognostic markers of GBM and from these 28, a panel of 4 higher and 5 lower survival markers were validated. With these 9 markers in hand, now pathologist can stratify GBM patients and can guide the therapeutic decision.

**Highlights:**

- A novel stratification of glioblastoma based on mass spectrometry was established.
- Three groups with different molecular features and survival were identified.
- This new classification could improve prognostication and may help therapeutic options.
- 8 prognosis markers for oncologist therapeutic decision have been validated.

## Introduction

Glioblastoma (GBM) represents the main malignant primary brain tumor with an incidence of 3.21 per 100,000 population (1). The prognosis is poor with a median survival estimated at 16 months in clinical studies (2–8) and around 12 months in contemporary population-based studies (9). Approximately 5% of patients survive more than 5 years (10). Favorable therapy-independent prognostic factors include lower age and higher neurological performance status at diagnosis. Furthermore, low postoperative residual tumor volume has been associated with improved outcome. In a cohort of 232 patients with centrally confirmed glioblastoma who survived at least 5 years, the median age at diagnosis was 52 years (range 21-77 years) and most patients had a gross total resection initially (11).

Morphological criteria for the diagnosis of GBM according to the World Health Organization (WHO) central nervous system tumor classification of 2016 (12) include mitotic activity, anaplastic nuclear features, microvascular proliferation and necrosis. This definition comprised three groups of tumors, (i) isocitrate dehydrogenase (IDH) wild-type GBM which represents at least 90% of GBM and includes as morphological variants giant cell GBM, gliosarcoma and epithelioid glioblastoma; (ii) IDH-mutant GBM which represents less than 10% of GBM and (iii) glioblastoma *not otherwise specified* (NOS) when the IDH status is unknown. These diagnostic concepts are very likely to be revised in the upcoming revision of the WHO classification based on cIMPACT-NOW (update 3 (13) and 5 (14)) recommendations to allow a molecular definition of GBM based on the detection of EGFR amplification, a +7/−10 genotype and TERT promoter mutation in a IDH wildtype astrocytic tumor (Brat et al. 2018) and by renaming IDH-mutant GBM as IDH-mutant astrocytoma, WHO grade 4 (15–17). Standard treatment of GBM includes maximum safe resection followed by radiotherapy with concomitant and maintenance temozolomide (18).

Efforts to further subclassify glioblastoma have been confined to the genomic, transcriptomic and epigenetic level. In 2008, the Cancer Genome Atlas (TCGA) group delineated three main signaling pathways affected by genetic alterations in GBM, receptor tyrosine kinase/RAS/PI3K, p53 and RB (19,20). Genome methylation profiling in adult patients with IDH wildtype GBM allowed the definition of three epigenetic subtypes, (i) receptor tyrosine kinase (RTK) I often with PDGFR amplification, (ii) RTK II or classic often with EGFR amplification, CDKN2A/B deletion, and PTEN mutation, and (iii) mesenchymal (21). Any clinical relevance of the methylation classes remains controversial. The DNA methylation-based classification of CNS tumors has meanwhile evolved to a comprehensive machine-learning approach that will shape the next revision of the WHO classification (22), resulting also in the delineation of further rare methylation classes of GBM. Prior to the introduction of methylation profiling, a classification of glioblastoma based on transcriptional profiling have revealed four subtypes of GBM: proneural, neural, classic and mesenchymal (23). The neural subtype is no longer maintained, and it appears now that transcriptomic profiles are less homogeneous and stable than genome or methylome classifiers. Despite these efforts, these approached have found limited clinical application and only a few biomarkers are being used in clinic. IDH wild-type glioblastoma, which represents the most aggressive glioma subtype, lack of predictive markers and the stratification of patients would allow to identify specific altered signaling pathways which may be useful for the development of targeted therapies. Proteomic approaches have been less frequently explored, although they can identify and quantify the final product of altered genomics and transcriptomics and may better characterize the activation of specific pathways (24–27). The determination of specific proteomic signatures could help to improve the distinction between the different GBM subtypes and to guide the therapeutic management.

Here, we stratified 46 IDH WT-GBM tumors based on a spatially resolved proteomic approach guided by mass spectrometry imaging. Our strategy provides new insights into intertumoral and intratumoral heterogeneities by taking into account the glioblastoma microenvironment which is of prime importance in tumor development. Based on our pan-proteomic study, 5 prognostic markers were also identified at the outcome of our analysis.

## Materials & Methods

### Patient samples and consent

Patients with newly diagnosed glioblastoma were prospectively enrolled between September 2014 and November 2018 at Lille University Hospital, France. Patients were adult, had no medical history of other cancers or previous cancer treatment, no known genetic disease potentially leading to cancer and no neurodegenerative disease. Tumors samples were processed within 2 hours after sample extraction in the surgery room to limit the risk of degradation of proteins.

Immunohistochemistry analyses, deoxyribonucleic acid (DNA) extraction and quantification of tumor samples are reported in Note S1 (**Supplementary Data 1**).

Approval was obtained from the research ethics committee (ID-RCB 2014-A00185-42) before initiation of the study. The study adhered to the principles of the Declaration of Helsinki and the Guidelines for Good Clinical Practice and is registered at NCT02473484. Informed consent was obtained from patients.

### MALDI mass spectrometry imaging (MALDI MSI)

A Leica CM1510S cryostat (Leica Microsystems, Nanterre, France) was used to cut twelve micrometer sections in order to perform the MALDI MSI analysis (28–32). These tissue sections were deposited on ITO-coated glass slides (LaserBio Labs, Valbonne, France) and vacuum-dried during 15 min. Tissue sections were then soaked in different solutions to remove abundant lipids: (1) 1 min in 70% ethanol, (2) 1 min in 100% ethanol, (3) 1 min in acetone and (4) 30 s in chloroform with concomitant drying between washings. An electrospray nebulizer connected to a syringe pump (flow rate 180 nL/min) was used to uniformly spray a trypsin solution (60 μg/mL in NH4HCO3 50 mM) on the tissue surface for 15 min. ImagePrep (Bruker Daltonics, Bremen, Germany) was used as an incubation chamber by microspraying water heated to 37 °C for 2 h (60 cycles with 2 s spraying, 180 s incubation and 60 s drying using the nitrogen flow). For optimal digestion, a constant humidity atmosphere was maintained inside the spray chamber by filling a small container with 95°C water. After digestion, HCCA/ANI (Lemaire et al., 2006) a solid ionic matrix was deposited using ImagePrep. Briefly, 36 μL of aniline were added to 5 mL of a solution of 10 mg/mL HCCA dissolved in ACN/0.1% TFA aqueous (7:3, v/v). A real-time control of the deposition is performed by monitoring scattered light to obtain a uniform layer of matrix. MALDI MSI experiments were done on an Ultraflex II MALDI-TOF/TOF instrument (Bruker) with a smartbeam II solid state laser. Mass spectra were acquired in positive reflector mode between 800–4000 m/z range. Recorded spectra were averaged from 400 laser shots per pixel acquired at 200Hz laser repletion rate and. with a 70 μm spatial resolution raster.

### MALDI MSI data processing and analysis

The MALDI-MSI data were analyzed using SCiLS Lab software (SCiLS Lab 2019, SCiLS GmbH). Common processing methods for MALDI MSI were applied with a baseline removal using a convolution method and data were normalized using Total Ion Count (TIC) method (33,34). Then, the resulting pre-processing data were clustered to obtain a spatial segmentation using the bisecting k means algorithm (35). Different spatial segmentations were performed. First, an individual segmentation was applied to each tissue separately. Then, the data from all tissues were clustered together to obtain a global segmentation. Briefly, the spatial segmentation consists of grouping all spectra according to their similarity using a clustering algorithm and all pixels of a same cluster are colour coded. To limit the pixel-to-pixel variability, edge-preserving image denoising was applied. Note that a color is arbitrary assigned to a cluster and that several disconnected regions can have the same color, i.e. the same molecular content. The results of segmentation are represented on a dendrogram resulting from a hierarchical clustering. The branches of the dendrogram were defined based on a distance calculation between each cluster. The selection of different branches of the dendrogram will give a segmentation map where regions of distinct molecular composition were differentially color-coded. The individual segmentation provides information concerning the heterogeneity of the tissue section and the global segmentation is used to group patients with a similar molecular signature. For comparison, global segmentation was also performed using the Ward clustering method with IMAGEREVEAL MS Ver.1.1 (Shimadzu). The global spatial segmentation allowed to determine regions of interest (ROIs) which were then be subjected to on-tissue microdigestion followed by microextraction for protein identifications.

### SpiderMass analyses

The global design of the instrument setup has been described (36). Briefly, the system is composed of three parts including a laser system for micro-sampling of tissues set remotely, a transfer line allowing for transfer of the micro-sampled material to the third part, which is the mass spectrometer itself (37). The first part is composed of a tunable wavelength OPO which is tunable from 2.8 μm to 3.1 μm (Radiant version 1.0.1, OPOTEK Inc., Carlsbad, CA, USA) pumped by a pulsed Nd:YAG laser (pulse duration: 5 ns, λ=1064 nm, Quantel, Les Ulis, France). A biocompatible laser fiber (450 μm inner diameter; length of 1 m; Infrared Fiber Systems, Silver Spring, CO, USA) is connected to the laser system output and a handpiece including a 4 cm focusing lens is attached to the end of the laser fiber. The handpiece with a 4cm focusing lens allows the user to hold the system and screen the surface of raw tissues at a resolution of 400 μm. In these experiments the irradiation time was fixed to 10 sec at 4 mJ/pulse laser energy corresponding to a laser fluence of ~3 J/cm2. The laser energy was measured at the focal point of the focusing lens using a power meter (ThorLabs, Maisons-Laffitte, France). The second part of the system corresponds to a 3 meter length transfer line made from a Tygon ND 100-65 tubing (2.4 mm inner diameter, 4 mm outer diameter, Akron, USA). The transfer line is attached on one side onto the laser hand piece at the end of the laser fiber and on its other side directly connected to the mass spectrometer (Xevo, Waters, Manchester, United Kingdom) from which the conventional electrospray source was removed and replaced by an atmospheric pressure interface (37). Each acquisition was accompanied by a 150 μL/min isopropanol infusion. Spectral acquisition was performed both in positive and negative ion resolution mode with a scan time of 1 sec. Prior to SpiderMass analysis, the samples were taken out of the −20°C freezer and thawed to RT for 30 s. The spectral acquisition sequence was composed of 2 or 3 acquisitions using 1-sec irradiation periods. The ROI were selected using the morphological controls and acquired peptide MALDI-MSI data prior to each SpiderMass acquisition in order to ensure that each acquisition was performed on the same histological area (38).

### Classification model construction

For data analysis, all raw data files produced with the SpiderMass instrument were imported into the Abstract Model Builder (AMX v.0.9.2092.0) software. After importation, spectra were first pre-processed. The pre-processing steps include background subtraction, total ion count normalization, lockmass correction and re-binning to a 0.1 or 0.2 Da window. All processed MS spectra obtained from the 30 histologically validated samples were then used to build a principal component analysis and linear discriminant analysis (PCA-LDA) classification model (38). The first step consisted of PCA to reduce data multidimensionality by generating features that explain most of the variance observed. These features were then subjected to supervised analysis using LDA by setting the classes that the model will be based upon. LDA attempts to classify the sample spectra and assess the model by cross validation. Cross-validation was carried out by either using the “20% out” or the “leave one patient out” methods. For the first method, 20% of MS spectra are randomly taken from the total spectra and the model is reconstructed from the remaining 80%. The remaining 20% of spectra are used to interrogate the reconstructed model. The permutation is automatically reiterated for 5 cycles before reporting the cross-validation results. For the second method, the spectra are grouped by patient and left out one by one; at each step the model without the patient is interrogated against this model.

#### Spatially-resolved proteomics

##### On-tissue digestion

A total of 122 ROIs were selected from MALDI-MSI. Spatially resolved microproteomics was performed on the predefined ROIs according to the previously published protocol (39). Briefly, tissue sections of 20μm thick were cut and subjected to different washes to remove lipids. Then, on-tissue digestion is performed using a LysC-trypsin solution (40 μg/mL in Tris-HCl 50mM, pH 8.0). This solution was deposited using a piezoelectric microspotter (CHIP-1000, Shimadzu, CO, Kyoto, Japan) on each ROIs with a total area of 1 mm^2^ (4×4 spots of 200 μm. Enzyme droplet was maintained for a total of 2 h digestion. After enzyme deposition 0.1% TFA was spotted for 25 cycles with 100 pL on each spot/cycle.

##### Microextraction by liquid microjunction

After tissue microdigestion, the triptic peptides were extracted using an automated platform, the TriVersa Nanomate platform (Advion Biosciences Inc., Ithaca, NY, USA) with Liquid Extraction Surface Analysis (LESA) option (39). Briefly, a volume of solvent was aspirated onto a tip and dispensed onto the digested region. The droplet formed was maintained between the tip and the tissue and then aspirated after 15s. the recovery solution is finally pooled in a low binding tube. Three extractions steps were performed per region using different solution: (1) 0.1% TFA, (2) ACN/0.1% TFA (8:2, v/v), and (3) MeOH: 0.1% TFA (7:3, v/v). Two extraction cycles per point were performed to increase the amount of material collected.

##### NanoLC-MS & MS/MS analysis

Prior to MS analysis, the reconstituted samples were desalted using C-18 Ziptip (Millipore, Saint-Quentin-en-Yvelines, France), eluted with 80% ACN and vacuum-dried. The dried samples were resuspended in 0.1% FA aqueous/ACN (98:2, v/v). Peptides separation was performed by reverse phase chromatography, using a NanoAcquity UPLC system (Waters) coupled to a Q-Exactive Orbitrap mass spectrometer (Thermo Scientific) via a nano electrospray source. A pre-concentration column (nanoAcquity Symmetry C18, 5 μm, 180 μm x 20 mm) and an analytical column (nanoAcquity BEH C18, 1.7 μm, 75 μm x 250 mm) were used. A 2 h linear gradient of acetonitrile in 0.1% formic acid (5%-35%) was applied, at the flow rate of 300 nl/min. For MS and MS/MS analysis, a data dependent mode was defined to analyze the 10 most intense ions of MS analysis (Top 10). The MS analysis was performed with an m/z mass range between 300 to 1600, a resolution of 70,000 FWHM, an AGC of 3e6 ions and a maximum injection time of 120 ms. The MS/MS analysis was performed with an m/z mass range between 200 to 2000, an AGC of 5e4 ions, a maximum injection time of 60 ms and the resolution was set at 17,500 FWHM. To avoid any batch effect during the analysis, the extractions were chosen at random to create analysis sequences.

##### Data analysis

All MS data were searched with MaxQuant software (40,41) (Version 1.5.3.30) using Andromeda search engine (42) against the complete proteome for *Homo sapiens* (UniProt, release July 2018, 20 412 entries).Trypsin was selected as enzyme and two missed cleavages were allowed, with N-terminal acetylation and methionine oxidation as variable modifications. The mass accuracies were set to 6 ppm and 20 ppm respectively for MS and MS/MS spectra. False discovery rate (FDR) at the peptide spectrum matches (PSM) and protein levels was estimated using a decoy version of the previously defined databases (reverse construction) and set to 1%. A minimum of 2 peptides with at least one unique is necessary to complete the identification of a protein. The MaxLFQ algorithm (43) was used to performed label-free quantification of the proteins. The resulting file was analyzed using Perseus software (version 1.6.0.7). First, hits from the reverse database, proteins with only modified peptides and potential contaminants were removed. Statistical analyses were performed using ANOVA with a truncation value based on “Benjamini Hocheberg FDR” of 5%. Three categorical annotation groups were used for the ANOVA, i.e. (1) the color group based on the three colors from Scils global segmentation of the 46 samples (Red; Yellow and Blue), (2) the patient groups which are determined by the main color present in each tumor sample (Groups A, B, C) and (3) the patients survival time (patients with an OS > to the third quartile, patients with an OS between the first and the third quartile and patients with an OS < to the first quartile). Proteins significantly different were selected and normalized by a Z-score with matrix access by rows. For representation, a hierarchical clustering was performed using the Euclidean parameter for the distance calculation, and the average option for linkage in the rows and columns of the trees with a maximum of 300 clusters.

### System biology analyses

An annotation analysis of gene ontology terms for the identified proteins were performed using PANTHER Classification System (version 14.1, http://www.pantherdb.org), FunRich (Version 3.1.3) (44) and the STRING database (version 11.0, www.string-db.org) (45). Potential interaction network was then loaded into Cytoscape 3.7.2 with relative expression data using Idmapper (46).The Reactome FI plugging was used to select a subnetwork of gene ontology terms and NCI database-associated disease-specific proteins.

The relationships between the differentially expressed proteins among all conditions were also depicted based on the Ariadne ResNet database (47) using Elseviers’ Pathway Studio (version 11.0, Elsevier). The subnetwork Enrichment Analysis (SNEA) algorithm was used to detect the statistically significant altered biological pathways in which the identified proteins are involved.

### Human Pathology Atlas

The glioma data contained in the Human pathology atlas (48) were used. Based on TCGA transcriptomics and antibody-based protein data from 153 patients, this database identified 268 potentially prognostic genes (201 unfavorable and 67 favorable prognoses). These data were compared to the proteins identified in our study.

### Alternative Proteins identification

RAW data obtained by nanoLC-MS/MS analysis were analyzed using Proteome Discoverer V2.3 (Thermo Scientific) LFQ quantification with the following parameters: trypsin as enzyme, 2 missed cleavages, methionine oxidation as variable modification and carbamidomethylation of cysteines as static modification, Precursor Mass Tolerance: 10 ppm and fragment mass tolerance: 0.6 Da. The validation was performed using Percolator with a FDR set to 0.001%. A consensus workflow was then applied for the statistical arrangement, using the high confidence protein identification. The protein database was uploaded from Openprot (https://openprot.org/) and included RefProt, novel isoforms and AltProts predicted from both Ensembl and RefSeq annotations (GRCh38.83, GRCh38.p7) (49–52) for a total of 658263 entries. The identified abundance was extracted to PD2.3 and loaded in Perseus to performed statistical analysis and graphical representation.

### Statistical analyses

First, descriptive analyses were performed on clinical data. Patients were divided into 3 groups according to the quartiles of the overall survival (<Q1, Q1-Q3, > Q3). The Cox model was used to determine which proteins were most associated with overall survival. Stepwise analysis and bootstrap methods (500 samples) were used to guarantee the robustness of the results. The proteins selected after this step were used to carry out a hierarchical classification (Euclidean distance and Ward’s method) on the 46 patients to determine if there were any subgroups (clusters). Finally, the clinical variables were analyzed according to the different clusters in order to provide a clinical description of the clusters obtained. Statistical analyses were performed using the SAS Software, V9.4.

### Confirmatory immunohistochemistry analyses

Survival group validation was performed using antibodies directed against ALCAM, RPS14, ANXA11, PPP1R12A, MAOB, IGHM, HSPD1 and LASP1. The tissues were incubated with a primary antibody at 4°C overnight, followed by application of a secondary antibody (alexa fluor conjugated antibody, 1/1 000 dilution) for 1 hour at RT. We used the following primary antibodies: ALCAM (R&D Systems; 1/40 dilution), RPS14 (Invitrogen, 1/100 dilution), ANXA11 (OriGene, 1/100 dilution), PPP1R12A (Invitrogen, 1/250 dilution), MAOB (Abbexa; 1/100 dilution), IGHM (Abcam, 1/50 dilution), HSPD1 (Abcam, 1/200 dilution) and LASP1 (Santa Cruz, 1/50 dilution). All slides were imaged on the Zeiss LSM700 confocal microscope. Three to four pictures were taken for each tumor section. Processing of the images and fluorescence intensity quantification was performed using ImageJ software.

## Results

### Clinical characteristics

Forty-six IDH WT-glioblastoma samples from a prospective cohort were collected (**Supp. Table 2, Supp. Figure 1A**). Thirty-one patients (67%) were male, the median age at diagnosis was 60 (interquartile range (IQR), 51-66), the median Karnofsky performance score at diagnosis was 90 (80–90). Twenty-six (57%) patients had a complete resection. A methylation of MGMT promoter was noted in 15 tumors (33%), an EGFR amplification was noted in 24 cases (52%) and a homozygous CDKN2A deletion in 34 cases (74%). A standard treatment was initiated in 42 patients (91%). At the time of the analysis, 38 patients (83%) had a progression with a median PFS of 11 months. After a median follow-up of 19.4 months (IQR 13.5-32), 39 (85%) had died. The median overall survival was 19.4 months. The pathologist (CAM) defined regions of interest for each tumor sample: tumor, necrosis, and endothelial proliferation, after hematoxylin eosin safran (HES) staining **(Supp. Figure 1B**).

### MALDI MSI allows patient grouping based on molecular features

Considering the heterogeneity of GBM, we conducted spatially resolved proteomic studies guided by mass spectrometry imaging (MSI) (**Figure 1A**). A comparison between the pathologist annotations and the MSI molecular images showed discrepancies for many samples (**Figure 1B**, **Supp. Data 1**). A global clustering was then performed by subjecting spectra from all tissue samples to spatial segmentation (**Figure 1C**). Three main regions were identified *i.e.* red, blue and yellow areas according to the segmentation map (**Figure 1C, Supp. Figure 1C**). We then assigned each tumor sample to a group. If a colored area was greater than 50% of the surface area, the tumor was included into the group associated to this color (**Figure 1D, Supp. Figure 1C**). Therefore, 11 samples mainly represented by the red region (86,5% of the tissue surface area) were assigned to group A. Nine samples mostly represented by the yellow region (69,9% of the tissue surface area) were assigned to group B and 22 samples mostly represented by the blue region (74,9% of the tissue surface area) to group C. The 4 remaining tumors could not be classified due to the presence of many different molecular regions (**Supp. Figure 1C**). These tumors were excluded from further proteomic analyses. Within a group, all patients shared common molecular characteristics, meaning that the spectra in each of these colored regions were similar. Some specific ions can be attributed to each region: m/z 967,621 and 1492,916 were specifically present in group A, m/z 1914,591, 2375,074 and 2376,274 were specific to group B and m/z 1473,312, 2045,815, 2046,615 and 2237,849 were specific to group C. Images of some group-specific ions are shown in **Figure S1D.** The Ward clustering method using IMAGEREVEAL MS Ver.1.1 software confirmed the segmentation of the 46 tumors into 3 groups with similar specific ions (**Supp. Figure 1E**). We also confirmed, with this method, the heterogeneity of the 4 outliers.

**Figure 1:**
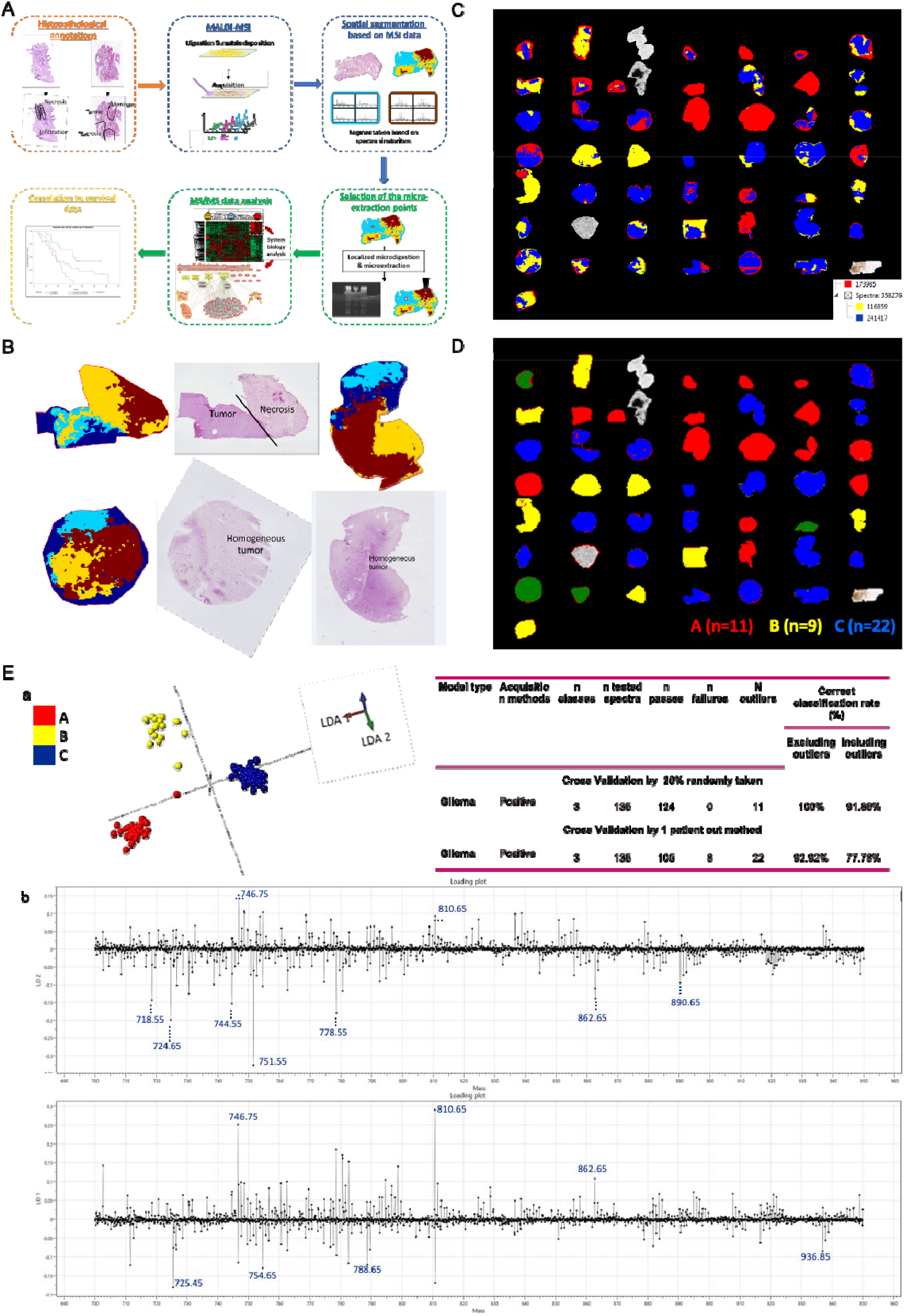
Histological, MALDI MSI and SpiderMass analysis. A. General workflow of the MALDI-MS imaging combined with microproteomics used for glioma classification. B. Representative annotated histopathology images of three glioblastoma samples and their corresponding segmentation map obtained from MALDI-MSI data. Colors represent molecularly different regions. Note that for 2 different tissues, similar colors are not equivalent to similar molecular groups. The segmentation map shows different clusters for each case and non-observable with HES coloration. C. Global segmentation maps of all tissues together after MALDI-MSI analysis. Colors represent molecularly different regions as shown in the corresponding dendrogram. The segmentation map gives 3 main clusters. D. Cases are grouped together based on their molecular profiles: group A (n=13) contains cases predominantly red, group B (n=9) contains cases predominantly yellow and group C (n=23) contains cases predominantly blue E. The built PCA-LDA classification model based on 3 glioma groups; Group A (red), Group B (yellow), Group C (blue). a) LDA representation of the 3-class PCA-LDA (right). The table (right) represents the “20% out” and “leave-one-patient-out” cross-validation results of the built classification model. b) LD2 loading spectra (top) indicate the discrimination between Group A (red) and Group B (yellow). The 10 most discriminatory lipid peaks are indicated by the blue dash line. LD1 loading spectra (bottom) indicate the discrimination between Group A (red) and Group C (blue). The 10 most discriminatory lipid peaks are indicated by the blue dash line.

In order to validate the classification obtained by MALDI MSI, we analysed 30 samples by SpiderMass technology (36,38). Following the acquisition of the MS spectra in positive ion mode, a PCA analysis of the generated spectra acquired from 30 tumor tissues was performed. The features of the PCA were subjected to a supervised analysis using linear discriminant analysis (LDA) (53,54) which yielded 3 groups (**Figure 1Ea**). According to **Figure 1Ea**, LDA 1 discriminated group A from group C and the LDA 2 separated group B from group A and C. The LDA analysis of the SpiderMass data therefore allowed the samples to be grouped in the same way as the MALDI-MSI classification. Some examples of discriminant ions (*m/z*) between the three groups, corresponding to lipids, are presented as their normalized intensities in **Figure 1Eb**. The most discriminating peaks for group A in LD2+ correspond to *m/z* 746.75, and 810.65; for group B in LD2-correspond to *m/z* 718.55, 724.65, 744.55, 751.55, 778.55, 862.65 and 890.65; for group C in LD1-correspond to *m/z* 725.4, 754.6, 788.65, and 936.85. To consolidate the classification, validation was performed using either 20% randomly patients taken out or the one-patient-out method (**Inset table in Figure 1E**). Excellent cross validation results were obtained using 20% randomly patient taken out method with 100% and 91.85% correct classification rates with and without outliers respectively and good classification using the one-patient-out method with 92.92% and 77.78% including or not outliers respectively (**Inset table in Figure 1E**). These results of outliers and misclassifications (mainly group B) reflect the fact that each group is not represented by only one colored region. We then correlated our classification to the clinical data (**Supp. Table 3, Supp. Figure 2**).

### Identification of specific signaling pathway signatures for each group

In order to understand the molecular differences between the three groups, spatially-resolved tissue proteomic was undertaken on the 46 tissue samples (55). On each tissue, 2 to 5 specific micro extraction points were selected according to the molecular regions identified by spatial segmentation of MALDI MSI data (**Supp. Data 1**) in order to analyse the tumor heterogeneity and micro-environment presenting with specific protein signatures in each group. This resulted in a total of 122 micro-extraction points. Each extraction point was associated with one of the three regions identified by MALDI-MSI (red, yellow and blue). In all tumor samples, 25 extractions were performed in the red region, 22 in the yellow region and 73 in the blue region. From these shotgun proteomic experiments, a total of 4875 proteins were identified (**Supp. Data 2**). Some proteins were unique to a patient, *i.e*. 354 unique proteins distributed among the patients (**Supp. Data 3**). From these proteins, several transcription factors (CD2 antigen cytoplasmic tail-binding protein 2), enzymes (MMP12, AKT1) but also some proteins related to the immune response (HLA-A3, HLA-B07), extracellular vesicles initiating factors (RAL-A1), tumor suppressors or specific tumor antigens like Melanoma antigen were identified.

To better understand the differences between each identified group, ANOVA tests with a Benjamini Hochberg FDR of 0.05 was performed. A total of 1185 proteins showed a significant difference in expression between the three groups (**Figure 2A, Supp. Data 4**). Two main branches were identified in the heatmap. The first branch was composed of 100% of samples extracted from the yellow region and correspond to the group B. The second branch separatess group A from group C. This branch is then separated into two sub-branches corresponding to group A and regrouping 78,6% of the samples extracted from the red region and 21,4% of the samples extracted from the blue region. The second sub-branch corresponds to the largest cluster, group C and contains 76,1% of the samples extracted from the blue region, 12,7% of the samples extracted from the red region and 11,3% of the samples extracted from the yellow region. These 3 regions are consistent with the regions identified by MALDI MSI spatial segmentation. We confirmed that each sample from the same colored region has the same proteomic profile (**Supp. Data 4**). Three specific clusters of over-expressed proteins for each group were identified using the heatmap (**Figure 2A**) *i.e.* cluster 1 corresponds to proteins overexpressed in group B; cluster 2, to proteins overexpressed in group A and cluster 3, to proteins overexpressed in group C. The lists of overexpressed proteins per group are presented in **Supplementary Data 4**.

**Figure 2.**
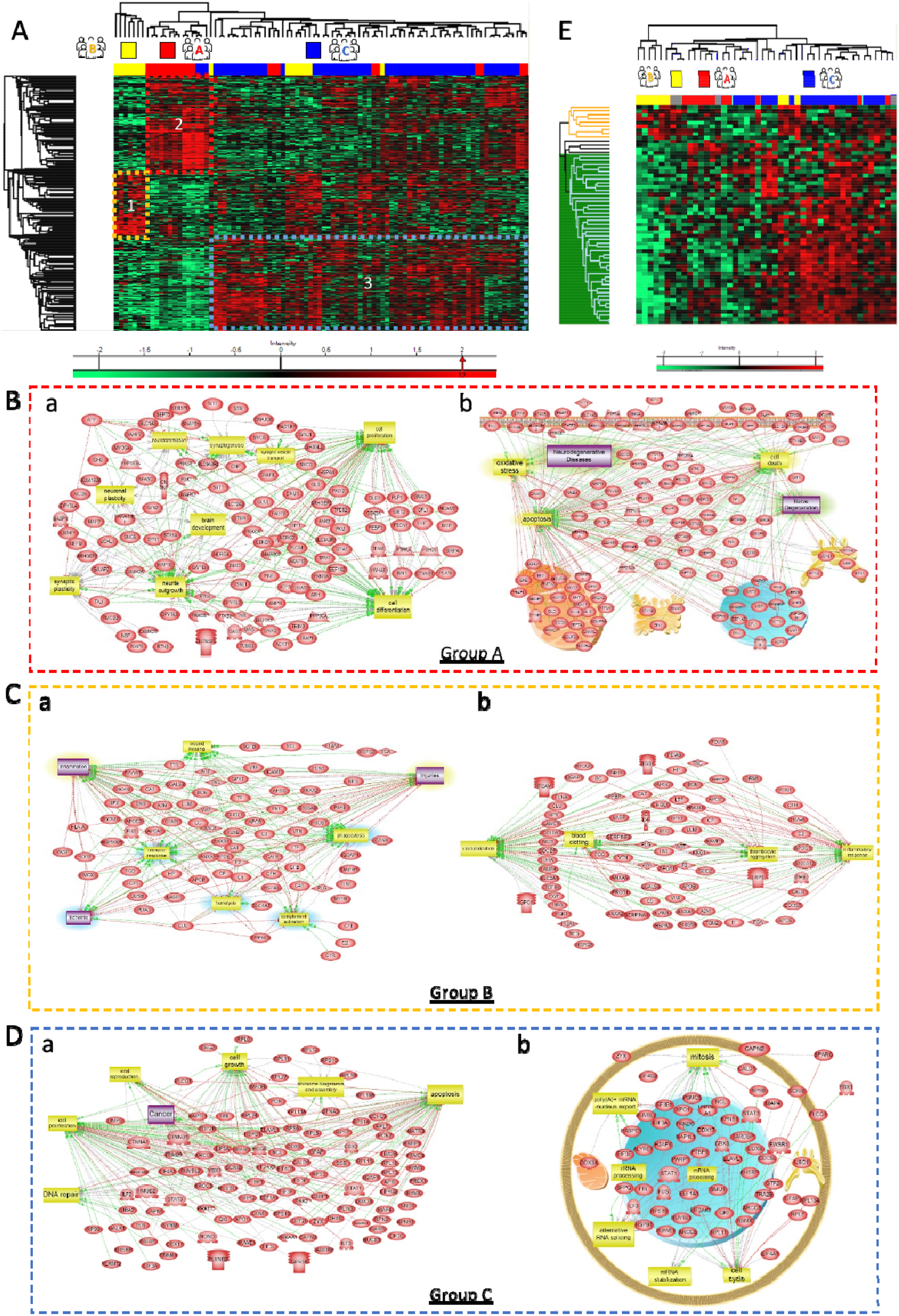
Shotgun microproteomics analysis. A. Heatmap of proteins with different regulation profiles as determined after label free quantification in the three groups highlighting the presence of 3 clusters. Shotgun proteomics was performed after on-tissue trypsin digestion followed by microextraction at the spots determined from MALDI MSI data. B. Pathway analysis of proteins overexpressed in group A reveals that a large majority of protein is involved in (a) neurogenesis, brain development, axon development and (b) more generally neurodegenerative disease and nerve degeneration. C. Pathway analysis of proteins overexpressed in group B reveals that majority of protein are involved in (a) injuries, inflammation and more generally immune system response but also in (b) blood coating and vascularization. D. Pathway analysis of proteins overexpressed in group C shows implication in (a) cancer, cell growth, DNA repair and viral reproduction with protein involved in (b) RNA splicing, metabolism and replication. E. Heatmap of alternatives proteins with different regulation profiles as determined after label free quantification in the three groups highlighting the presence of 3 clusters.

In group A (mainly represented in cluster 2), the proteins are associated with neuro-developmental genes, that are characteristic of neuronal/glial lineages or progenitor cells. Most proteins were related to neurogenesis and axon guidance (Dihydropyrimidinase-related protein 1 (CRMP1), Misshapen-like kinase 1 (MINK1), Neuromodulin (GAP43), Dihydropyrimidinase-related protein 5 (DPYSL5), Dihydropyrimidinase-related protein 4 (DPYSL4), Microtubule-associated protein tau (MAPT), Kinesin-like protein KIF2A (KIF2A), Neurofilament heavy polypeptide (NEFH), Unconventional myosin-XVIIIa (MYO18A), MAGUK p55 subfamily member 2 (MPP2), Alpha-internexin (INA), CLIP-associating protein 2 (CLASP2) (**Supp. Data 4**). Using the functional enrichments analysis tool of String database, the most representative Reactome pathway was devoted to axon guidance. Nine of the 16 proteins identified in this pathway are involved in neuron development projection, morphogenesis, and guidance (**Supp. Figure 3Aa**). System biology analyses using SNEA and Cytoscape confirmed that the proteins in group A (Cluster 2) are involved in neurite outgrowth, synaptogenesis, synaptic plasticity (**Figures 2Ba & 2Bb, Supp. Data 5**). Interestingly, among the identified proteins some are known to be involved in tumorigenesis like Mitogen-activated protein kinase 3 (MAPK3), Protein kinase C alpha type (PRKCA) and some were already identified in glioblastoma *e.g.* CRMP1, DPYSL2 (i.e. CRMP2), (56), DPYSL5 (i.e. CRMP5) (57), GAP43 (58,59,60), as well as Tau protein encoded by MAPT in low-grade glioma (61).

**Figure 3.**
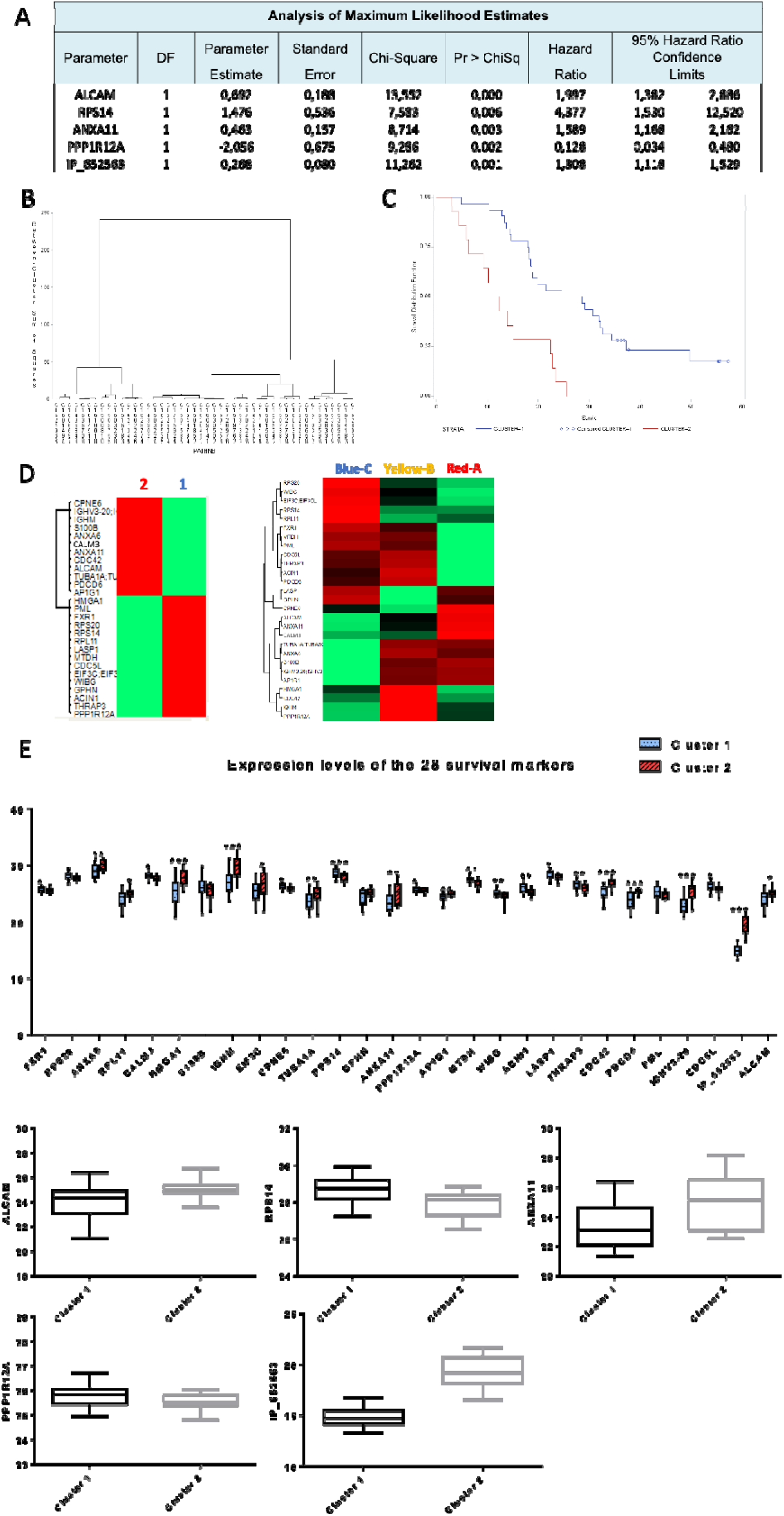
Proteomic and lifetime analysis. A. Analysis of maximum likelihood estimates of the 5 proteins significantly correlated with survival (ANXA11, RPS14, ALCAM, PPP1R12A and AltProt IP_652563) identified after a step by step analysis and boostrap procedure and B. patient clustering based on these proteins C. Overall survival of the 46 patients according to the expression of the 5 prognostic markers. Two clusters of patients were identified with a clear difference in their survival. Cluster 1 has the longer survival while cluster 2 has the shorter survival. D. Heatmap of the 28 proteins significant after Cox model (p=0.01) between the 2 groups of patients defined by their OS (left) and expression of these 28 proteins in each molecular group A, B and C (right) E. Boxplots of the 28 prognosis proteins significant after applying the Cox model. Their LFQ values were compared between patients of cluster 1 (long survival) and cluster 2 (short survival). F. Boxplots of the 5 prognostic markers identified after a step by step analysis and bootstrap procedure. Their LFQ values were compared between patients of cluster 1 (long survival) and cluster 2 (short survival).

Proteins overexpressed in group B (mainly represented in cluster 1) were linked to microglia activation and more generally immune system activation. Indeed, among the proteins identified, 10 proteins are linked to the immune response such as complement C1q subcomponent subunit C and B (C1QC and C1QB), complement factor H (CFH), haptoglobin (HP), kininogen 1 (KNG1), histidine-rich glycoprotein (HRG), transthyretin (TTR), grancalcin (GCA), proteins S100-A9 (S100A9) & S100-A12 (S100A12), erythrocyte band 7 integral membrane protein (STOM) and galectin-3-binding protein (LGALS3). Immunoglobulin heavy and light chains (IGHG2; IGKC; IGHG1; IGLC6; IGHM and IGHA1) and macrophage markers, Macrophage-capping protein (CAPG) were also detected (**Supp. Data 4**). Moreover, some proteins are related to iron transporters like ceruloplasmin (CP), serotransferrin (TF), hemopexin (HPX) and haptoglobin, and other proteins are associated to coagulation *e.g.* Transthyretin, Kininogen-1 (KNG1), Plasminogen (PLG). Most of these proteins are known to be present in human plasma (48). These results are in accordance with histological annotations reflecting that most of the tissues belonging to group B present intense proliferation of capillary endothelial cells with inflammation and hemorrhage (**Supp. Figure 1B**). The cytoscape and SNEA analysis, (**Figure 2Ba and b, Supp. Data 5**), clearly confirmed that most of the proteins are involved in complements, coagulation cascade, inflammation, ischemia, vascularization, wood healing, and cancer. Moreover, 11% of patients of group B present more thromboembolic event (**Supp. Table 3**). The same pathways were found in Reactome (**Supp. Figure 3Ab**). Some of these proteins have already been detected in TGCA glioma database (see below) and are mostly unfavorable prognosis indicators for patient survival *e.g* Grancalcin and CAPG (**Supp. Figure 3B**).

The overexpressed proteins in the group C (mainly represented in cluster 3) are mainly involved in tumor growth (Hepatoma-derived growth factor (HDGF), Developmentally-regulated GTP-binding protein 2 (DRG2)), but also in virus infection (Eukaryotic translation initiation factor 3 subunit L (EIF3L), Double-stranded RNA-binding protein Staufen homolog 1 (STAU1) and Interferon-induced double-stranded RNA-activated protein kinase (EIF2AK2)) (**Supp. Data 4**). KEGGS analyses confirmed a network of proteins involved in Epstein barr infection (**Supp. Figure 3Ac**). Cytoscape pathway analyses established that this group is linked to viral infection and antiviral immune response (**Figure 2Da**). System biology analyses confirmed the involvement of proteins in virus infection (transfection, reproduction) and transcriptomic modification at the RNA level (RNA splicing, metabolism, replication) (**Figure 2Db, Supp. Data 5**). Some other markers of the group C are known to be bad prognosis indicators such as EIF2AK2 and ZC3HAV1.

### Identification of alternative proteins

Using OpenProt alternative proteins database (Brunet et al., 2018), 257 AltProts were identified and 170 were quantified. After ANOVA tests with a p-value of 0.05, 58 were differentially expressed between groups (**Figure 2E**). In group A, four AltProts are over-expressed coming from ncRNA, IP_2390879 issued from LOC107985743, IP_244732 from KIFC3, involved in cell adhesion, IP_672223 from GBP1P1 and IP_710015 from LRRC37A9P (**Supp. Table 4**). In group B, we found a cluster of nine over-expressed AltProts. Five are transcribed from ncRNA, two are located at the 5’UTR of mRNA, one at the 3’UTR and one result from the frame shift in the CDS **(Supp. Table 4**). In group C, 45 AltProts are over-expressed: 24 from ncRNA, six from the 5’UTR, 10 from the 3’UTR and five result from the frame shift in the CDS (**Supp. Table 4**). Taken together, we identified several AltProts issued from ncRNA (~57%) which is in line with our previous work on glioma cell line (NCH82) (Cardon et al., 2020b). Like our previous work, we also identified height AltProts in the present study, especially in the group C such as IP_2323408 coming from LOC105376126 or IP_261897 from LOC100506371 the uncharacterized gene and currently described as a ncRNA: IP_755940, IP_593099, IP_774693, IP_572422 et IP_671464. These last five AtlProts are pseudogenes for: HNRNPA1P30, TUBB2BP1, TUBAP2, TUBBP1, and TPI1P1 respectively. These pseudogenes, for which no protein has been observed yet, have an expression of their transcripts in glioma cell lines (Expression Atlas) (Petryszak et al., 2016). Interestingly, the last one IP_079312, from the mRNA encoding EDARADD was correlated with a low survival rate in ovarian cancer patients (Cardon et al., 2020b). In another study, we found interaction partners of AltProts by XL-MS in NCH82 cells in physiological condition and during a phenotypic change (Cardon et al., 2020a). Five significantly variable AltProts were found in this study, one of them is also overexpressed in group B: IP_156671 which originates from the 3’UTR of the transcript coding for SLC13A1. The four others are overexpressed in group C: IP_261897 coming from a ncRNA, IP_063564, IP_256988 both issued from the 3’UTR region of the CLDN19 and TBX21 genes respectively and IP_073718 originate from a shift in the reading frame of the CCDC181 gene.

### Correlation between TCGA and proteomic data

We then compared our almost 5000 identified proteins to the TCGA database, on which 682 genes show an elevated expression in glioma; 282 of these 682 genes were found in our samples (**Supp. Table 5**). In these 682 genes, 268 genes are suggested as prognostic based on transcriptomics data from 153 patients; 201 genes associated with an unfavourable prognosis and 67 genes associated with a favourable prognosis. In our proteomics data, we found 12 proteins associated with an unfavourable prognosis: 7 proteins are over-expressed in group A (CEND1, DMTN, PAK1, MAP2K1, THY1, VSNL1 and FN3KRP), 2 proteins are over-expressed in group B (AEBP1 and PDIA4) and 3 protein are over-expressed in group C (POR, ERLIN2 and DBNL) (**Table 1**). We also found 9 proteins associated with a favourable prognosis: 7 proteins are over-expressed in group A (GLUD1, GDI2, SARS, SEPT2, PHGDH, KPNA3 and ARHGEF7), and 2 protein are overexpressed in group C (PABPC1 and RBBP4) (**Table 2**). In all groups we found prognosis proteins. We also established that among the proteins identified in the 3 different groups, those in group A are closely correlated with the TCGA glioma data. GBM hijacks mechanisms of neural development and contains subsets of glioblastoma stem cells (GSCs) which are cells with tumor-propagating potential. This cell state is mainly represented in tumors of group A. In fact, group A presents an overexpression of Synaptotagmin. 45% of patients of group A had EGFR amplification. The median survival was between 22.3 months (Table 2). Group B has the highest level of methylated MGMT promoter (56%) and EGFR amplification were noted in 44%. The survival median was 23 months (Table 2). For Group C, 64% of patients have an EGFR amplification. The survival median was 19.4 months (Table 2).

**Table 1:**
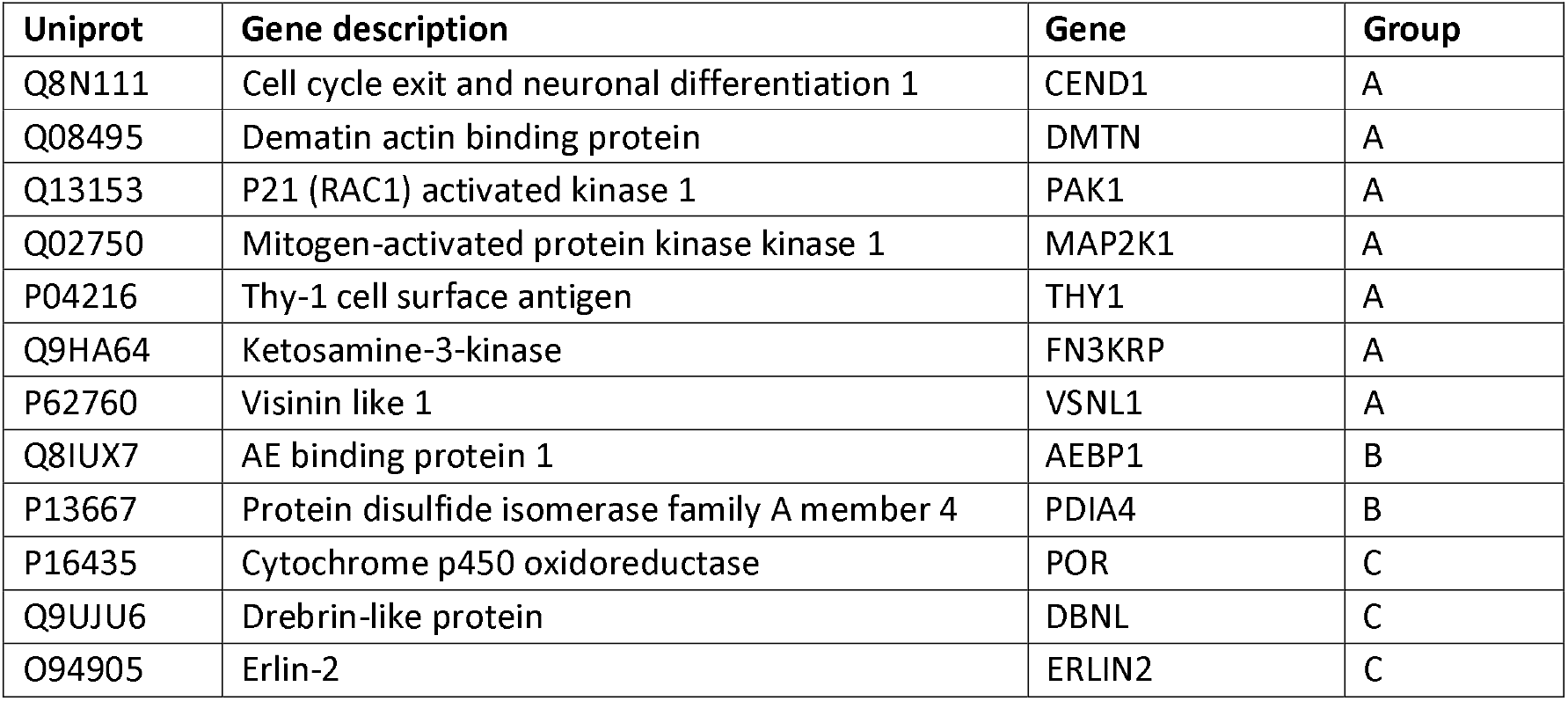
Proteins associated with unfavorable prognostic in glioma and identified in groups A, B and C.

**Table 2:** Proteins associated with favorable prognostic in glioma and identified in groups A, B and C.

| Uniprot | Gene description | Gene | Group |
| --- | --- | --- | --- |
| P00367 | Glutamate dehydrogenase 1 | GLUD1 | A |
| Q14155 | Rho guanine nucleotide exchange factor 7 | ARHGEF7 | A |
| P50395 | GDP dissociation inhibitor 2 | GDI2 | A |
| P49591 | Seryl-tRNA synthetase | SARS | A |
| Q15019 | Septin-2 | SEPT2 | A |
| O43175 | D-3-phosphoglycerate dehydrogenase | PHGDH | A |
| O00505 | Importin subunit alpha-4 | KPNA3 | A |
| P11940 | Poly(A) binding protein cytoplasmic 1 | PABPC1 | C |

### Clinical data integrated to OMICs Data

The statistical analysis of the overall survival is correlated with MGMT status (**Supp. Figure 2c**) and KPS (**Supp. Figure 2b**) but not with the extent of resection (**Supp. Figure 2d**). In order to find new prognostic proteins from our proteomic data, we performed an ANOVA test on the entire proteomic dataset (n=46) according to patients’ OS. The cohort has been divided into 3 groups: 11 patients (25%) with OS > to the third quartile, 23 patients (50%) with an OS between the first and the third quartiles and 12 patients (25%) with an OS < to the first quartile were included in this analysis. 114 proteins and 10 AltProt have shown significance between these 3 groups of patients defined by their OS (**Supp. Data 6**). Then, using Cox model, 28 proteins were significant with a p=0.01 (**Table 3, Figure 3E**). After a step by step analysis and bootstrap procedure, 5 proteins remained highly significantly correlated with survival: ALCAM, RPS14, ANXA11, PPP1R12A and the AltProt IP_652563 (**Figure 3A, 3F**). Based on the expression of these 5 proteins, 2 clusters of patients were identified (respectively cluster 1 and cluster 2) (**Figure 3B**). The OS of the patients from the 2 clusters was determined (**Figure 3D**). Patients of cluster 2 (n=14) have the worst OS, whereas patients of cluster 1 (n=32) have the highest (**Figure 3C**). Among the 5 proteins significantly correlated with survival, IP_652563 is an AltProt issued from ncRNA (**Suppl. Table 6**). This ncRNA is transcribed from the ENSG00000206028 gene which is expressed in glioma cell lines (ref Expression Atlas). This AltProt is a bad prognosis indicator as well as ALCAM and ANXA11 whereas PRR1R12A and RPS14 are good prognosis markers. We confirmed the expression of these 5 markers in the two clusters of patients (**Figure 3F**) based on the LFQ proteomic values. We further validated the expression of 4 of the 5 prognosis markers (ALCAM, RPS14, ANXA11 and PPP1R12A) by immunohistochemistry in the two clusters of patients (**Figure 4A**). For the AltProt, we could not do such validation since no antibody exists. We confirmed in patients from cluster 2 (worst OS), a higher expression of ANXA11, which correlates well with the proteomic data (**Figure 4B**). ALCAM was found to be higher expressed by proteomics in cluster 2. Although not significant, a slight increase of fluorescence is observed in tumors of cluster 2 as well. The expression of ALCAM is associated to blood vessels as shown on **Figure 4A** and is known to participate to immune cells infiltration. Even though no difference in fluorescence is measured, blood vessels seem to show different morphologies between patients of cluster 1 and 2 as shown on **Figure 4A.** RPS14 and PPP1R12A are higher expressed in the tumors of cluster 1 (longer OS), which was also found by proteomics. These five prognostic markers were the most significant ones but in order to validate and increase the power of our classification, we also included other markers such as IGHM and LASP1 which are included in the list of the 51 more significant prognostic markers as shown on **Figure 3E** and **Supp. Data 7**. We were also interested in HSPD1 and MAOB expression in glioblastoma since these two proteins are known to be involved in several cancers including glioma. The expression of HSPD1 and LASP1 is associated to a higher survival while the expression of MAOB and IGHM is associated to a lower survival. As shown on **Figure 4C**, HSPD1 and LASP1 were found higher expressed in tumors of cluster 1 whereas MAOB and IGHM are higher expressed in tumors of cluster 2. Therefore, we have identified a panel of markers associated to higher survival (PPP1R12A, RPS14, HSPD1 and LASP1) and another panel associated to lower survival (ALCAM, ANXA11, MAOB and IGHM).

**Table 3.**
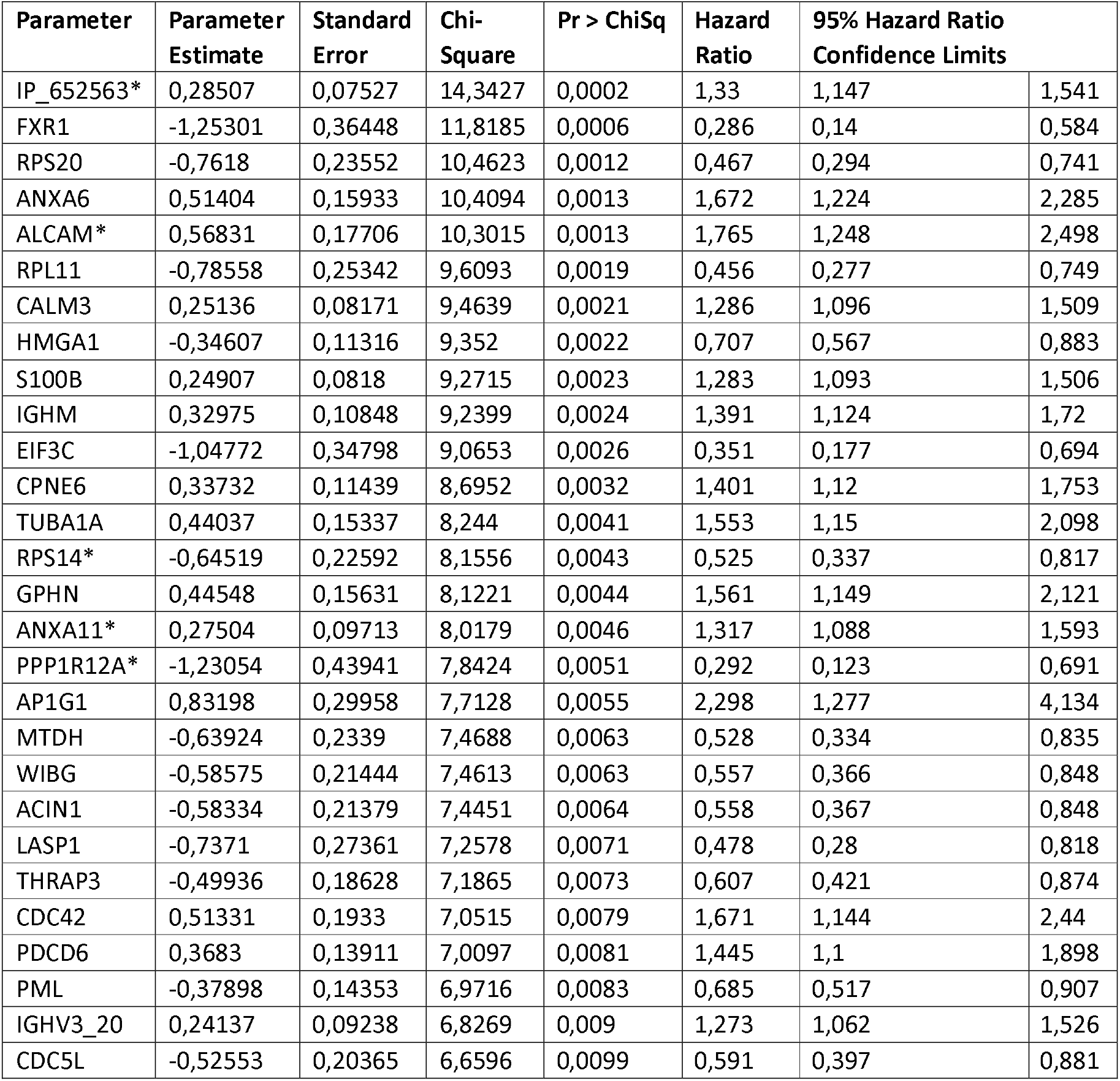
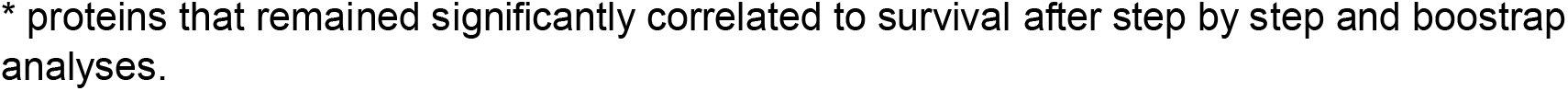
Proteins associated with survival after Cox model p = 0.05 ^*^proteins that remained significantly correlated to survival after step by step and boostrap analyses.

**Figure 4.**
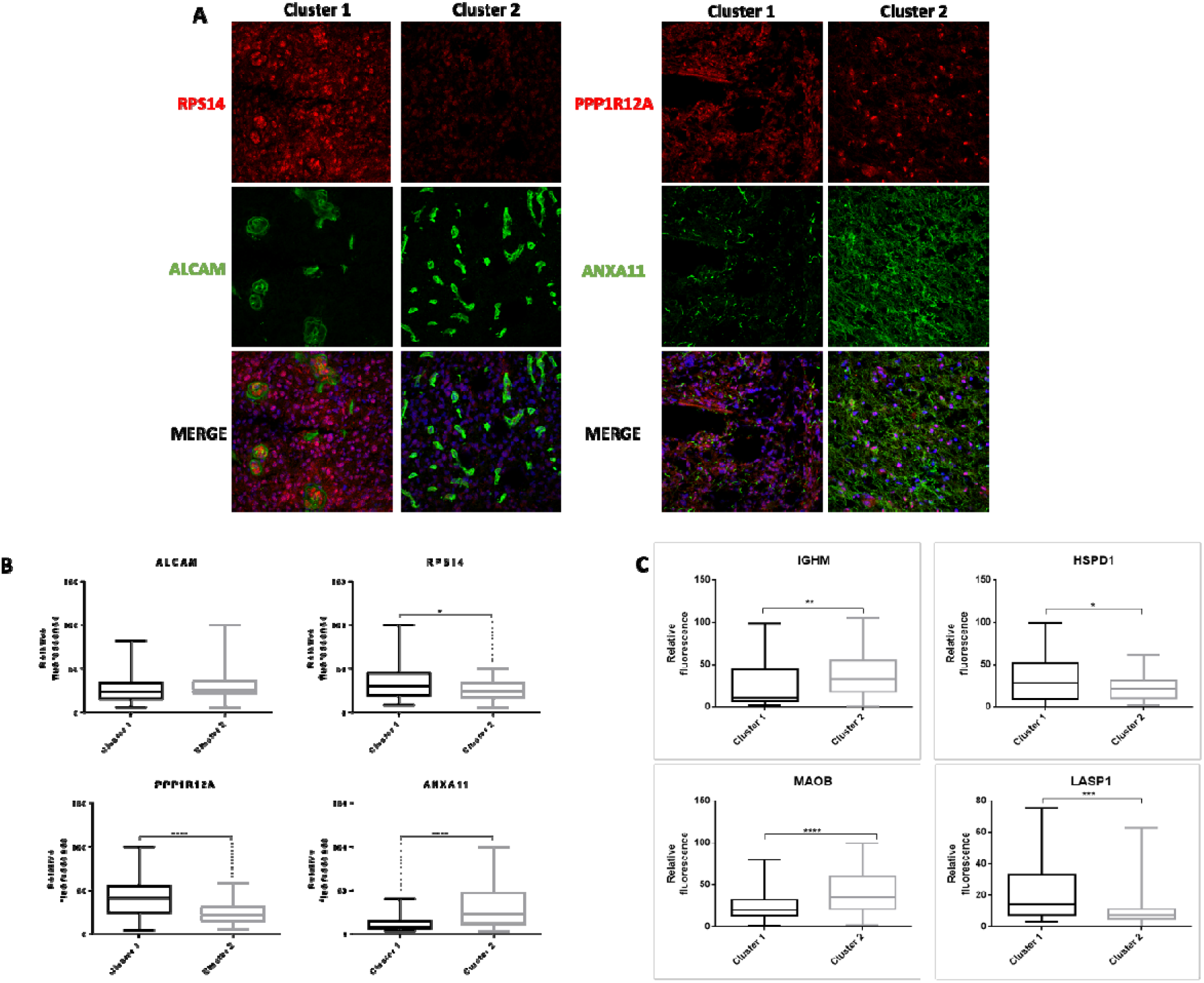
Validation immunohistochemistry of the panel of survival markers identified. A. Representative fluorescence images of the 4 proteins in the two OS clusters of patients. ANXA11 and ALCAM are associated to a bad prognosis while PPP1R12A and RPS14 are related to a good prognosis. Images were acquired with a confocal microscope at 40x magnification. B. Quantification of fluorescence intensities of the 4 proteins in the two OS clusters. Images taken from 14 tumors of cluster 1 and 9 tumors of cluster 2 were quantified. For each tumor, 3 to 4 images were acquired and quantified. Significant differences were identified using unpaired t test with **** p<0.0001; *** p<0.001; ** p<0,01 and * p<0.05. C. Quantification of fluorescence intensities of 4 additional prognostic proteins identified after the Cox model procedure: HSPD1, LASP1 whose expression is associated to a longer survival and MAOB, IGHM whose expression is associated to a shorter survival. Images taken from the 46 tumors were quantified. For each tumor, 3 to 4 images were acquired and quantified. Significant differences were identified using unpaired t test with **** p<0.0001; *** p<0.001; ** p<0,01 and * p<0.05.

Then, in order to know, if these prognostic markers are related to one of the molecular groups A, B or C, we compared the expression of the 28 prognostic markers identified between the 2 OS clusters and the 3 molecular groups identified by MALDI-MSI (**Figure 3D**). We found that group C is more related to cluster of survival 1 with the co-expression of 12 markers including RPS14 and LASP1 associated to a longer survival. Group A shares the expression of 9 markers with cluster 2, including ALCAM, ANXA11 associated to a shorter survival. Group B has both markers of long and short survival such as PPP1R12A and IGHM/ respectively. Therefore, group C express more good prognosis markers while group A is more related to bad prognosis based on the expression of these markers. Taken together, we established a new classification of glioblastoma based on molecular and cellular features which can be very useful for the development of new therapeutic options and personalized medicine for patients.

## Discussion

Here, we investigated glioblastoma biology by a proteomic approach at a low spatial resolution in order to capture the tumor microenvironment. A non-targeted MALDI-MSI analysis followed by spatial segmentation using different algorithms allowed to derive a new subclassification of glioblastoma. We successfully validated these observations with SpiderMass technology with a 93% good classification. Three subgroups were identified (A-Red, B-Yellow and C-Blue regions). In order to decode the biological pathways involved in these three groups, we performed a spatially resolved proteomic that confirmed the data. Molecular signatures of different tumor subtypes were identified among the groups.

Group A is associated with neuro-developmental genes, characteristic of neuronal/glial lineages or neural progenitor cells (NPC) (**Figure 2B**). These included nervous system development markers (like CRMP family, GAP43, MAPT), oligodendrocyte development and differentiation markers (like ABI1, ASPA, CNP, CNTNAP1), stem and progenitor cell signatures (like TRIM2). However, NPC-like state is correlated with presence of markers for immature neurons (beta-3-tubulin), markers for mature neurons (NeuN) and markers indicative for synapses (synaptophysin, SV2A) (62). In our data, we found Stathmin 1, NEFH, NEFM and NEFL (63) which are also markers of the NPC-like state of the GSC. Patients of group B present the lowest OS and proteins enriched in this group are linked to immune status with macrophages infiltration, (**Figure 2C**) such as complement factors, immunoglobulin heavy and light chains (IGHG2; IGKC; IGHM; IGHG1; IGLC6 and IGHA1), macrophage markers (CAPG) and coagulation cascade proteins (HP, KNG1, HRG, TTR, GCA, S100A9, STOM). These results are in line with a study of (64), in which eight immune related genes (FOXO3, IL6, IL10, ZBTB16, CCL18, AIMP1, FCGR2B, and MMP9) were identified and used as unfavorable prognostic markers in GBM. High-risk patients conferred an enhanced intensity of local immune response compared to low-risk ones. From the 8 signature genes, AIMP2 has been identified in group B also. Patients of group B also express GSC markers but with a “stem-to-invasion” path. CD44, NES and VIM, enriched in group B, are markers of the mesenchymal like state.

The presence of class I self-antigen HLA proteins (HLA-A3 and HLA-B07) in group B is interesting since a positive correlation between HLA and some cancers has been demonstrated, such as cervical or nasopharyngeal carcinomas (65). In a previous study based on HLA antigen frequencies in patients with glioma compared with the control population, patients positive for HLA-A*25 had a 3.0-fold increased risk of glioma (p = 0.04), patients positive for HLA-B*27, a 2.7-fold risk (p = 0.03). In contrast, the relationship between HLA-B*07 and glioma is very rare (66), as well as for HLA-A*3 (67). Taken together, these data clearly confrimed that there is interpatient molecular heterogeneity that may be related to tumor phenotype and cellular plasticity (63) but not directly with transcriptional classification of glioblastoma (proneural, neural, classic and mesenchymal) (23). Finally, systemic biology analyses revealed that group C is linked to an anti-viral immune response and viral infection. Recent studies have reported a link between glioblastoma and perinatal viral exposure (68–71). It has recently been shown that Epsein-Barr virus can be implicated in GBM etiology (72). Moreover, some recent works have also reported that Cytomegalovirus (CMV) promotes murine glioblastoma growth via pericytes recruitment and angiogenesis (73). In human, CMV nucleic acids and proteins have been observed within GBM tumor tissue (74).

The comparison with the TGCA specific glioma genes signature showed that 28 of them were statistically different among the 3 groups identified in our study. Most of the proteins were identified in group A and are related to nervous system development, neuron differentiation axon guidance, 3 proteins were identified in group B and are linked to cytokine secretion and 5 both in groups A & C related to Notch signalling. Notch signaling is an evolutionarily conserved pathway that regulates important biological processes, such as cell proliferation, apoptosis, migration, self-renewal, and differentiation. Growing evidence reveals that Notch signaling is highly active in glioma stem cells, in which it suppresses differentiation and maintains stem-like properties, contributing to glioblastoma tumorigenesis and conventional-treatment resistance (75).

Taken together, we established that glioblastomas can be divided into 3 groups. Each group has a distinct molecular pattern, reflecting a specific molecular phenotype of the tumors. These different groups may be explained by an early differentiation due to the presence, in primary tumours, of subpopulation of cells with a distinct functional profile as well as the existence of cells with a high invasive potency. A recent study (76) proposed that glioblastoma stem cells (GSCs) acquire a high invasive activity through a mechanism called the ‘stem-to-invasion path’ and that long noncoding RNAs are one of the key factors. It has been demonstrated that these non-coding genomic regions can result to the synthesis of proteins, so called alternative proteins forming an unexplored ghost proteome with function in cancer (49). 170 alternative proteins (AltProts) were found significantly variable in the three groups identified above. If the function of these AltProts are still poorly understood, it has been shown that they can have a role in regulation of transcription and can also be present in extracellular vesicles (77). Finally, more than 50% of the AltProts identified in the present study come from the translation of ncRNAs transcribed from pseudogenes. Recently it has been demonstrating that pseudogenes can also be used as signatures for glioma prognosis. 6 pseudogenes (SP3P, ANXA2P3, PTTG3P, LPAL2, CLCA3P, and TDH) have been found to be associated with overall survival in glioma (78). Nine other pseudogenes (TP73-AS1, AC078883.3, RP11-944L7.4, HAR1B, RP4-635E18.7, HOTAIR, SAPCD1-AS1, AC104653.1, and RP5-1172N10.2.) constitute a set of prognosis markers to predict survival of patients with glioma (79). All these results provide novel insights into the biological role of pseudogenes in cancer and especially in glioma. Additionally, the novel identified AltProts translated from ncRNAs add additional information to the already known pseudogenes in glioma.

Finally, this study allowed us to highlight the presence of 5 new prognostic proteins for glioblastoma: PPP1R12A and RPS14 are favourable prognostic markers while ALCAM, ANXA11, and AltProt IP_652563 are unfavourable prognostic markers. We found that the expression of these unfavourable protein markers is higher in the tumors belonging to the cluster 2 (worst OS). ANXA11 was also found as a bad prognostic marker for glioblastoma in a recent study (80) and more expressed in the proteomic group A. ALCAM was found to be overexpressed in the proteomic group A as well. The favourable prognostic markers are more expressed in the proteomic group C and the OS cluster 1. We also validated 4 other markers in this cohort: HSPD1, LASP1, two favourable prognostic markers and MAOB, IGHM, two unfavourable prognostic markers. Moreover, 2 known unfavourable prognostic proteins, coming from the Human Pathology Atlas, are overexpressed in the group B (AEBP1 and PDIA4).

If we come back to the molecular signature of the patients of group B, we have demonstrated that the proteins are mainly involved in inflammation, phagocytosis, oxidative stress, and ischemia and are related to macrophages infiltration. The brain neoplasm cells could influence the polarization state of the tumor associated macrophages (81). In turn, innate immunity cells have a decisive role through regulation of the acquired immune response, but also through humoral cross-talking with cancer cells in the tumor microenvironment. Thus, we can assume that immunotherapeutic strategies would be much more suited for patients of group B. Further studies need to be conducted to see the correlation between the proteomic signatures we highlight and response to treatment. In this context, our study could be an interesting starting point to guide the development of new personalized therapeutic strategies and better treatment decisions.

## Supporting information

manuscript

## Abbreviations

A: astrocytoma
ACN: acetonitrile
ATRX: alpha-thalassemia/mental retardation syndrome X-linked
CDKN2A: cyclin-dependent kinase inhibitor 2A
CGH: array comparative genomic hybridization
DNA: deoxyribonucleic acid
EGFR: epidermal growth factor receptor
F: female
FDR: false discovery rate
FFPE: formalin-fixed paraffin-embedded
gCIMP: CpG island methylator phenotype
HCD: Higher energy Collision Dissociation
HES: Hematoxylin Eosin Safran
IDH: Isocitrate dehydrogenase
LC: Liquid Chromatography
H3F3A: H3 Histone, Family 3A
LESA: Liquid Extraction Surface Analysis
LFQ: Label-Free Quantification
M: male
MALDI: Matrix-Assisted Laser Desorption/Ionization
MALDI MSI: MALDI Mass Spectrometry Imaging
TOF: Time-Of-Flight
MeOH: Methanol
MGMT: O^6^-methylguanine-DNA methyltransferase
MRI: Magnetic Resonance Imaging
MSI: Mass Spectrometry Imaging
O: oligodendroglioma
PSM: peptide spectrum matches
PTEN: phosphatase and tensin homolog deleted on chromosome 10
ROI: Region of interest
RNA: Ribonucleic acid
SNEA: Subnetwork Enrichment Analysis
TERT: telomerase reverse transcriptase
TFA: Trifluoroacetic acid
TP53: tumor protein p53
WHO: World Health Organization

## Acknowledgments

This research was supported by grants from the Ministère de L’Education Nationale, de L’Enseignement Supérieur et de la Recherche, ANR (IF), SIRIC ONCOLille (MS), Grant INCa-DGOS-Inserm 6041aa (IF, MS), and INSERM, Ligue contre le Cancer (EL).

## AUTHORS CONTRIBUTION

Conceptualization : MS, IF, ELR, Methodology : MD, MW, LD, MS, IF, JP, SA, NO, PDC Software : LD, MW, TC, FZ; Validation: MD, LD, CAM,FE Formal Analysis : MS, MD, LD, TC, MW; Investigation : MD, MW, LD, ELR, IF, MS, Resources : Data curation : MW, MS, LD, MD, Writing : MS, MD, ELR, MW, TC, LD, MW Original Draft : MS, LD, MD, MW Supervision, Project : MW, ELR, IF, MS Administration : ELR, MS, IF Funding Acquisition IF, MS; MW

## Supplementary Figures

**Supp. Figure 1. A.** Scanned pictures after hematoxylin-eosin staining of the 46 glioblastoma tumors and **B.** Pathologist annotations of the glioblastoma samples. The pathologist annotated the different regions of interest for each sample (tumors, endothelial proliferation, necrosis, infiltration, blood…). The tissues are classified according to the three groups obtained after MALDI-MSI data segmentation (groups Red, Blue and Yellow). **C.** Spatial segmentation of all tissues and grouping according to the largest molecular area (+50%) represented in each tissue and **D.** MALDI MSI images of characteristics m/z ions for each group. **E.** Ward clustering method give 3 main branches with same characteristic ions. **F**. Principal component analysis (PCA) of each individual spectrum reveals separation between the three groups.

**Supp Figure 2.** A. Global survival curve of all patients according the Karnofsky indice (b), MGMT statut (c) and resection quality (d).

**Supp. Figure 3. A. a)** Analysis of proteins overexpressed in group A shows an involvement in axon guidance. **b)** Proteins overexpressed in group B and mainly involved in complements, coagulation cascade and inflammation **c)** Analysis of overexpressed proteins in group C shows a network of proteins involved in Epstein barr infection. **B.** Correlation between Grancalcin and CAPG expression and glioma patient survival according the TGCA data. Patients were divided based on level of expression into “low” or “high”. **C.** Analysis of overall survival of patients reveals no significant difference between the 3 proteomic groups.ressed in group A shows an involvement in axon guidance. **b)** Proteins overexpressed in group B and mainly involved in complements, coagulation cascade and inflammation **c)** Analysis of overexpressed proteins in group C shows a network of proteins involved in Epstein barr infection. **B.** Correlation between Grancalcin and CAPG expression and glioma patient survival according the TGCA data. Patients were divided based on level of expression into “low” or “high”. **C.** Analysis of overall survival of patients reveals no significant difference between the 3 proteomic groups.

## DATA AVAILIBILITY

Proteomic datasets including MaxQuant files and annotated MS/MS datasets were uploaded to the ProteomeXchange Consortium via the PRIDE database, and then assigned the dataset identifier PXD016165”.

