## Supplementary material for "Overall patient’s survival of glioblastoma associated to molecular markers: a pan-proteomic prospective study": manuscript

**Supplementary Note 1.** Immunohistochemistry analyses, deoxyribonucleic acid (DNA) extraction and quantification of tumor samples

**Supplementary Table 1.** Identified specific proteins per patients

**Supplementary Table 2.** Ghost proteins overexpressed in the 3 groups A B C

**Supplementary Table 3.** 282 of the identified proteins correspond to genes involved in glioma in TGCA database. These proteins were identified from the entire proteomic dataset of the 147 samples

**Supplementary Table 4.** Ghost proteins linked to overall survival and the 3 groups

**Supplementary Figure 1.** Classification of the 46 tumors by MALDI-MSI and relation to the anatomopathologist annotations.

**Supplementary Figure 2.** Survival curve of all patients according the Karnofsky performance status, MGMT promoter methylation status and quality of resection.

**Supplementary Figure 3.** String analysis of overexpressed proteins in each of the three proteomic groups and relation to patient survival.

**Supplementary Note 1.** Immunohistochemistry analyses, deoxyribonucleic acid (DNA) extraction and quantification of tumor samples

**Immunohistochemistry analyses**

ATRX nuclear expression were determined by immunohistochemistry on formalin-fixed paraffin-embedded (FFPE) tumor tissue samples. Four µm sections were labeled in an Ultra automate (Ventana-Roche Tissue Diagnostics, Tucson AZ), after antigen retrieval procedures (ATRX: citrate pH 6.0), according to suppliers protocols. ATRX was determined as positive when cases with more than 10% positive tumor cells were scored positive (ATRX expression). ATRX was explored using, ref HPA001906 Rabbit polyclonal, at dilution 1/200 dilution.

**Deoxyribonucleic acid (DNA) extraction and quantification**

Molecular analyses were performed on FFPE tissues. The following tests were performed: Comparative genomic hybridization (CGH)-array, O^6^-methylguanine-DNA methyltransferase (MGMT) promoter methylation. All tissues used for DNA extraction were histologically evaluated to determine the tumor cell content. Analysis were perfromed on all tissue samples. Samples with less than 40% of tumor cells content were considered as not interpretable when no molecular abnormalitities were found. DNA extraction from FFPE was performed using the kit QIAamp DNA FFPE Tissue (Qiagen). CGH Profiles were determined using a SurePrint G3 Human CGH Microarray Kit, 8x60K (Aligent) and the CytoGenomics v2.7 software. The limit of resolution was 1 Mb. Presence of 1p/19q codeletion, gain of chromosome 7, loss of chromosome 10, amplification of the EGFR gene and homozygous deletion of the Cyclin-Dependent Kinase Inhibitor 2A (CDKN2A) gene was systematically evaluated. The MGMT promoter methylation status (CpGs 74-78) was determined after bisulfite treatment by pyrosequencing on a PyroMark Q96 with kit MGMT PyroMark (Qiagen). The presence of a methylation was score positive when a minimum of 8% of methylation was observed.

**Supplementary Table 1.** Identified specific proteins per patients

| Patient Number | **DNA damage resistance** | **Transcription factors modulators** | **Translation factors modulators** | **Enzymes** | **Other**  **(immunity, tumor suppressor, uncharacterized factor)** |
| --- | --- | --- | --- | --- | --- |
| 2 |  |  |  | MMP12 | C1QA |
| 3 |  | Major centromere autoantigen B;  Transcription factor SOX-8 | Ribosomal protein L7 like; U6 snRNA-associated Sm-like protein LSm4 | Pseudouridylate synthase 7 |  |
| 4 |  |  |  | COMM domain-containing protein 10 | LIM domain-containing protein 1 |
| 5 |  | Transcription elongation factor SPT4;  CD2 antigen cytoplasmic tail-binding protein 2 |  | Rho GTPase-activating protein 31; ATP-dependent RNA helicase DDX19B; Mitochondrial intermediate peptidase; Serpin B8 | Uncharacterized protein C2orf72 |
| 6 |  |  |  |  | Actin, alpha skeletal muscle; |
| 8 | TELO2-interacting protein 1 homolog | Histone H1t; Far upstream element-binding protein 1; Nucleoside diphosphate kinase B |  |  | RalA-binding protein 1; RAB9B; VAMP8; Centromere protein V; Exosome complex component RRP41; HMGH5; Collagen alpha-6(VI) chain; CLIC2; cAMP-dependent protein kinase catalytic subunit gamma; HLA-A3; MICOS complex subunit MIC60; Copper Actin, cytoplasmic 2; chaperone for superoxide dismutase; Protocadherin gamma-B2 |
| 9 |  | Heterogeneous nuclear ribonucleoprotein D-like; Transcription initiation factor TFIID subunit 9 |  |  | Haptoglobin; ATP6V0D2; proline-rich AKT1 substrate 1; Adenylate kinase isoenzyme 6 |
| 15 |  |  |  |  | Tenascin-N |
| 17 | DR1 |  |  |  | Programmed cell death protein 2-like; ImmuNotglobulin-like domain-containing receptor 2; PDE3A; AHDC1 |
| 22 |  |  |  |  | Cell migration-inducing and hyaluronan-binding protein; GTPase IMAP family member 1; |
| 27 |  |  |  |  | Transforming growth factor beta-1; Syntaxin 11; SELP - P-selectin; MelaNotma antigen preferentially expressed in tumors; Trem-like transcript 1 protein; Caveolae-associated protein; Perlaxin; Endoplasmic reticulum-Golgi intermediate compartment protein 2 |
| 33 |  |  |  |  | HLA-A3; Interleukin-1 receptor accessory protein-like 1 |
| 37 |  |  |  |  | Potassium voltage-gated channel subfamily C member 3; CPSF2 |
| 41 |  |  |  |  | Src kinase-associated phosphopr Sprouty-related, EVH1 domain-containing protein 1; RAB39A |
| 49 |  |  |  | Brain-enriched guanylate kinase-associated protein; Ephrin type-B receptor 3 | HLA-B7 |

**Supplementary Table 2. Clinical characteristics**

|  | **Total population (n=46)** |
| --- | --- |
| **Gender**  female, n (%)  male, n (%) | 15 (33)  31 (67) |
| **Age at diagnosis (years)**  median (IQR) | 60 (51-66) |
| **Karnofsky performance score at diagnosis**  median (IQ)  0-80, n (%)  90-100, n (%) | 90 (80-90)  14 (30)  32 (70) |
| **Main location of the tumor**  frontal, n (%)  occipital, n (%)  parietal, n (%)  temporal, n (%) | 11 (24)  3 (6)  12 (26)  20 (43) |
| **Type of surgical resection**  complete, n (%)  partial, n (%)  biopsies, n (%) | 26 (57)  19 (41)  1 (2) |
| **Nuclear expression of ATRX**  lost, n (%)  maintained, n (%) | 4 (9)  42 (91) |
| **MGMT promoter methylation status**  not methylated, n (%)  methylated, n (%) | 31 (67)  15 (33) |
| **EGFR amplification**  no, n (%)  yes, n (%) | 22 (48)  24 (52) |
| **Chromosome 7 gain combined with chromosome 10 loss (+7/-10)**  no, n (%)  yes, n (%) | 12 (26)  34 (74) |
| **EGFR amplification combined with 7 gain / 10 loss**  EGFR amplification or gain 7 / lost 10  EGFR amplification without gain 7 / lost 10  EGFR amplification and gain 7 / lost 10  gain 7 / lost 10 without EGFR amplification | 41 (89)  7 (15)  17 (37)  17 (37) |
| **Homozygous CDKN2A deletion**  no, n (%)  yes, n (%) | 18 (39)  28 (61) |
| **Median follow-up (months)**  median (IQR) | 19.4 (13.5-32.0) |
| **Initial treatment**  RT/TMZ followed by 6 cycles of TMZ, n (%)  RT/TMZ then more than 6 months TMZ, n (%)  RT/TMZ followed by less than 6 cycles of TMZ, n (%)  other treatment*, n (%)  clinical study, n (%)  no treatment, n (%) | 18 (39)  4 (9)  20 (43)  2 (4)  1 (2)  1 (2) |
| **Progression**  yes, n (%)  no, n (%)  unknown, n (%) | 38 (83)  3 (6)  5 (11) |
| **Progression-free survival (months)**  median (IQR) | 10.6 (7.1-16.3) |
| **Treatment at first progression (n=38)**  yes, n (%)  no, n (%) | 33** (87)  5 (13) |
| **Death**  yes, n (%)  no, n (%) | 39 (85)  7 (15) |
| **Survival from surgery (months)**  median (IQR) | 19.4 (13.5-32.0) |
| **Survival**  upper IQR, n (%)  intermediate IQR, n (%)  lower IQR, n (%) |  |
|  | 12 (26)  23 (50)  11 (24) |

* one patient: RT only, one patient: 6 cycles TMZ then SRT

** including surgery in 4 cases

**Abbreviations:**

EGFR, epidermal growth factor receptor

IQR: interquartile range

MGMT: O^6^-methylguanine DNA methyltransferase

RT: radiotherapy

SRT: stereotactic radiotherapy

TMZ: temozolomide

**Supplementary Table 3. Clinical characteristics of each groups**

| **COLOR GROUPS (n=45)** | **A (n=11)**  (red) | **B (n=9)**  (yellow) | **C (n=22)**  (blue) |
| --- | --- | --- | --- |
| **Gender**  female, n (%)  male, n (%) | 4 (36)  7 (64) | 2 (22)  7 (78) | 7 (32)  15 (68) |
| **Age at diagnosis (years)**  median (IQR) | 50 (46-57) | 60 (59-64) | 63 (51-68) |
| **Karnofsky performance score at diagnosis**  median (IQ)  0-80, n (%)  90-100, n (%) | 90 (65-95)  5 (45)  6 (55) | 90 (90-100)  0  9 (100) | 90 (80-90)  7 (32)  15 (68) |
| **Main location of the tumor**  frontal, n (%)  occipital, n (%)  parietal, n (%)  temporal, n (%) | 2 (18)  1 (9)  4 (36)  4 (36) | 1 (11)  0  3 (33)  5 (56) | 7 (32)  2 (9)  4 (18)  9 (9) |
| **Type of surgical resection**  complete, n (%)  partial, n (%)  biopsies, n (%) | 4 (36)  7 (64)  0 | 8 (89)  1 (11)  0 | 10 (45)  11 (50)  1 (5) |
| **Nuclear expression of ATRX**  lost, n (%)  maintained, n (%) | 0  11 (100) | 0  9 (100) | 3 (14)  19 (86) |
| **MGMT promoter methylation status**  not methylated, n (%)  methylated, n (%) | 10 (91)  1 (9) | 4 (44)  5 (56) | 13 (59)  9 (41) |
| **EGFR amplification**  no, n (%)  yes, n (%) | 6 (55)  5 (45) | 5 (56)  4 (44) | 8 (36)  14 (64) |
| **Chromosome 7 gain combined with chromosome 10 loss (+7/-10)**  no, n (%)  yes, n (%) | 3 (27)  8 (73) | 1 (11)  8 (89) | 8 (36)  14 (64) |
| **EGFR amplification combined with 7 gain / 10 loss**  EGFR amplification or gain 7 / lost 10  EGFR amplification without gain 7 / lost 10  EGFR amplification and gain 7 / lost 10  gain 7 / lost 10 without EGFR amplification | 8 (73)  0  7 (64)  3 (27) | 8 (89)  0 (0)  4 (44)  4 (44) | 21 (95)  7 (32)  7 (32)  7 (32) |
| **Homozygous CDKN2A deletion**  no, n (%)  yes, n (%) | 5 (45)  6 (55) | 3 (33)  6 (67) | 7 (32)  15 (68) |
| **Median follow-up (months)**  median (IQR) | 22.3 (11.8-34.9) | 23 (15-29) | 19.5 (14.7-32) |
| **Initial treatment**  RT/TMZ followed by 6 cycles of TMZ, n (%)  RT/TMZ followed by more than 6 cycles of TMZ, n (%)  RT/TMZ followed by less than 6 cycles of TMZ, n (%)  other treatment*, n (%)  clinical study, n (%)  no treatment, n (%) | 4 (36)  1 (9)  4 (36)  2 (18)  0  0 | 6 (67)  0  2 (22)  0  1 (11)  0 | 8 (36)  3 (14)  10 (45)  0  0  1 (5) |
| **Progression**  yes, n (%)  no, n (%)  unknown, n (%) | 11 (100)  0  0 | 7 (78)  1 (11)  1 (11) | 17 (77)  2 (9)  3 (14) |
| **Progression-free survival (months)**  median (IQR) | 9.1 (6.3-21.7) | 12.6 (10-19) | 10 (5.7-14.5) |
| **Treatment at first progression**  yes, n (%)  no, n (%) | 10 (91)  1 (9) | 7 (100)  0 | 14 (64)  3 (14) |
| **Death**  yes, n (%)  no, n (%) | 10 (91)  1 (9) | 7 (78)  2 (22) | 18 (82)  4 (18) |
| **Survival from surgery (months)**  median (IQR) | 22.3 (11.8-34.9) | 23 (15-29) | 19.4 (14.8-32) |
| **Survival**  upper IQR, n (%)  intermediate IQR, n (%)  lower IQR, n (%) | 4 (36)  3 (27)  4 (36) | 2 (22)  6 (67)  1 (11) | 5 (23)  12 (52)  5 (23) |
| **Epilepsy**  no  before diagnosis or at diagnosis  during follow-up | 6 (55)  4 (36)  1 (9) | 2 (22)  4 (44)  3 (33) | 10 (45)  6 (27)  6 (27) |
| **Thromboembolic event**  no  before diagnosis  during follow-up | 11 (100)  0  0 | 8 (89)  0  1 (11) | 20 (91)  1 (5)  1 (5) |

* one patient: RT only, one patient: 6 cycles TMZ then SRT

**Abbreviations:**

EGFR, epidermal growth factor receptor

IDH: isocitrate dehydrogenase

IQR: interquartile range

MGMT: O^6^-methylguanine DNA methyltransferase

RT: radiotherapy

TMZ: temozolomide

**Supplementary Table 4.** Ghost proteins overexpressed in the 3 groups A B C

| GROUP | protein accession | protein length (a.a.) | molecular weight (kDa) | isoelectric point | gene symbol | type | localization |
| --- | --- | --- | --- | --- | --- | --- | --- |
| C | IP_063564 | 113 | 12.39 | 9.26 | CLDN19 | mRNA | 3'UTR |
| C | IP_065285 | 48 | 5.17 | 8.97 | C8B | mRNA | CDS |
| C | IP_072691 | 36 | 4.26 | 8.51 | CD84 | mRNA | 3'UTR |
| C | IP_073718 | 85 | 9.63 | 10.84 | CCDC181 | mRNA | CDS |
| C | IP_079312 | 388 | 42.34 | 5.71 | EDARADD | mRNA | 3'UTR |
| C | IP_092740 | 46 | 5.08 | 10.69 | TTN | mRNA | CDS |
| C | IP_219633 | 35 | 3.95 | 5.67 | POSTN | mRNA | 3'UTR |
| C | IP_2277691 | 46 | 5.53 | 8.78 | RTTN | mRNA | CDS |
| C | IP_2303139 | 56 | 6.24 | 6.49 | LOC105369735 | ncRNA | - |
| C | IP_2323408 | 61 | 6.86 | 8.55 | LOC105376126 | ncRNA | - |
| C | IP_2348387 | 31 | 3.34 | 7.26 | COL19A1 | mRNA | 5'UTR |
| C | IP_235425 | 38 | 4.24 | 10.31 | HCN4 | mRNA | 3'UTR |
| C | IP_2376460 | 57 | 6.73 | 7.36 | EPHA6 | mRNA | 3'UTR |
| C | IP_2393992 | 43 | 5.22 | 11.65 | LOC105376830 | ncRNA | - |
| C | IP_256988 | 39 | 4.52 | 9.45 | TBX21 | mRNA | 3'UTR |
| C | IP_261897 | 41 | 4.44 | 9.67 | AC108004.2 | ncRNA | - |
| C | IP_557239 | 90 | 10.19 | 9.53 | TUBB4AP1 | ncRNA | - |
| C | IP_559054 | 30 | 3.56 | 4.27 | SLITRK2 | mRNA | 5'UTR |
| C | IP_563986 | 424 | 47.8 | 4.43 | KRT8P11 | ncRNA | - |
| C | IP_572422 | 212 | 24.18 | 4.64 | TUBBP1 | ncRNA | - |
| C | IP_580018 | 247 | 29.04 | 11.36 | RPL7P32 | ncRNA | - |
| C | IP_590938 | 74 | 8.75 | 10.05 | HMGB3P18 | ncRNA | - |
| C | IP_629960 | 30 | 3.57 | 9.69 | LRRC34 | ncRNA | - |
| C | IP_669601 | 75 | 8.32 | 10.63 | HIST2H2BB | ncRNA | - |
| C | IP_671464 | 249 | 26.94 | 8.21 | TPI1P1 | ncRNA | - |
| C | IP_701463 | 179 | 20.51 | 9.05 | CTD-2541J13.2 | ncRNA | - |
| C | IP_739889 | 139 | 15.18 | 5.67 | GOLGA6L9 | ncRNA | - |
| C | IP_752010 | 59 | 6.39 | 10.62 | MIPOL1 | mRNA | 3'UTR |
| C | IP_774693 | 75 | 8.68 | 3.92 | TUBAP2 | ncRNA | - |
| C | IP_774695 | 374 | 41.01 | 5.68 | TUBAP2 | ncRNA | - |
| A | IP_2390879 | 36 | 4.08 | 5.49 | LOC107985743 | ncRNA | - |
| A | IP_244732 | 78 | 8.56 | 7.13 | KIFC3 | mRNA | 5'UTR |
| B | IP_156671 | 120 | 14.22 | 10.43 | SLC13A1 | mRNA | 3'UTR |
| B | IP_222588 | 58 | 6.32 | 11.27 | CPB2-AS1 | ncRNA | - |
| B | IP_2389079 | 29 | 2.98 | 8.35 | MYT1L | mRNA | 5'UTR |
| B | IP_265416 | 32 | 3.81 | 10.78 | LIPG | mRNA | CDS |
| B | IP_273562 | 76 | 8.34 | 11.89 | LGI4 | mRNA | 5'UTR |
| B | IP_278725 | 69 | 7.77 | 10.83 | ZNF888 | mRNA | CDS |
| B | IP_592742 | 86 | 9.64 | 5.86 | KRT18P64 | ncRNA | - |
| B | IP_602497 | 38 | 4.24 | 5.65 | CTD-2151A2.2 | ncRNA | - |
| B | IP_672223 | 178 | 20.37 | 5.14 | GBP1P1 | ncRNA | - |

**Supplementary Table 5.** 282 of the identified proteins correspond to genes involved in glioma in TGCA database. These proteins were identified from the entire proteomic dataset of the 147 samples

| Majority protein IDs | Protein names | Gene names | Overexpressed in |
| --- | --- | --- | --- |
| P49418 | Amphiphysin | AMPH | A |
| Q01484 | Ankyrin-2 | ANK2 | A |
| Q8N7J2 | APC membrane recruitment protein 2 | AMER2 | A |
| Q8N126 | Cell adhesion molecule 3 | CADM3 | A |
| Q8N111 | Cell cycle exit and neuronal differentiation protein 1 | CEND1 | A |
| Q12860 | Contactin-1 | CNTN1 | A |
| Q8TAM6 | Ermin | ERMN | A |
| P09972 | Fructose-bisphosphate aldolase C | ALDOC | A |
| P46821 | Microtubule-associated protein 1B | MAP1B | A |
| P11137 | Microtubule-associated protein 2 | MAP2 | A |
| P60201 | Myelin proteolipid protein | PLP1 | A |
| P20916 | Myelin-associated glycoprotein | MAG | A |
| O94856 | Neurofascin | NFASC | A |
| Q15818 | Neuronal pentraxin-1 | NPTX1 | A |
| Q9UH03 | Neuronal-specific septin-3 | sept-03 | A |
| Q9UN36 | Protein NDRG2 | NDRG2 | A |
| Q9Y2J0 | Rabphilin-3A | RPH3A | A |
| P50993 | Sodium/potassium-transporting ATPase subunit alpha-2 | ATP1A2 | A |
| P13637 | Sodium/potassium-transporting ATPase subunit alpha-3 | ATP1A3 | A |
| P14415 | Sodium/potassium-transporting ATPase subunit beta-2 | ATP1B2 | A |
| Q92777 | Synapsin-2 | SYN2 | A |
| Q92752 | Tenascin-R | TNR | A |
| Q9UI15 | Transgelin-3 | TAGLN3 | A |
| P36222 | Chitinase-3-like protein 1 | CHI3L1 | B |
| P01019 | Angiotensinogen | AGT | B |
| P48681 | Nestin | NES | B |
| P14136 | Glial fibrillary acidic protein | GFAP | C |
| Q96PU8 | Protein quaking | QKI | C |
| O94805 | Actin-like protein 6B | ACTL6B | Not specific |
| Q08462 | Adenylate cyclase type 2 | ADCY2 | Not specific |
| P02511 | Alpha-crystallin B chain | CRYAB | Not specific |
| Q99767 | Amyloid beta A4 precursor protein-binding family A member 2 | APBA2 | Not specific |
| P51693 | Amyloid-like protein 1 | APLP1 | Not specific |
| Q7Z6G8 | Ankyrin repeat and sterile alpha motif domain-containing protein 1B | ANKS1B | Not specific |
| P55087 | Aquaporin-4 | AQP4 | Not specific |
| Q99490 | Arf-GAP with GTPase, ANK repeat and PH domain-containing protein 2 | AGAP2 | Not specific |
| O14525 | Astrotactin-1 | ASTN1 | Not specific |
| Q9UBJ2 | ATP-binding cassette sub-family D member 2 | ABCD2 | Not specific |
| Q16620 | BDNF/NT-3 growth factors receptor | NTRK2 | Not specific |
| Q96KN2 | Beta-Ala-His dipeptidase | CNDP1 | Not specific |
| Q96GW7 | Brevican core protein | BCAN | Not specific |
| A6NE02 | BTB/POZ domain-containing protein 17 | BTBD17 | Not specific |
| Q9Y6N8 | Cadherin-10 | CDH10 | Not specific |
| Q9HBT6 | Cadherin-20 | CDH20 | Not specific |
| P55283 | Cadherin-4 | CDH4 | Not specific |
| Q14123 | Calcium/calmodulin-dependent 3,5-cyclic nucleotide phosphodiesterase 1C | PDE1C | Not specific |
| Q8NCB2 | CaM kinase-like vesicle-associated protein | CAMKV | Not specific |
| Q07343 | cAMP-specific 3,5-cyclic phosphodiesterase 4B | PDE4B | Not specific |
| Q9UDT6 | CAP-Gly domain-containing linker protein 2 | CLIP2 | Not specific |
| P07451 | Carbonic anhydrase 3 | CA3 | Not specific |
| A5YM72 | CarNotsine synthase 1 | CARNS1 | Not specific |
| P26232 | Catenin alpha-2 | CTNNA2 | Not specific |
| Q9UI47 | Catenin alpha-3 | CTNNA3 | Not specific |
| Q9UQB3 | Catenin delta-2 | CTNND2 | Not specific |
| Q86WG3 | Caytaxin | ATCAY | Not specific |
| Q8N3J6 | Cell adhesion molecule 2 | CADM2 | Not specific |
| Q8NFZ8 | Cell adhesion molecule 4 | CADM4 | Not specific |
| P27544 | Ceramide synthase 1 | CERS1 | Not specific |
| Q15782 | Chitinase-3-like protein 2 | CHI3L2 | Not specific |
| O95196 | Chondroitin sulfate proteoglycan 5 | CSPG5 | Not specific |
| P26992 | Ciliary neurotrophic factor receptor subunit alpha | CNTFR | Not specific |
| Q8IUQ0 | Clavesin-1 | CLVS1 | Not specific |
| Q5SYC1 | Clavesin-2 | CLVS2 | Not specific |
| Q2UY09 | Collagen alpha-1(XXVIII) chain | COL28A1 | Not specific |
| Q8WXI2 | Connector enhancer of kinase suppressor of ras 2 | CNKSR2 | Not specific |
| Q02246 | Contactin-2 | CNTN2 | Not specific |
| O95741 | Copine-6 | CPNE6 | Not specific |
| Q9UQ03 | Coronin-2B | CORO2B | Not specific |
| Q9ULE3 | DENN domain-containing protein 2A | DENND2A | Not specific |
| Q9Y6T7 | Diacylglycerol kinase beta | DGKB | Not specific |
| P49619 | Diacylglycerol kinase gamma | DGKG | Not specific |
| P42658 | Dipeptidyl amiNotpeptidase-like protein 6 | DPP6 | Not specific |
| O95886 | Disks large-associated protein 3 | DLGAP3 | Not specific |
| Q9Y4J8 | Dystrobrevin alpha | DTNA | Not specific |
| Q9C026 | E3 ubiquitin-protein ligase TRIM9 | TRIM9 | Not specific |
| Q6UWR7 | Ectonucleotide pyrophosphatase/phosphodiesterase family member 6 | ENPP6 | Not specific |
| Q14576 | ELAV-like protein 3 | ELAVL3 | Not specific |
| P26378 | ELAV-like protein 4 | ELAVL4 | Not specific |
| Q9Y6R1 | Electrogenic sodium bicarbonate cotransporter 1 | SLC4A4 | Not specific |
| Q2Y0W8 | Electroneutral sodium bicarbonate exchanger 1 | SLC4A8 | Not specific |
| Q9NT22 | EMILIN-3 | EMILIN3 | Not specific |
| P00533 | Epidermal growth factor receptor | EGFR | Not specific |
| Q8TBG4 | EthaNotlamine-phosphate phospho-lyase | ETNPPL | Not specific |
| P43003 | Excitatory amiNot acid transporter 1 | SLC1A3 | Not specific |
| P43004 | Excitatory amiNot acid transporter 2 | SLC1A2 | Not specific |
| Q99689 | Fasciculation and elongation protein zeta-1 | FEZ1 | Not specific |
| O15540 | Fatty acid-binding protein, brain | FABP7 | Not specific |
| Q7Z6J6 | FERM domain-containing protein 5 | FRMD5 | Not specific |
| P05230 | Fibroblast growth factor 1 | FGF1 | Not specific |
| Q9BTV5 | Fibronectin type III and SPRY domain-containing protein 1 | FSD1 | Not specific |
| Q9NZ56 | Formin-2 | FMN2 | Not specific |
| P14867 | Gamma-amiNotbutyric acid receptor subunit alpha-1 | GABRA1 | Not specific |
| P18505 | Gamma-amiNotbutyric acid receptor subunit beta-1 | GABRB1 | Not specific |
| O75899 | Gamma-amiNotbutyric acid type B receptor subunit 2 | GABBR2 | Not specific |
| O75936 | Gamma-butyrobetaine dioxygenase | BBOX1 | Not specific |
| Q96MZ0 | Ganglioside-induced differentiation-associated protein 1-like 1 | GDAP1L1 | Not specific |
| Q6ZMI3 | Gliomedin | GLDN | Not specific |
| Q05329 | Glutamate decarboxylase 2 | GAD2 | Not specific |
| P42262 | Glutamate receptor 2 | GRIA2 | Not specific |
| P42263 | Glutamate receptor 3 | GRIA3 | Not specific |
| P48058 | Glutamate receptor 4 | GRIA4 | Not specific |
| Q13003 | Glutamate receptor ioNottropic, kainate 3 | GRIK3 | Not specific |
| Q9HCC8 | GlycerophosphoiNotsitol iNotsitolphosphodiesterase GDPD2 | GDPD2 | Not specific |
| O95057 | GTP-binding protein Di-Ras1 | DIRAS1 | Not specific |
| Q96HU8 | GTP-binding protein Di-Ras2 | DIRAS2 | Not specific |
| P63215 | Guanine nucleotide-binding protein G(I)/G(S)/G(O) subunit gamma-3 | GNG3 | Not specific |
| O60262 | Guanine nucleotide-binding protein G(I)/G(S)/G(O) subunit gamma-7 | GNG7 | Not specific |
| Q14CZ8 | Hepatocyte cell adhesion molecule | HEPACAM | Not specific |
| Q9UM19 | Hippocalcin-like protein 4 | HPCAL4 | Not specific |
| Q9GZV7 | Hyaluronan and proteoglycan link protein 2 | HAPLN2 | Not specific |
| Q5DX21 | ImmuNotglobulin superfamily member 11 | IGSF11 | Not specific |
| Q96ID5 | ImmuNotglobulin superfamily member 21 | IGSF21 | Not specific |
| Q71H61 | ImmuNotglobulin-like domain-containing receptor 2 | ILDR2 | Not specific |
| Q13683 | Integrin alpha-7 | ITGA7 | Not specific |
| P57087 | Junctional adhesion molecule B | JAM2 | Not specific |
| Q12840 | Kinesin heavy chain isoform 5A | KIF5A | Not specific |
| Q12756 | Kinesin-like protein KIF1A | KIF1A | Not specific |
| Q9NS86 | LanC-like protein 2 | LANCL2 | Not specific |
| O95970 | Leucine-rich glioma-inactivated protein 1 | LGI1 | Not specific |
| Q9NT99 | Leucine-rich repeat-containing protein 4B | LRRC4B | Not specific |
| Q13449 | Limbic system-associated membrane protein | LSAMP | Not specific |
| P06858 | Lipoprotein lipase | LPL | Not specific |
| Q9UKU0 | Long-chain-fatty-acid--CoA ligase 6 | ACSL6 | Not specific |
| Q96GR2 | Long-chain-fatty-acid--CoA ligase ACSBG1 | ACSBG1 | Not specific |
| Q15049 | Membrane protein MLC1 | MLC1 | Not specific |
| Q86UL8 | Membrane-associated guanylate kinase, WW and PDZ domain-containing protein 2 | MAGI2 | Not specific |
| Q9Y2H9 | Microtubule-associated serine/threonine-protein kinase 1 | MAST1 | Not specific |
| P02689 | Myelin P2 protein | PMP2 | Not specific |
| Q16653 | Myelin-oligodendrocyte glycoprotein | MOG | Not specific |
| O15069 | NAC-alpha domain-containing protein 1 | NACAD | Not specific |
| P15882 | N-chimaerin | CHN1 | Not specific |
| P13591 | Neural cell adhesion molecule 1 | NCAM1 | Not specific |
| O00533 | Neural cell adhesion molecule L1-like protein | CHL1 | Not specific |
| Q9ULB1 | Neurexin-1 | NRXN1 | Not specific |
| O14594 | Neurocan core protein | NCAN | Not specific |
| Q8N2Q7 | Neuroligin-1 | NLGN1 | Not specific |
| Q9NZ94 | Neuroligin-3 | NLGN3 | Not specific |
| Q92823 | Neuronal cell adhesion molecule | NRCAM | Not specific |
| P51674 | Neuronal membrane glycoprotein M6-a | GPM6A | Not specific |
| Q13491 | Neuronal membrane glycoprotein M6-b | GPM6B | Not specific |
| O43602 | Neuronal migration protein doublecortin | DCX | Not specific |
| P84074 | Neuron-specific calcium-binding protein hippocalcin | HPCA | Not specific |
| Q99784 | Notelin | OLFM1 | Not specific |
| Q16288 | NT-3 growth factor receptor | NTRK3 | Not specific |
| P23515 | Oligodendrocyte-myelin glycoprotein | OMG | Not specific |
| Q14982 | Opioid-binding protein/cell adhesion molecule | OPCML | Not specific |
| P10451 | Osteopontin | SPP1 | Not specific |
| Q9UL42 | Paraneoplastic antigen Ma2 | PNMA2 | Not specific |
| P26022 | Pentraxin-related protein PTX3 | PTX3 | Not specific |
| Q6UXB8 | Peptidase inhibitor 16 | PI16 | Not specific |
| Q9P0Z9 | Peroxisomal sarcosine oxidase | PIPOX | Not specific |
| Q8IYB4 | PEX5-related protein | PEX5L | Not specific |
| Q9BQI7 | PH and SEC7 domain-containing protein 2 | PSD2 | Not specific |
| Q16816 | Phosphorylase b kinase gamma catalytic chain, skeletal muscle/heart isoform | PHKG1 | Not specific |
| Q96FC7 | PhytaNotyl-CoA hydroxylase-interacting protein-like | PHYHIPL | Not specific |
| P16234 | Platelet-derived growth factor receptor alpha | PDGFRA | Not specific |
| Q9UF11 | Pleckstrin homology domain-containing family B member 1 | PLEKHB1 | Not specific |
| Q9HCM2 | Plexin-A4 | PLXNA4 | Not specific |
| P16389 | Potassium voltage-gated channel subfamily A member 2 | KCNA2 | Not specific |
| O43526 | Potassium voltage-gated channel subfamily KQT member 2 | KCNQ2 | Not specific |
| Q9UL51 | Potassium/sodium hyperpolarization-activated cyclic nucleotide-gated channel 2 | HCN2 | Not specific |
| P20265 | POU domain, class 3, transcription factor 2 | POU3F2 | Not specific |
| Q8TBB6 | Probable cationic amiNot acid transporter | SLC7A14 | Not specific |
| Q8N9I9 | Probable E3 ubiquitin-protein ligase DTX3 | DTX3 | Not specific |
| Q8IYK4 | Procollagen galactosyltransferase 2 | COLGALT2 | Not specific |
| P01213 | Proenkephalin-B | PDYN | Not specific |
| Q13015 | Protein AF1q | MLLT11 | Not specific |
| Q86XD5 | Protein FAM131B | FAM131B | Not specific |
| Q6P995 | Protein FAM171B | FAM171B | Not specific |
| Q9BWQ8 | Protein lifeguard 2 | FAIM2 | Not specific |
| P04271 | Protein S100-B | S100B | Not specific |
| A6NL88 | Protein shisa-7 | SHISA7 | Not specific |
| P60059 | Protein transport protein Sec61 subunit gamma | SEC61G | Not specific |
| Q9H313 | Protein tweety homolog 1 | TTYH1 | Not specific |
| Q9UPW8 | Protein unc-13 homolog A | UNC13A | Not specific |
| Q9Y5G2 | Protocadherin gamma-B2 | PCDHGB2 | Not specific |
| Q9Y5F8 | Protocadherin gamma-B7 | PCDHGB7 | Not specific |
| Q9UN70 | Protocadherin gamma-C3 | PCDHGC3 | Not specific |
| Q52LD8 | Raftlin-2 | RFTN2 | Not specific |
| P50749 | Ras association domain-containing protein 2 | RASSF2 | Not specific |
| Q14088 | Ras-related protein Rab-33A | RAB33A | Not specific |
| Q96DA2 | Ras-related protein Rab-39B | RAB39B | Not specific |
| Q9BRK0 | Receptor expression-enhancing protein 2 | REEP2 | Not specific |
| P23471 | Receptor-type tyrosine-protein phosphatase zeta | PTPRZ1 | Not specific |
| Q16849 | Receptor-type tyrosine-protein phosphatase-like N | PTPRN | Not specific |
| P49758 | Regulator of G-protein signaling 6 | RGS6 | Not specific |
| P49802 | Regulator of G-protein signaling 7 | RGS7 | Not specific |
| Q96B86 | Repulsive guidance molecule A | RGMA | Not specific |
| P12271 | Retinaldehyde-binding protein 1 | RLBP1 | Not specific |
| Q15052 | Rho guanine nucleotide exchange factor 6 | ARHGEF6 | Not specific |
| P51513 | RNA-binding protein Notva-1 | NOTVA1 | Not specific |
| Q9UNW9 | RNA-binding protein Notva-2 | NOTVA2 | Not specific |
| P13521 | Secretogranin-2 | SCG2 | Not specific |
| Q8WXD2 | Secretogranin-3 | SCG3 | Not specific |
| Q9BYH1 | Seizure 6-like protein | SEZ6L | Not specific |
| Q6ZU15 | Septin-14 | sept-14 | Not specific |
| Q8TDC3 | Serine/threonine-protein kinase BRSK1 | BRSK1 | Not specific |
| Q8IWQ3 | Serine/threonine-protein kinase BRSK2 | BRSK2 | Not specific |
| O15075 | Serine/threonine-protein kinase DCLK1 | DCLK1 | Not specific |
| Q8N568 | Serine/threonine-protein kinase DCLK2 | DCLK2 | Not specific |
| Q92529 | SHC-transforming protein 3 | SHC3 | Not specific |
| Q9H156 | SLIT and NTRK-like protein 2 | SLITRK2 | Not specific |
| Q9H2S1 | Small conductance calcium-activated potassium channel protein 2 | KCNN2 | Not specific |
| Q9UGI6 | Small conductance calcium-activated potassium channel protein 3 | KCNN3 | Not specific |
| P30531 | Sodium- and chloride-dependent GABA transporter 1 | SLC6A1 | Not specific |
| P48066 | Sodium- and chloride-dependent GABA transporter 3 | SLC6A11 | Not specific |
| Q99250 | Sodium channel protein type 2 subunit alpha | SCN2A | Not specific |
| Q9NY72 | Sodium channel subunit beta-3 | SCN3B | Not specific |
| Q6U841 | Sodium-driven chloride bicarbonate exchanger | SLC4A10 | Not specific |
| P22732 | Solute carrier family 2, facilitated glucose transporter member 5 | SLC2A5 | Not specific |
| Q14515 | SPARC-like protein 1 | SPARCL1 | Not specific |
| Q86SK9 | Stearoyl-CoA desaturase 5 | SCD5 | Not specific |
| P78539 | Sushi repeat-containing protein SRPX | SRPX | Not specific |
| Q7L0J3 | Synaptic vesicle glycoprotein 2A | SV2A | Not specific |
| Q9HCJ6 | Synaptic vesicle membrane protein VAT-1 homolog-like | VAT1L | Not specific |
| P08247 | Synaptophysin | SYP | Not specific |
| Q9BT88 | Synaptotagmin-11 | SYT11 | Not specific |
| Q8N6N2 | Tetratricopeptide repeat protein 9B | TTC9B | Not specific |
| Q33E94 | Transcription factor RFX4 | RFX4 | Not specific |
| P48431 | Transcription factor SOX-2 | SOX2 | Not specific |
| P57073 | Transcription factor SOX-8 | SOX8 | Not specific |
| P28289 | Tropomodulin-1 | TMOD1 | Not specific |
| Q71U36 | Tubulin alpha-1A chain | TUBA1A | Not specific |
| Q9BVA1 | Tubulin beta-2B chain | TUBB2B | Not specific |
| Q9P2U7 | Vesicular glutamate transporter 1 | SLC17A7 | Not specific |
| O60359 | Voltage-dependent calcium channel gamma-3 subunit | CACNG3 | Not specific |
| Q9Y279 | V-set and immuNotglobulin domain-containing protein 4 | VSIG4 | Not specific |

**Supplementary Table 6.** Alternative proteins linked to overall survival and the 3 groups (A, B and C)

| Group ABC | overall surival cluster | protein accession | protein length (a.a.) | molecular weight (kDa) | isoelectric point | gene symbol | type |
| --- | --- | --- | --- | --- | --- | --- | --- |
| - | 2 | IP_064355 | 62 | 6.75 | 10.17 | MOB3C | mRNA |
| - | 2 | IP_091795 | 37 | 4.52 | 10.36 | SCN9A | mRNA |
| - | 2 | IP_140988 | 33 | 3.64 | 9.54 | TAF8 | mRNA |
| B | 2 | IP_156671 | 120 | 14.22 | 10.43 | SLC13A1 | mRNA |
| - | 2 | IP_186061 | 45 | 5.4 | 8.51 | HECTD2 | mRNA |
| B | 2 | IP_222588 | 58 | 6.32 | 11.27 | CPB2-AS1 | ncRNA |
| - | 2 | IP_2279437 | 67 | 6.74 | 8.54 | AANAT | mRNA |
| - | 2 | IP_2316274 | 45 | 5.04 | 7.62 | LOC105378456 | ncRNA |
| - | 2 | IP_2339128 | 60 | 6.57 | 9.26 | LOC105375130 | ncRNA |
| B | 2 | IP_2389079 | 29 | 2.98 | 8.35 | MYT1L | mRNA |
| - | 2 | IP_2396412 | 53 | 6.07 | 10.63 | ABL2 | mRNA |
| B | 2 | IP_273562 | 76 | 8.34 | 11.89 | LGI4 | mRNA |
| B | 2 | IP_278725 | 69 | 7.77 | 10.83 | ZNF888 | mRNA |
| B | 2 | IP_592742 | 86 | 9.64 | 5.86 | KRT18P64 | ncRNA |
| - | 2 | IP_599034 | 52 | 6.04 | 9.81 | CTB-14A14.2 | ncRNA |
| B | 2 | IP_602497 | 38 | 4.24 | 5.65 | CTD-2151A2.2 | ncRNA |
| - | 2 | IP_609092 | 39 | 4.85 | 6.05 | PCDHGA2 | mRNA |
| - | 2 | IP_652563 | 96 | 10.65 | 10.14 | CTA-373H7.7 | ncRNA |
| - | 2 | IP_711623 | 62 | 6.98 | 12.51 | NPLOC4 | ncRNA |
| - | 2 | IP_724416 | 101 | 10.91 | 11.65 | CTD-2385L22.2 | ncRNA |
| - | 2 | IP_758919 | 79 | 8.77 | 10.39 | FZD10-AS1 | ncRNA |
| - | 2 | IP_774832 | 168 | 17.95 | 8.66 | GAPDHP70 | ncRNA |
| C | - | IP_2277691 | 46 | 5.53 | 8.78 | RTTN | mRNA |
| A | - | IP_2390879 | 36 | 4.08 | 5.49 | LOC107985743 | ncRNA |
| - | - | IP_243434 | 54 | 6.12 | 12.04 | SRCAP | mRNA |
| - | - | IP_557241 | 70 | 7.86 | 9.75 | TUBB4AP1 | ncRNA |
| - | - | IP_741482 | 100 | 10.87 | 12.39 | DPP8 | mRNA |
| C | - | IP_774695 | 374 | 41.01 | 5.68 | TUBAP2 | ncRNA |
| - | - | IP_794851 | 40 | 4.31 | 8.53 | SEC31B | ncRNA |

**Supplementary Figure 1.** Classification of the 46 tumors by MALDI-MSI and relation to the anatomopathologist annotations.

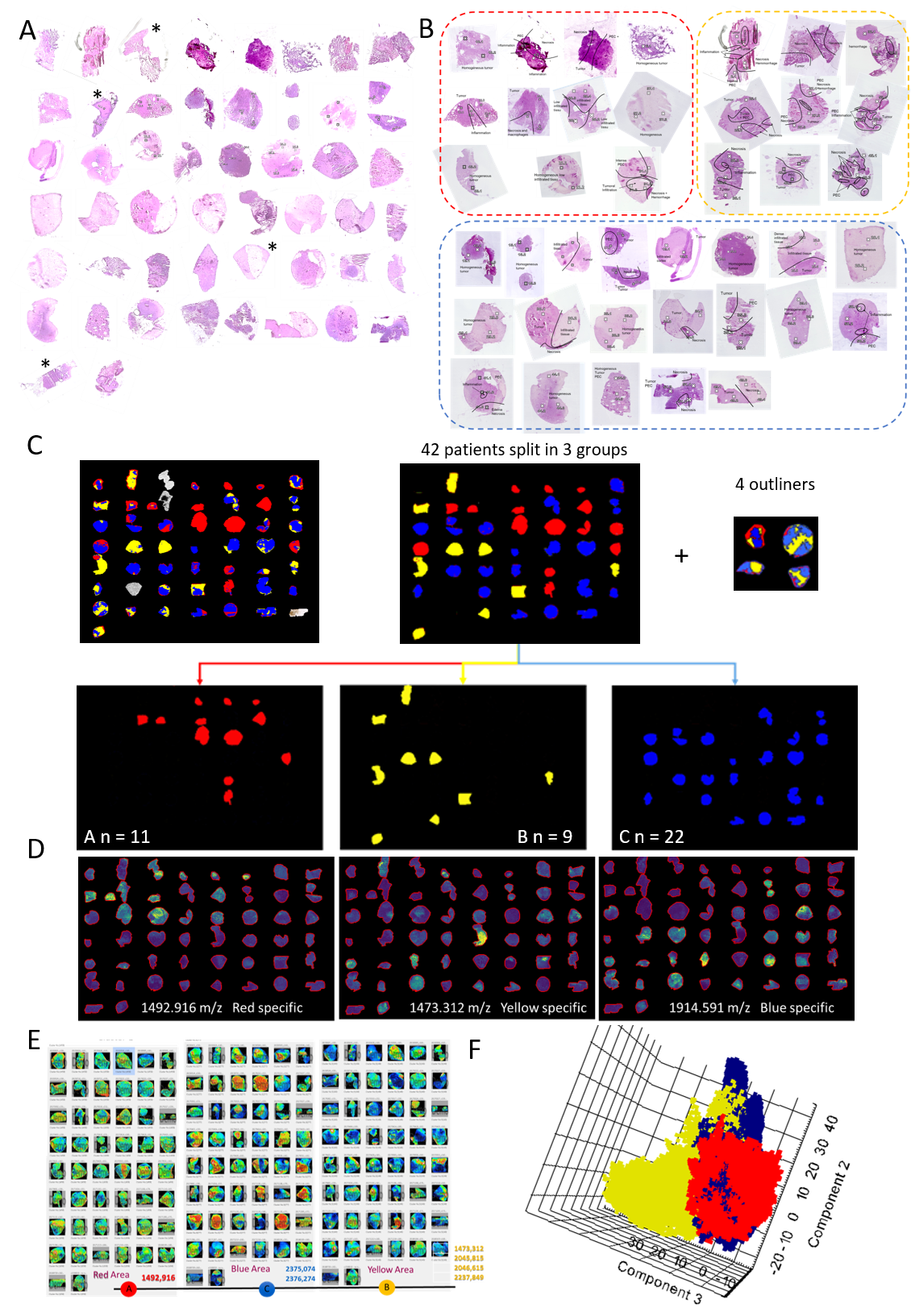

**Legend:**

**A.** Scanned pictures after hematoxylin-eosin staining of the 46 glioblastoma tumors

**B.** Pathologist annotations of the glioblastoma samples. The pathologist annotated the different regions of interest for each sample (tumors, endothelial proliferation, necrosis, infiltration, blood...). The tissues are classified according to the three groups obtained after MALDI-MSI data segmentation (groups Red, Blue and Yellow).

**C.** Spatial segmentation of all tissues and grouping according to the largest molecular area (+50%) represented in each tissue

**D.** MALDI MSI images of characteristics m/z ions for each group.

**E.** Ward clustering method give 3 main branches with same characteristic ions.

**F**. Principal component analysis (PCA) of each individual spectra reveals separation between the three groups.

**Supplementary Figure 2.** Survival curves of all patients according the Karnofsky performance status, MGMT promoter methylation status and quality of resection.

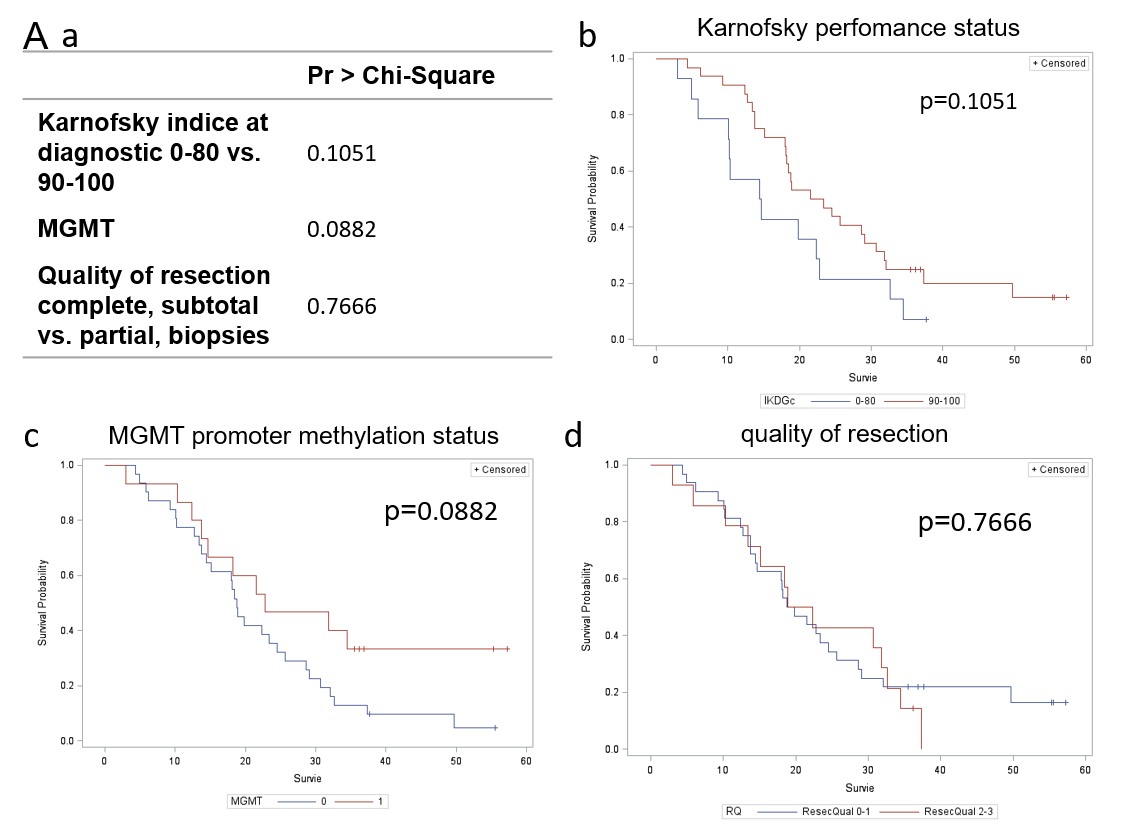

**Legend:**

Survival curve of all patients according the Karnofsky performance status (b), MGMT status (c) and resection quality (d).

**Supplementary Figure 3.** String analysis of overexpressed proteins in each of the three proteomic groups and relation to patient survival

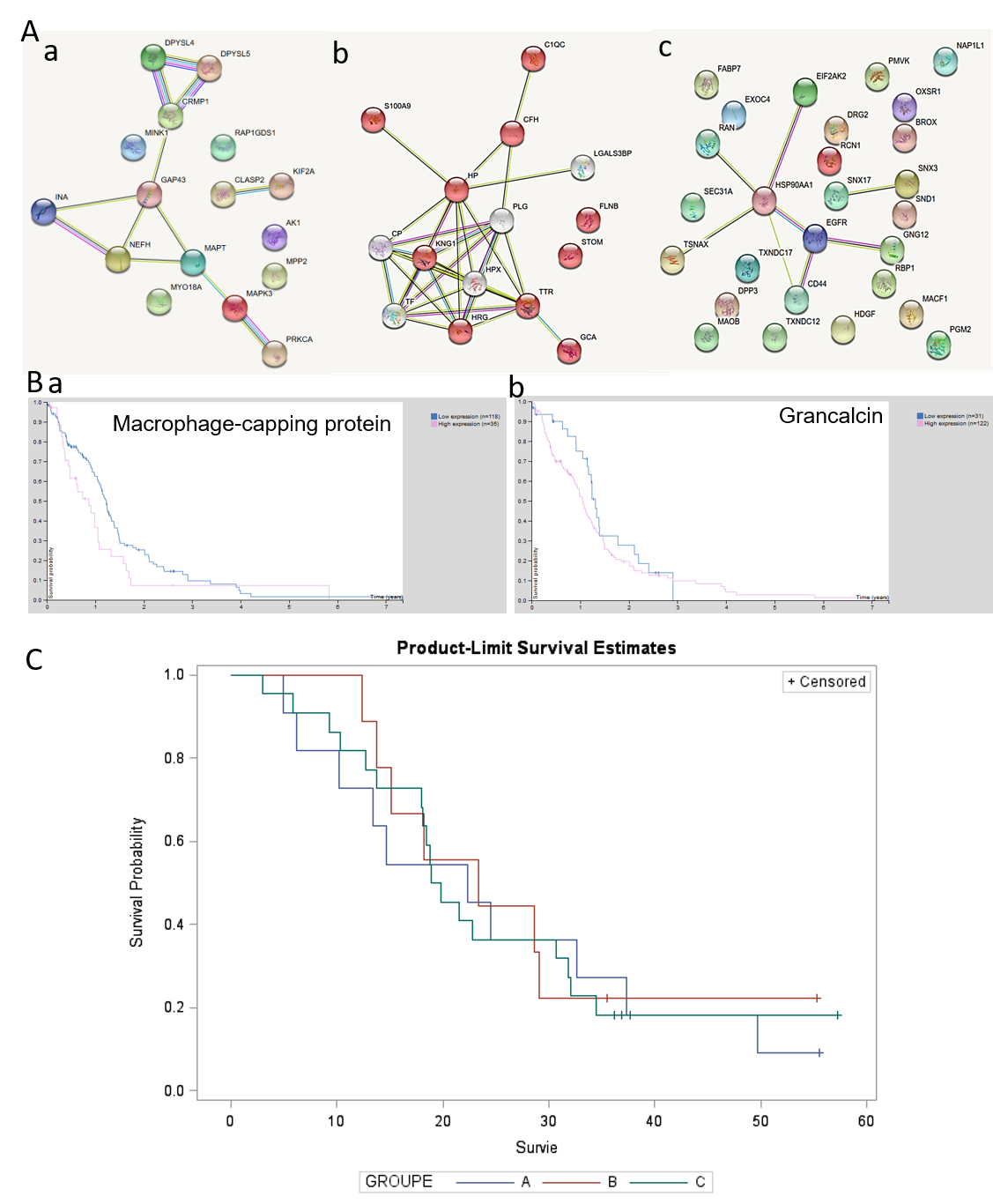

**Legend:**

**A. a)** Analysis of proteins overexpressed in group A shows an involvement in axon guidance. **b)** Proteins overexpressed in group B and mainly involved in complements, coagulation cascade and inflammation **c)** Analysis of overexpressed proteins in group C shows a network of proteins involved in Epstein barr infection. **B.** Correlation between Grancalcin and CAPG expression and glioma patient survival according the TGCA data. Patients were divided based on level of expression into “low” or “high”. **C.** Analysis of overall survival (months) of patients reveals no significant difference between the 3 proteomic groups.
