## Supplementary material for "Overall patient’s survival of glioblastoma associated to molecular markers: a pan-proteomic prospective study": supp data 1

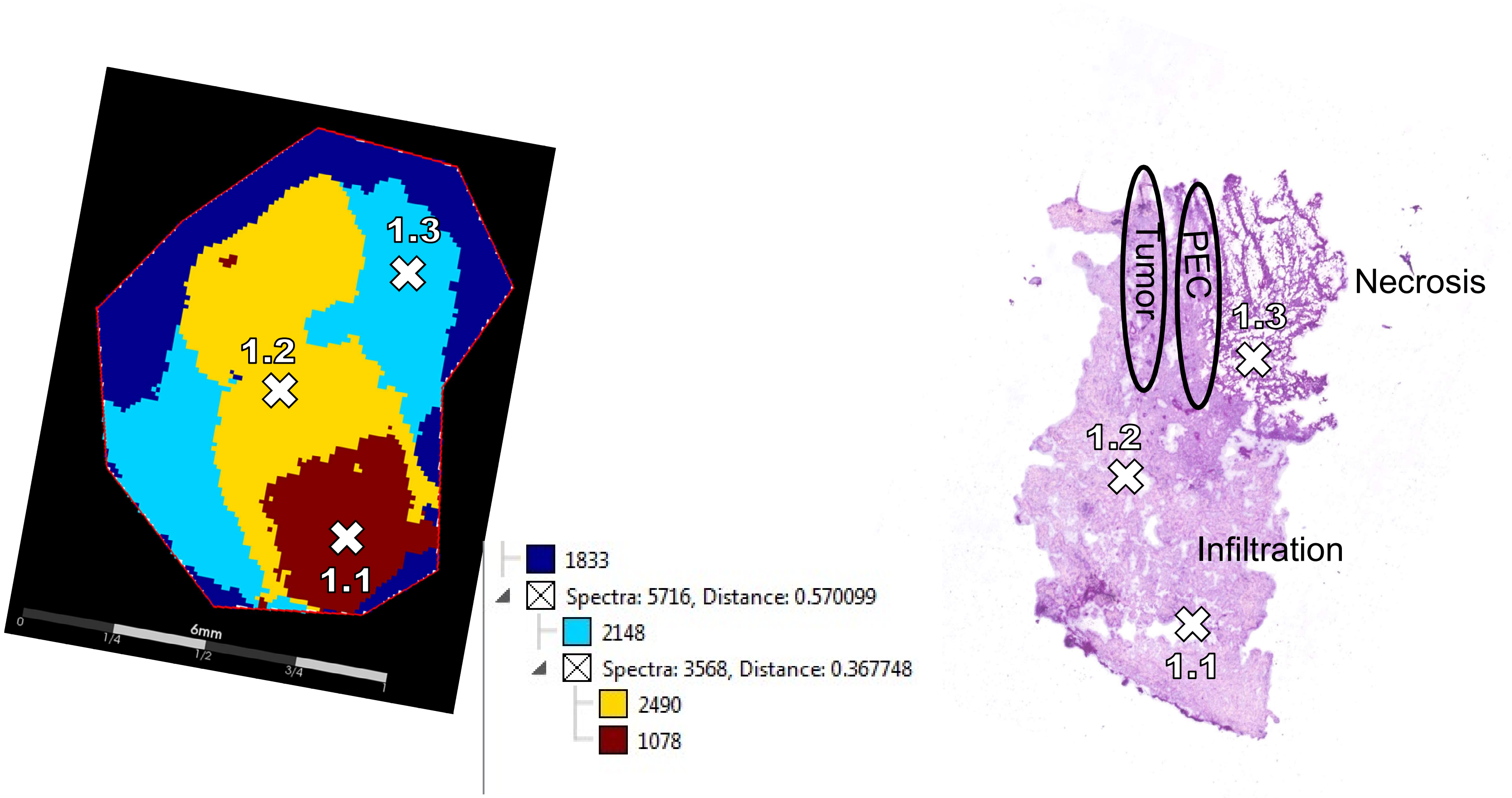
1.3
1.2
1.1
6mm
0
1/4
1/2
3/4
1
1833
Spectra: 5716, Distance: 0.570099
2148
Spectra: 3568, Distance: 0.367748
2490
1078
Tumor
PEC
Necrosis
1.3
1.2
Infiltration
1.1

2

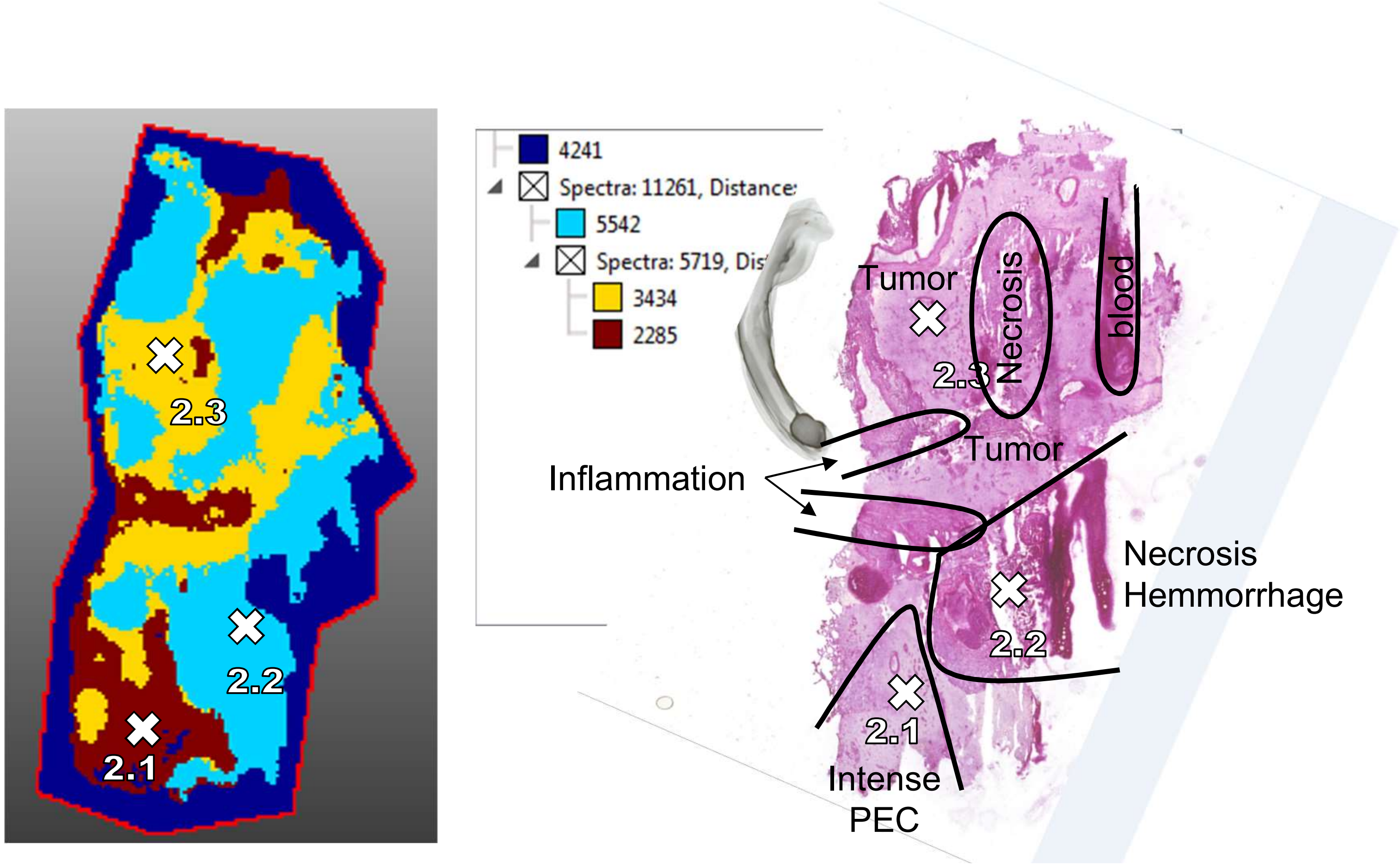

2.3
2.2
2.1
4241
Spectra: 11261, Distance:
5542
Spectra: 5719, Dis
3434
2285
Tumor
Necrosis
blood
2.3
Tumor
Inflammation
Necrosis
Hemmorrhage
2.2
2.1
Intense
PEC

3

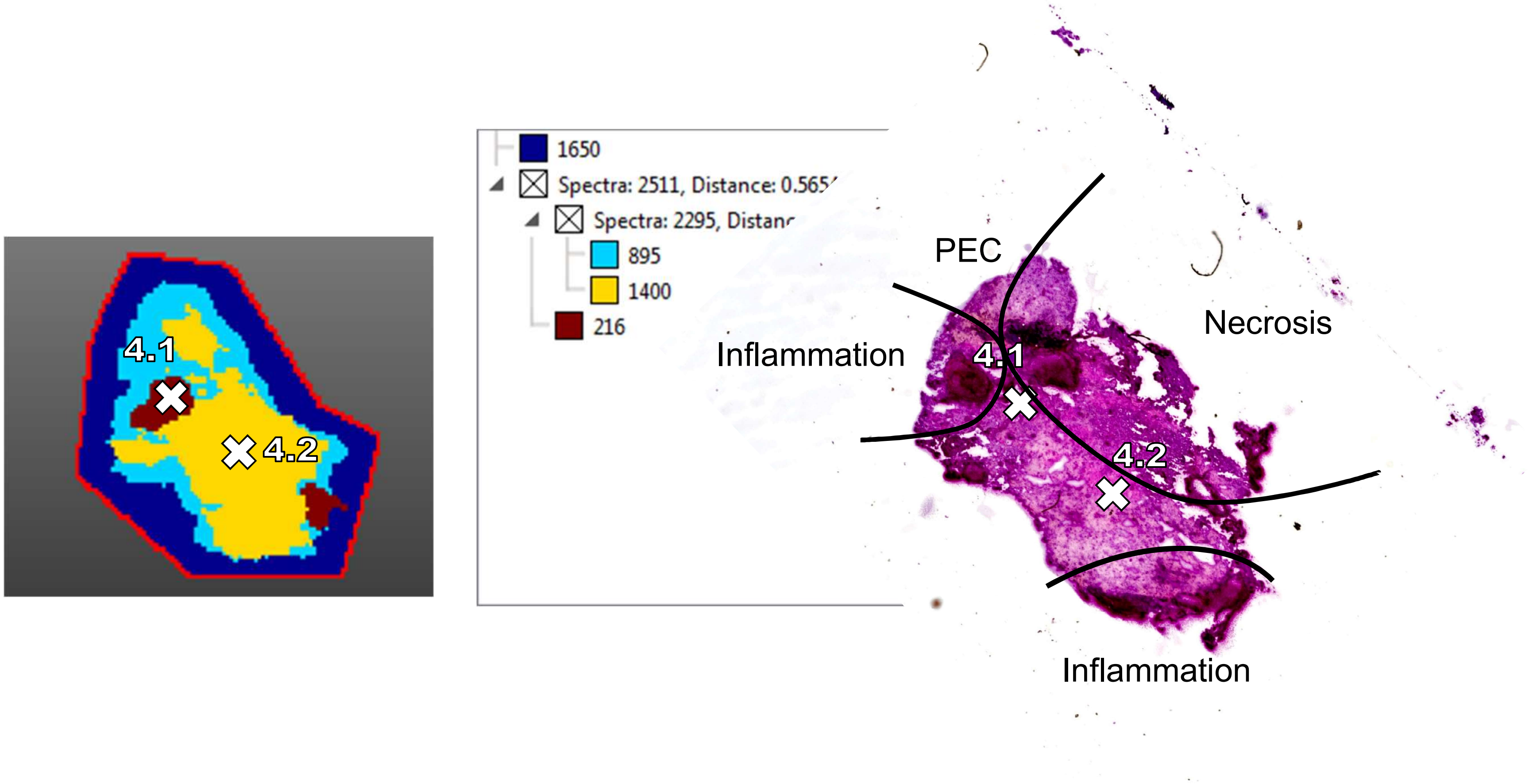
4.1
4.2
1650
Spectra: 2511, Distance: 0.565
Spectra: 2295, Distanc
895
1400
216
PEC
Necrosis
Inflammation
4.1
4.2
Inflammation

4

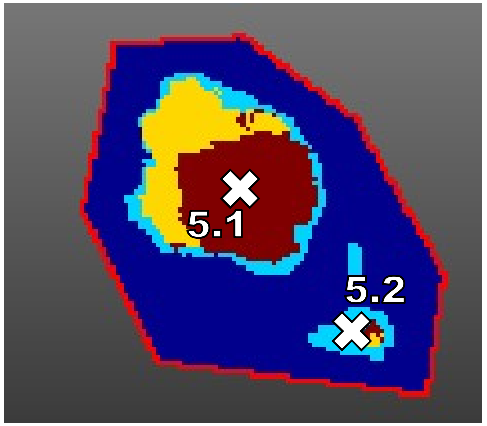


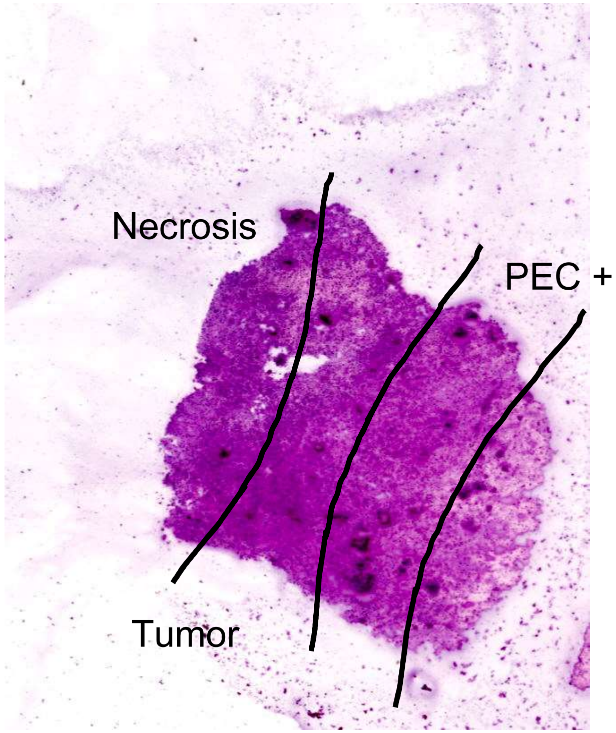

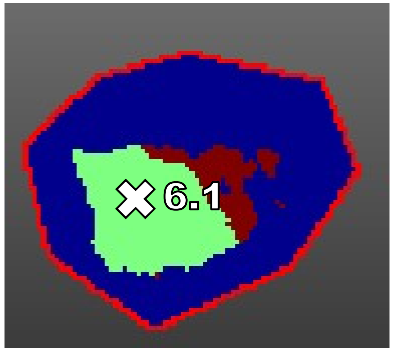


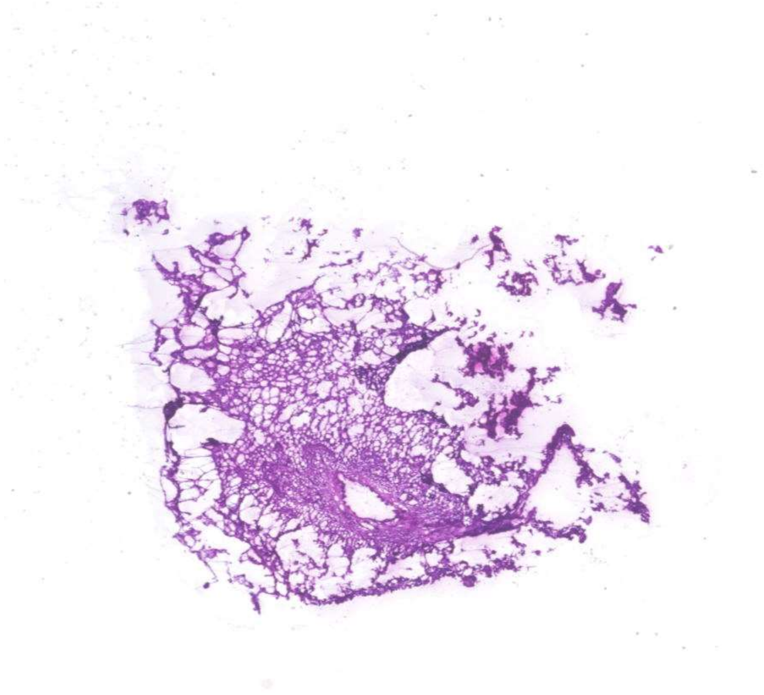

Homogeneous tumor

6

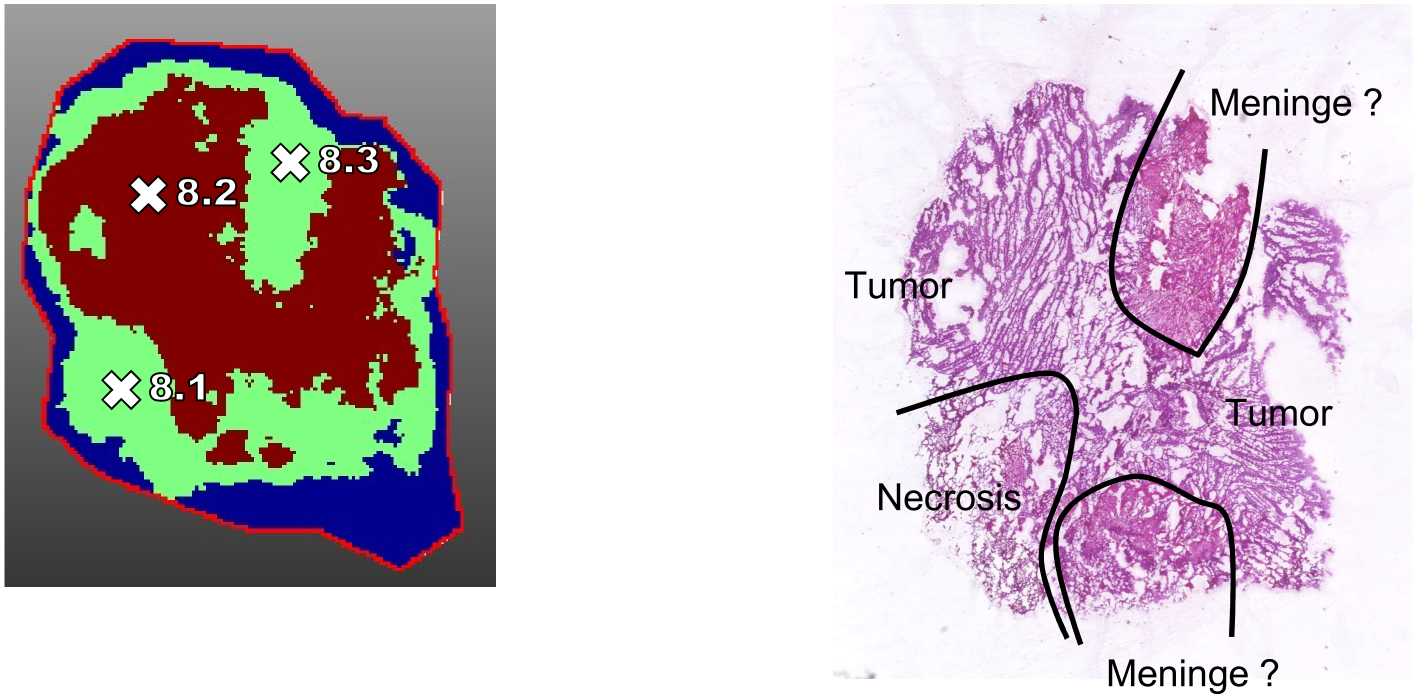

7

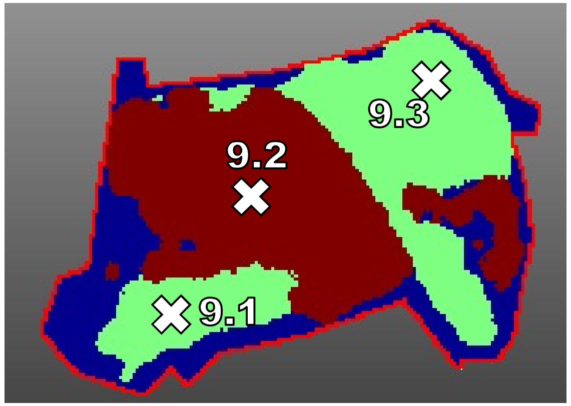


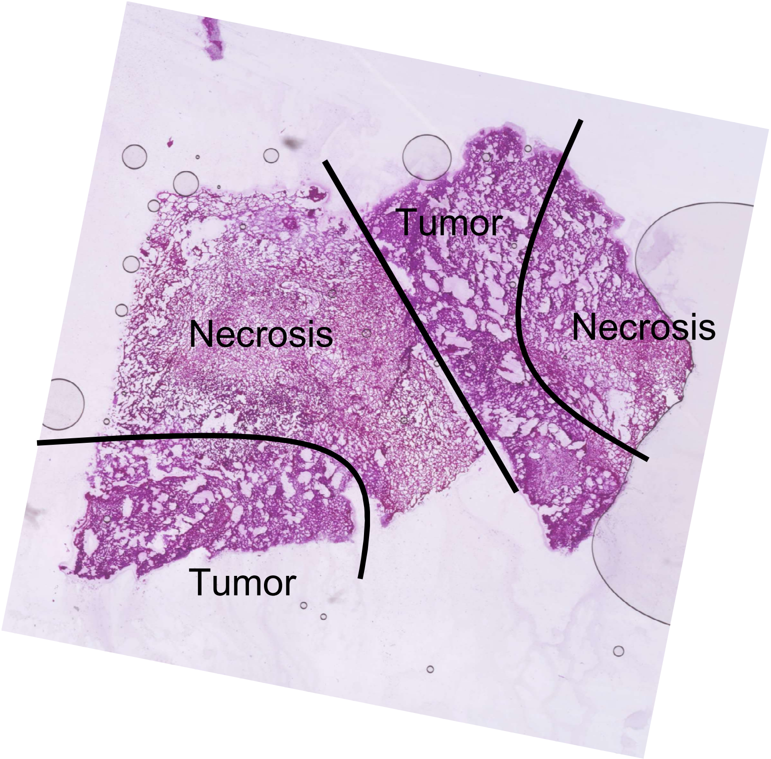

8

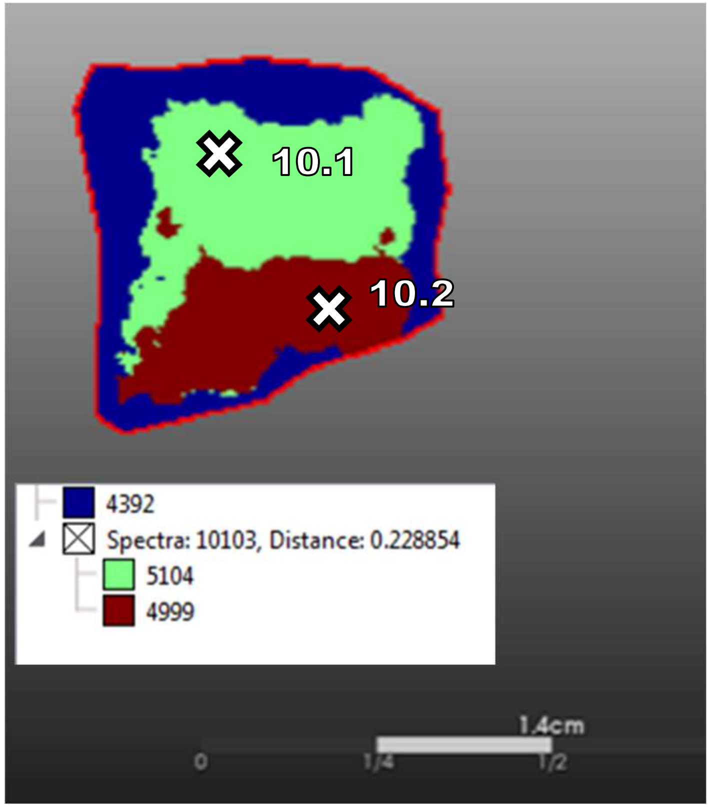


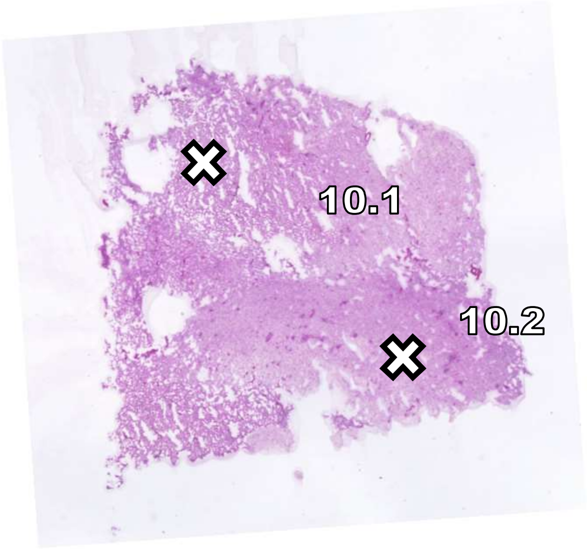


Homogeneous tumor

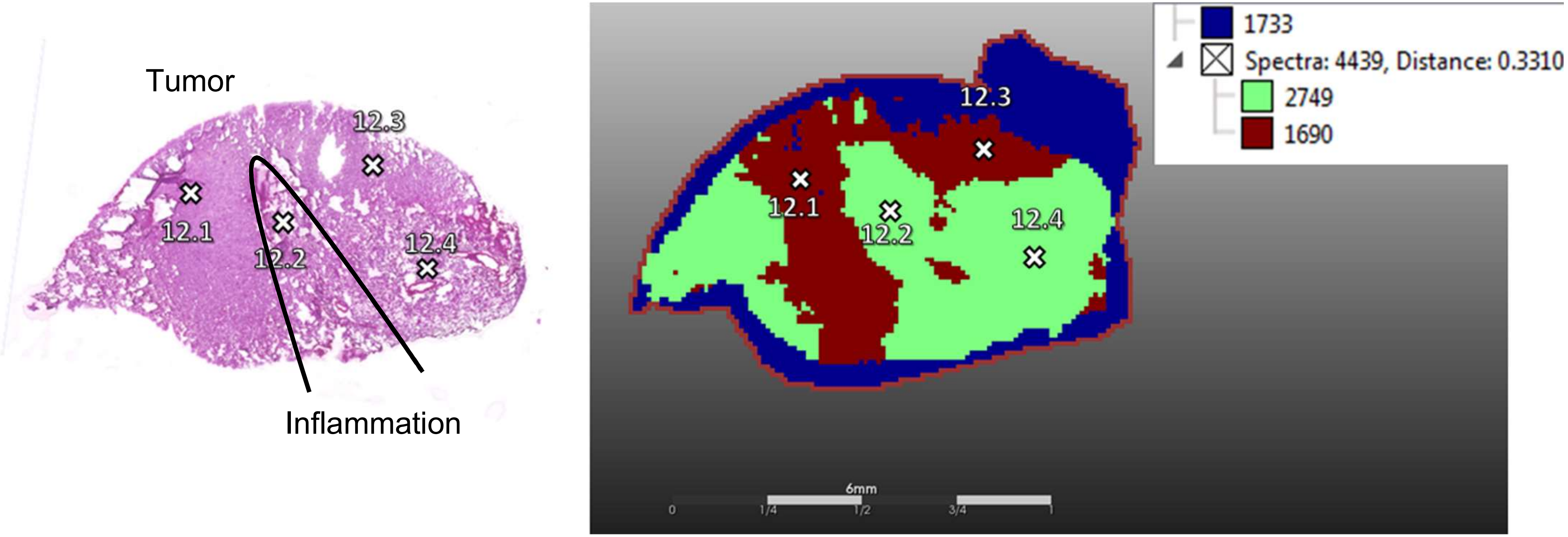
Tumor
12.3
12.1
12.2
12.4
Inflammation
1733
Spectra: 4439, Distance: 0.3310
2749
1690
12.3
12.1
12.2
12.4
6mm
0
1/4
1/2
3/4
1

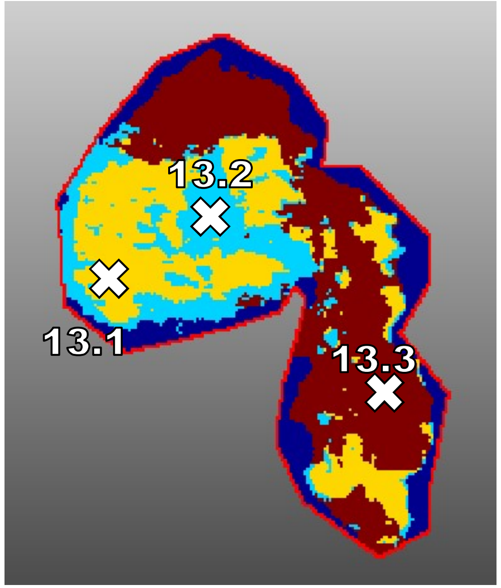
13.2
13.1
13.3

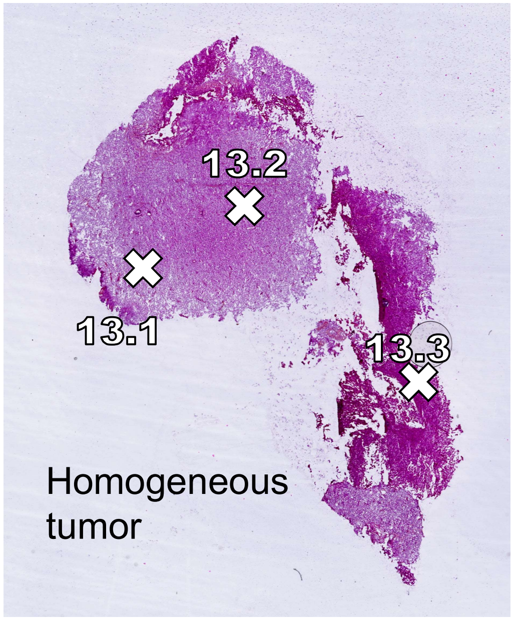
13.2
13.1
13.3
Homogeneous
tumor

11

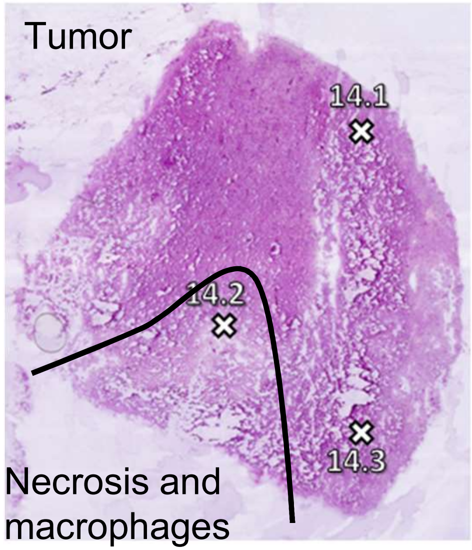
Tumor
14.1
14.2
14.3
Necrosis and
macrophages

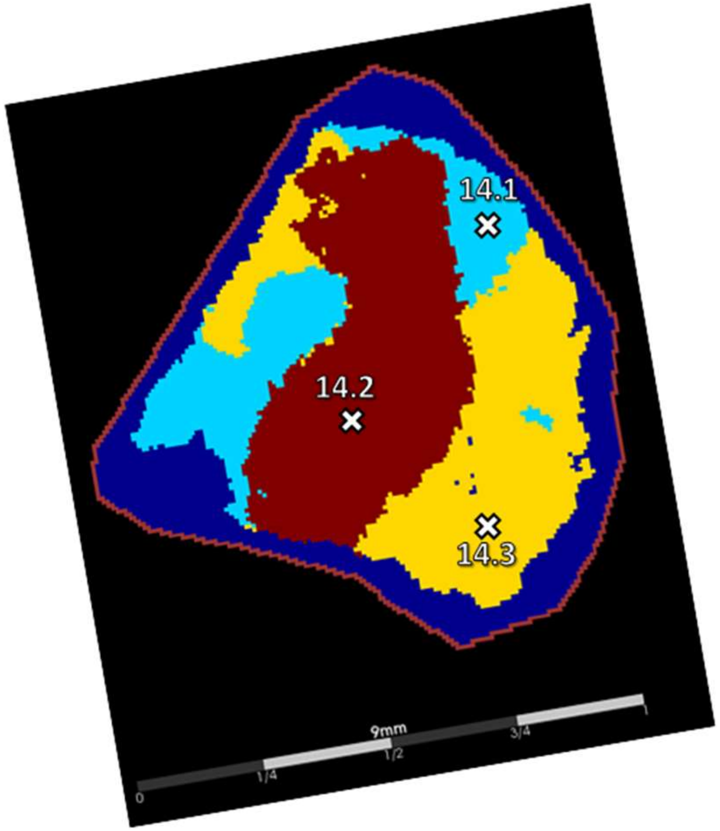
14.1
14.2
14.3
9mm
0
1/4
1/2
3/4
1

12

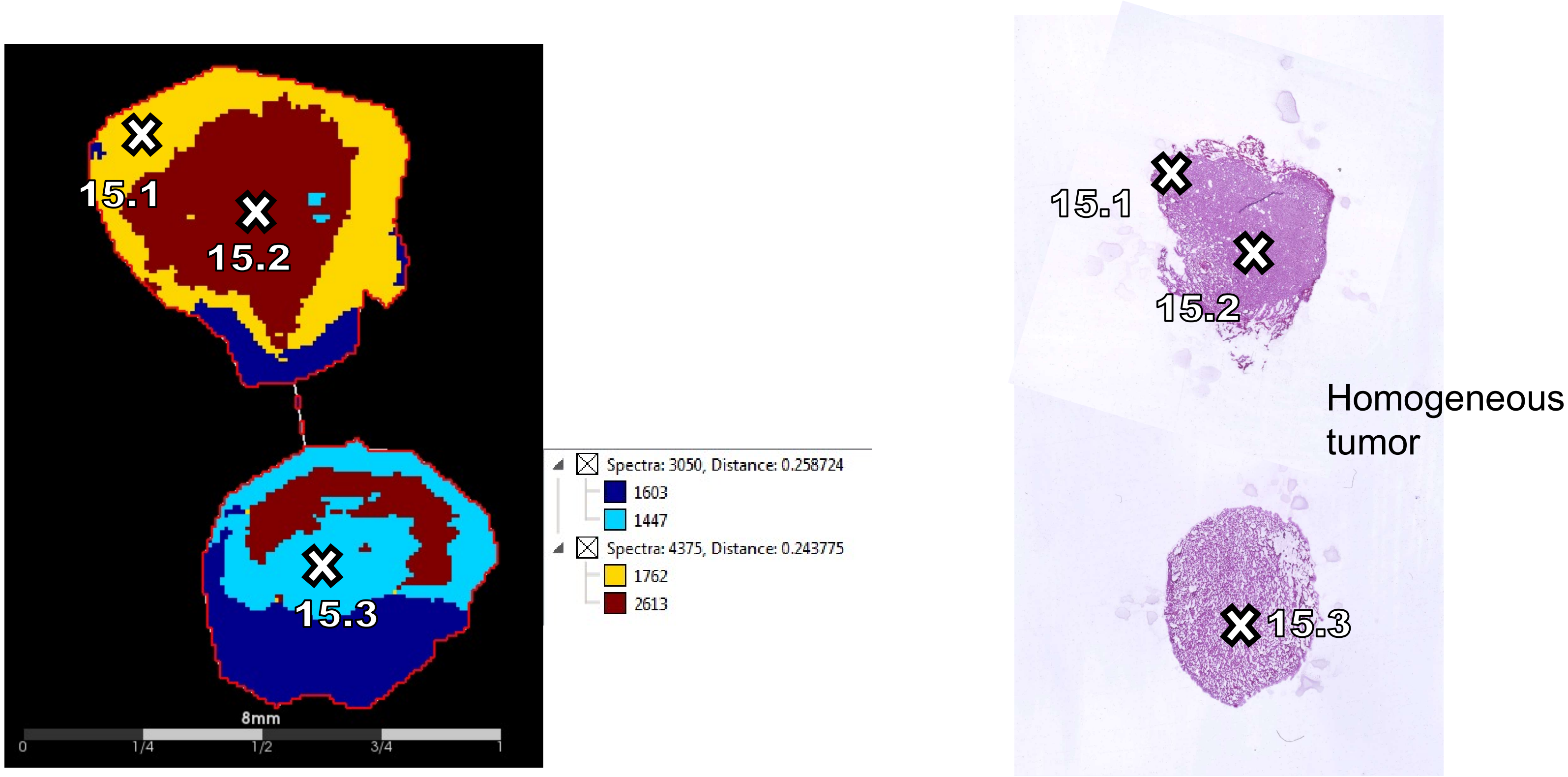
15.1
15.2
15.3
8mm
0
1/4
1/2
3/4
1
Spectra: 3050, Distance: 0.258724
1603
1447
Spectra: 4375, Distance: 0.243775
1762
2613
15.1
15.2
15.3
Homogeneous tumor

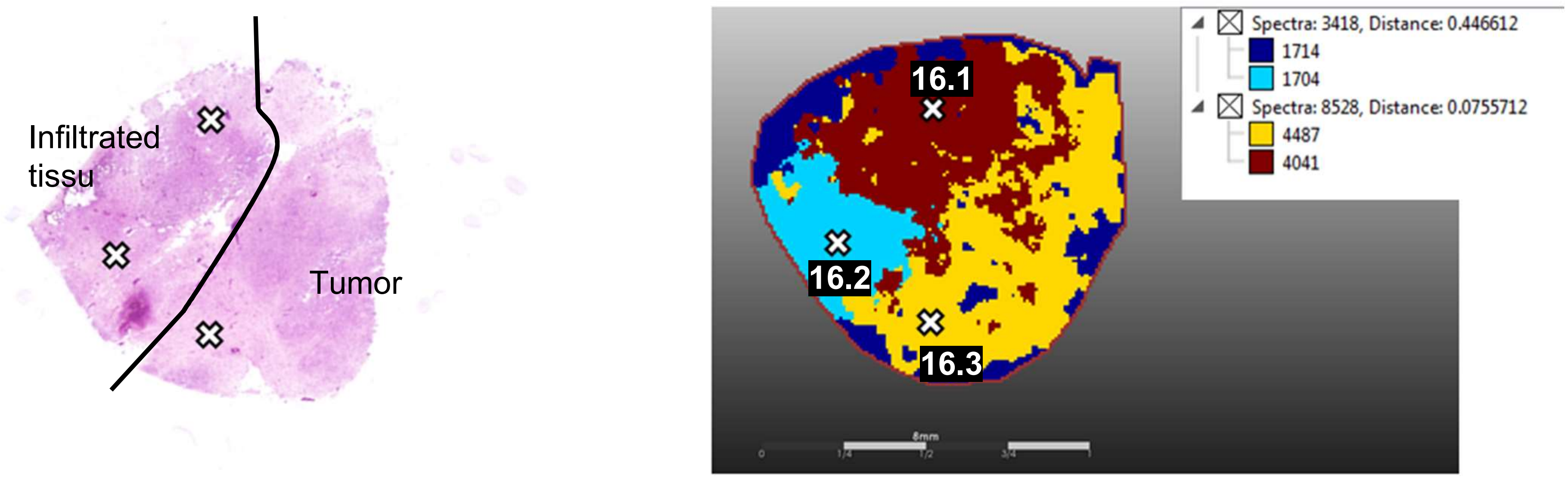
Infiltrated tissu
Tumor
Spectra: 3418, Distance: 0.446612
1714
1704
Spectra: 8528, Distance: 0.0755712
4487
4041
16.1
16.2
16.3

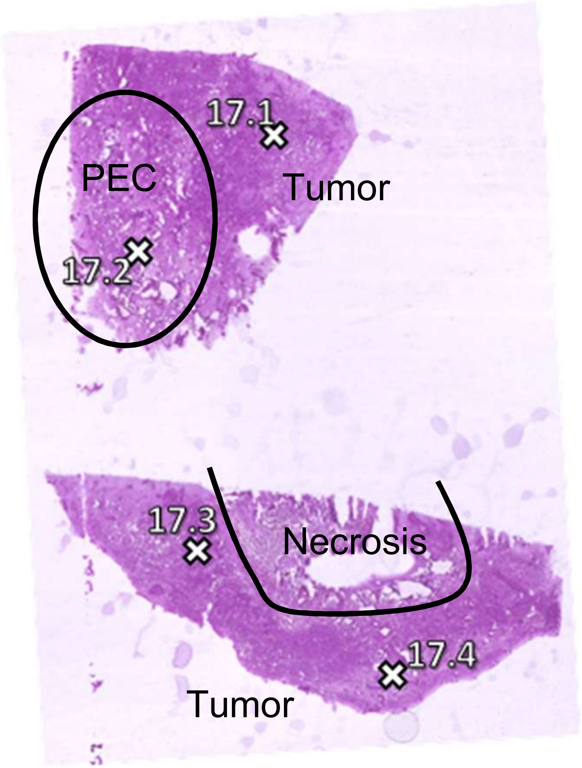
PEC
17.1
17.2
Tumor
17.3
Necrosis
17.4
Tumor

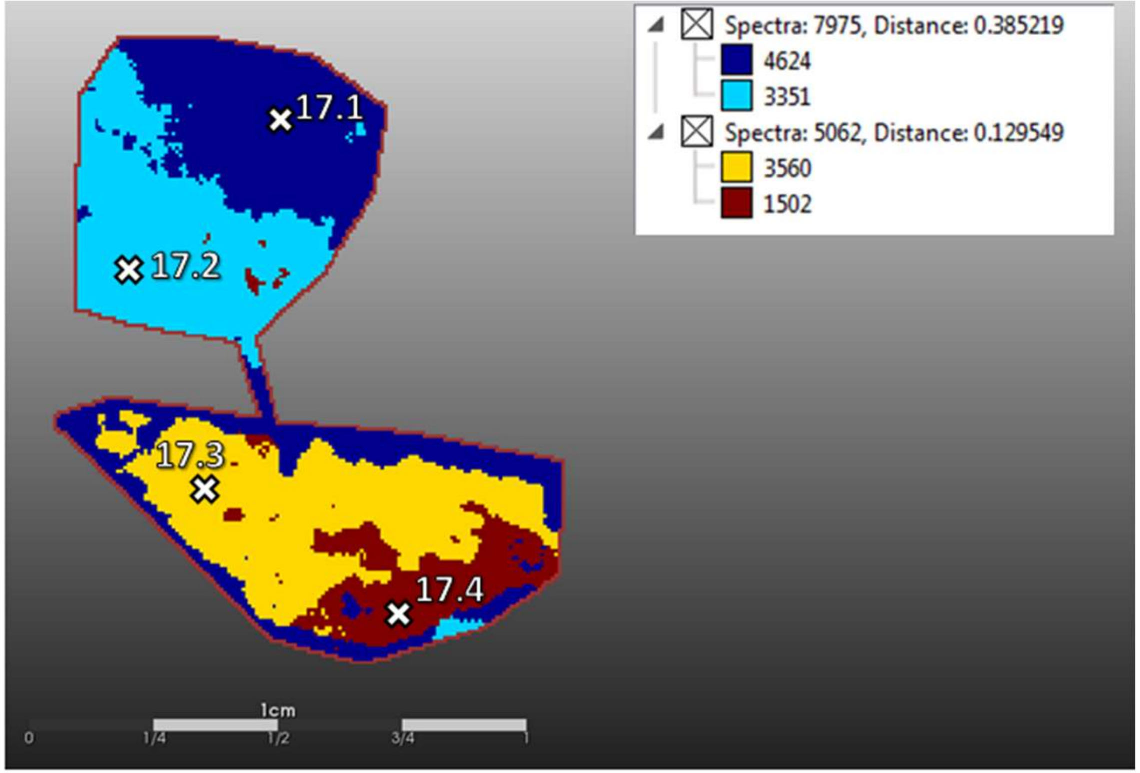
17.1
17.2
17.3
17.4
Spectra: 7975, Distance: 0.385219
4624
3351
Spectra: 5062, Distance: 0.129549
3560
1502
1cm
0
1/4
1/2
3/4
1

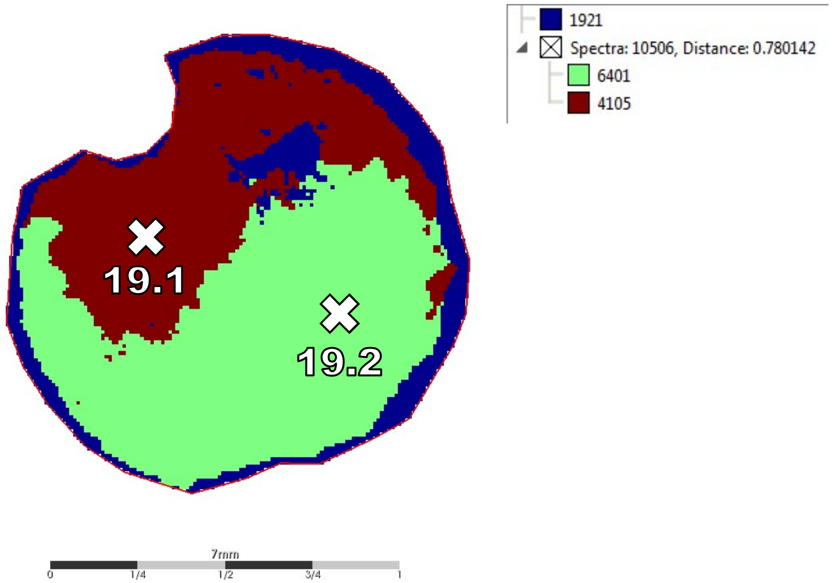
1921
Spectra: 10506, Distance: 0.780142
6401
4105
19.1
19.2
7mm
0
1/4
1/2
3/4
1

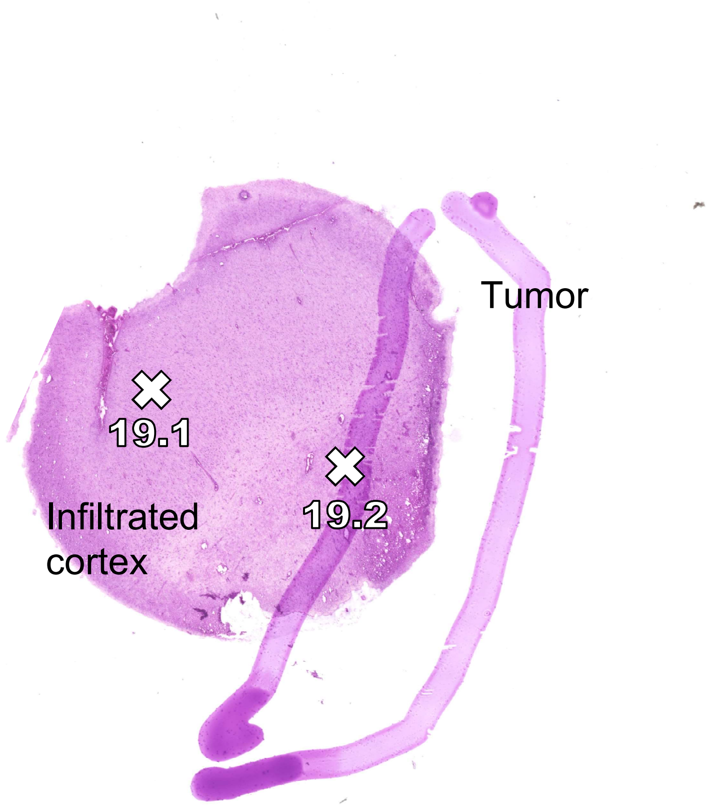
Tumor
19.1
19.2
Infiltrated cortex

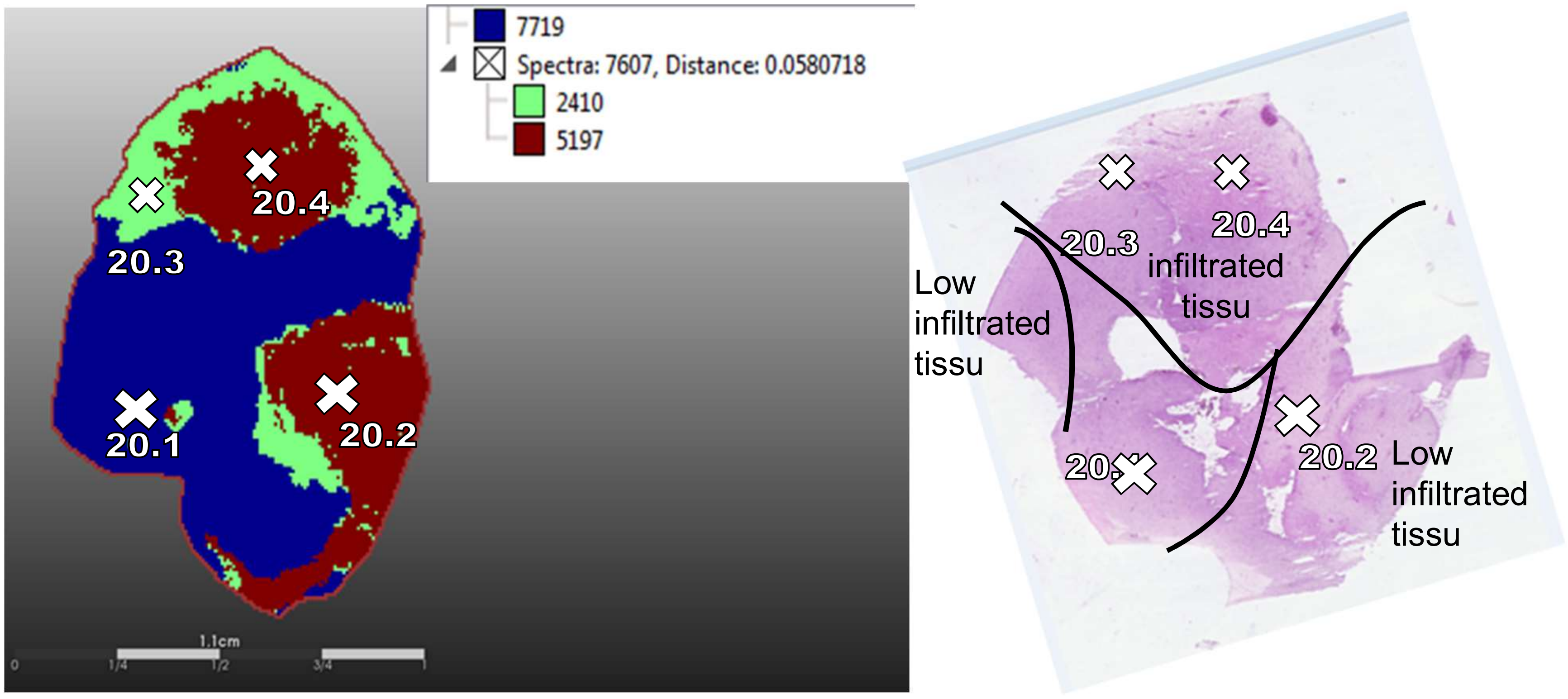
7719
Spectra: 7607, Distance: 0.0580718
2410
5197
20.4
20.3
20.1
20.2
1.1cm
0
1/4
1/2
3/4
1
20.3
20.4
infiltrated
tissu
Low
infiltrated
tissu
20.2
Low
infiltrated
tissu

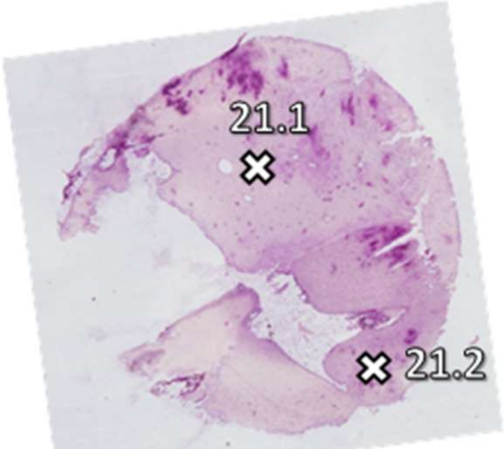


Homogeneous low
infiltrated tissu

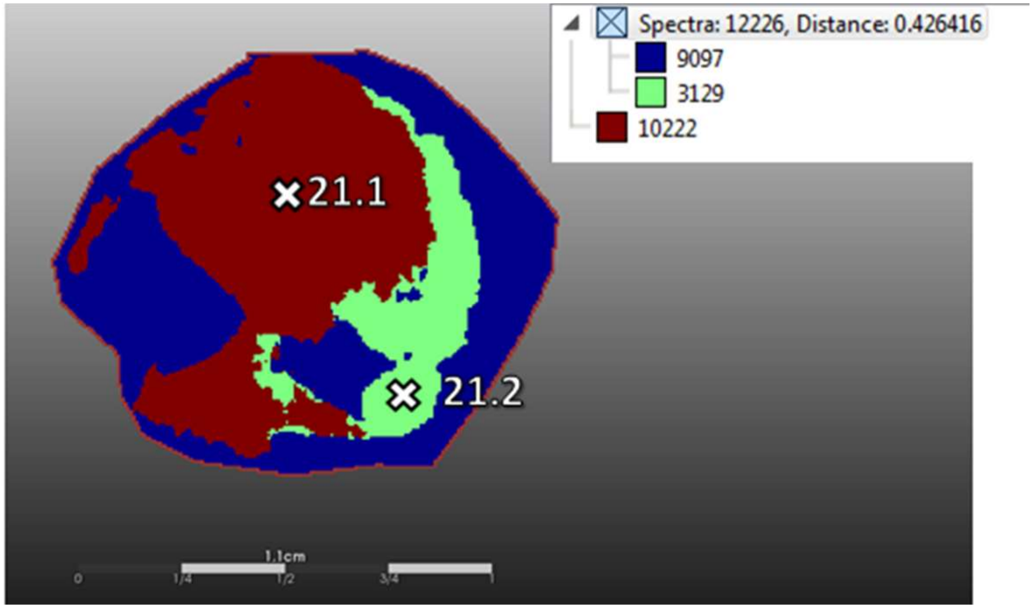

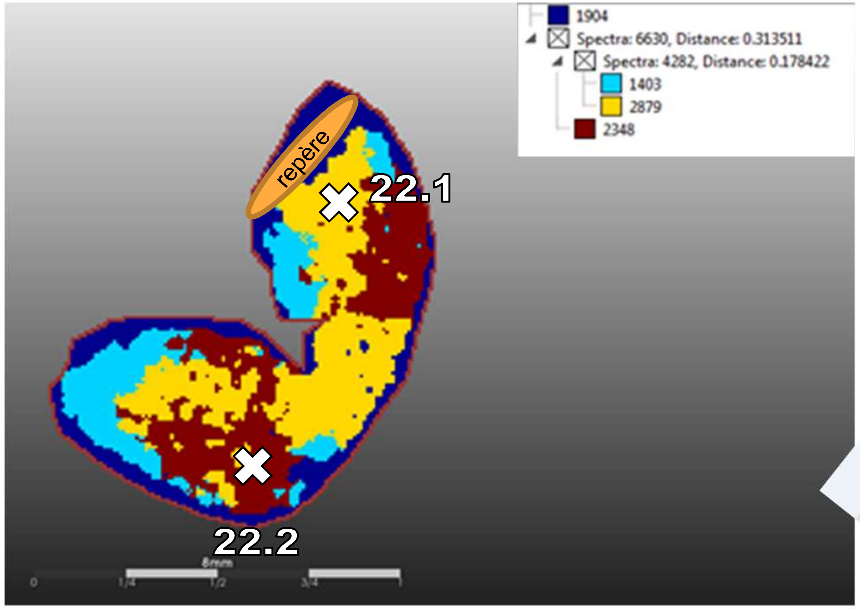
1904
Spectra: 6630, Distance: 0.313511
Spectra: 4282, Distance: 0.178422
1403
2879
2348
repère
22.1
22.2

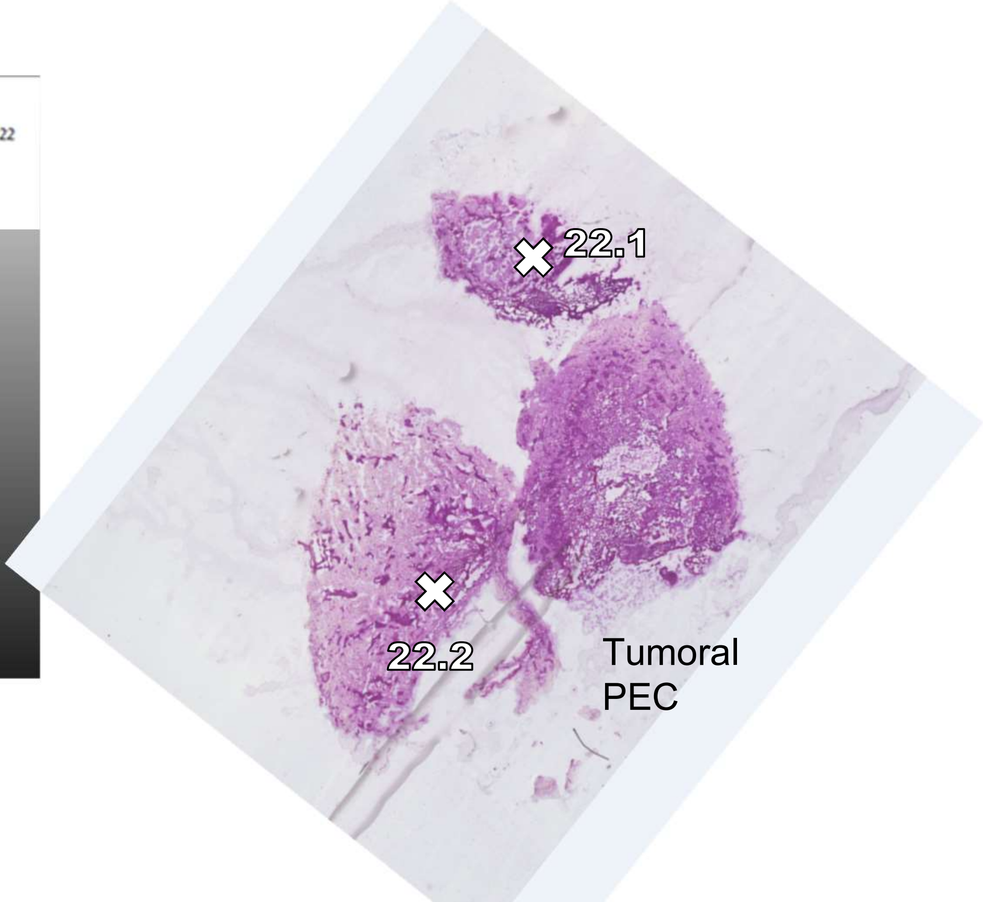
22.1
22.2
Tumoral
PEC

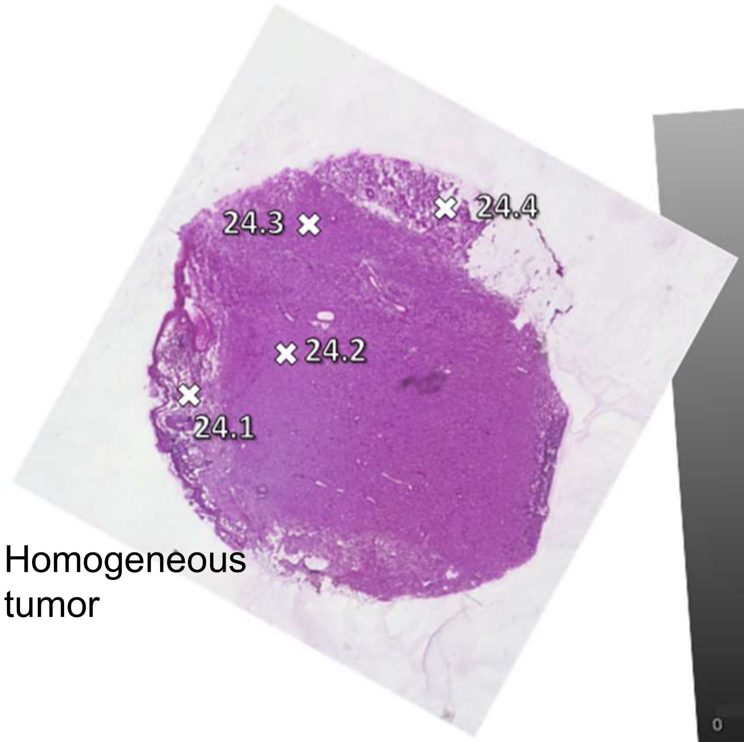


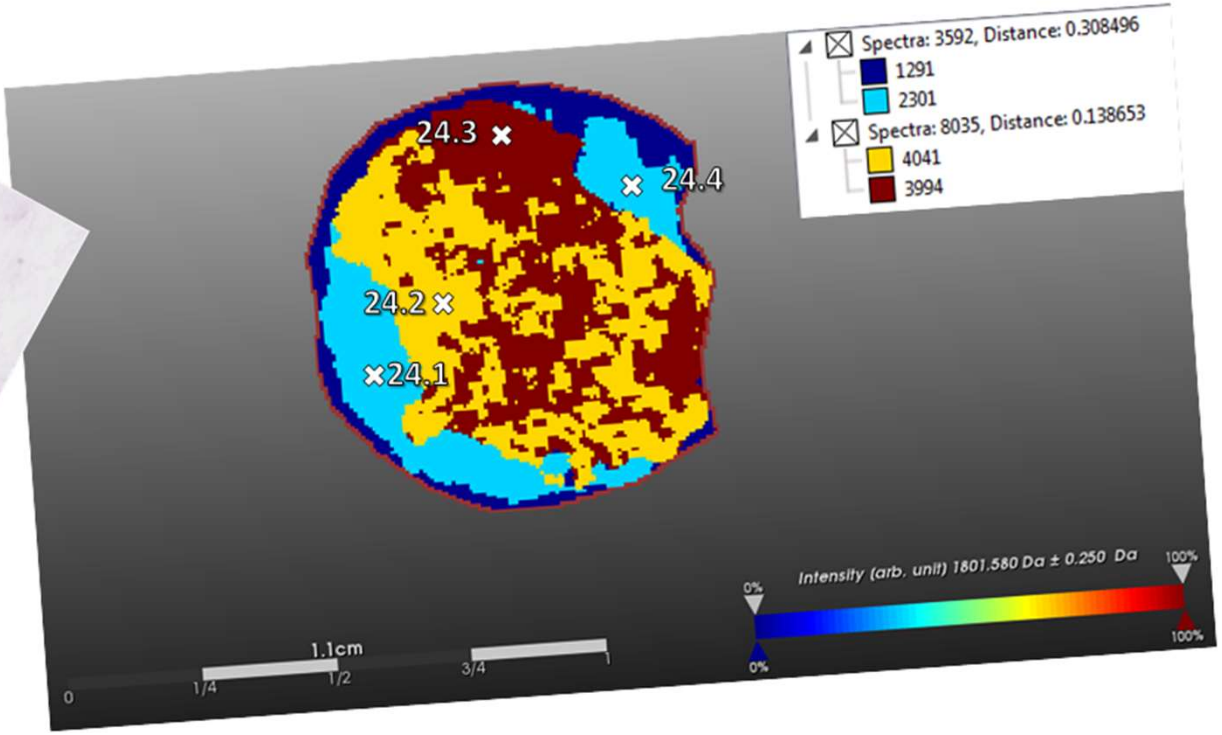


Homogeneous tumor

20

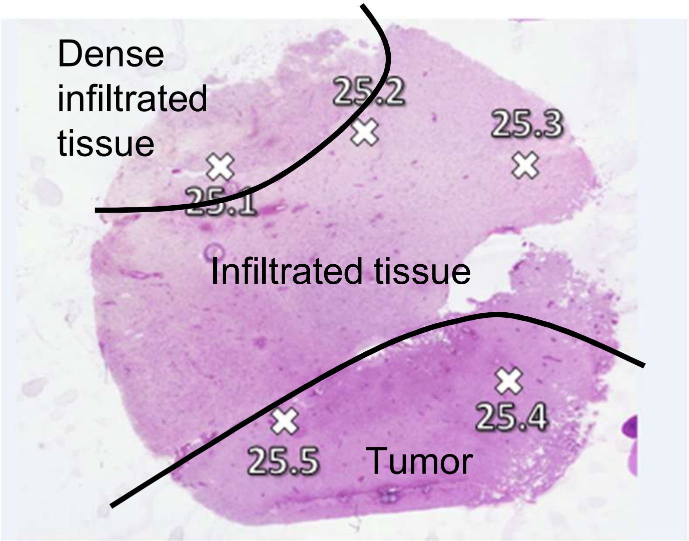
Dense
infiltrated
tissue
25.1
25.2
25.3
Infiltrated tissue
25.4
25.5
Tumor

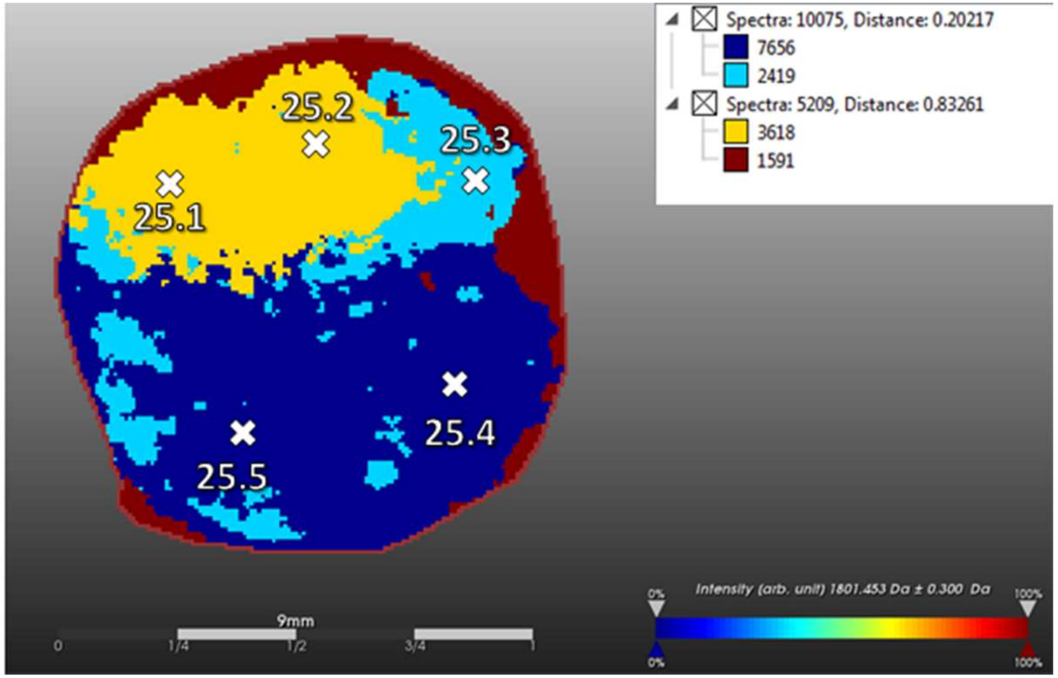
Spectra: 10075, Distance: 0.20217
7656
2419
Spectra: 5209, Distance: 0.83261
3618
1591
25.1
25.2
25.3
25.4
25.5
9mm
Intensity (arb. unit) 1801.453 Da ± 0.300 Da

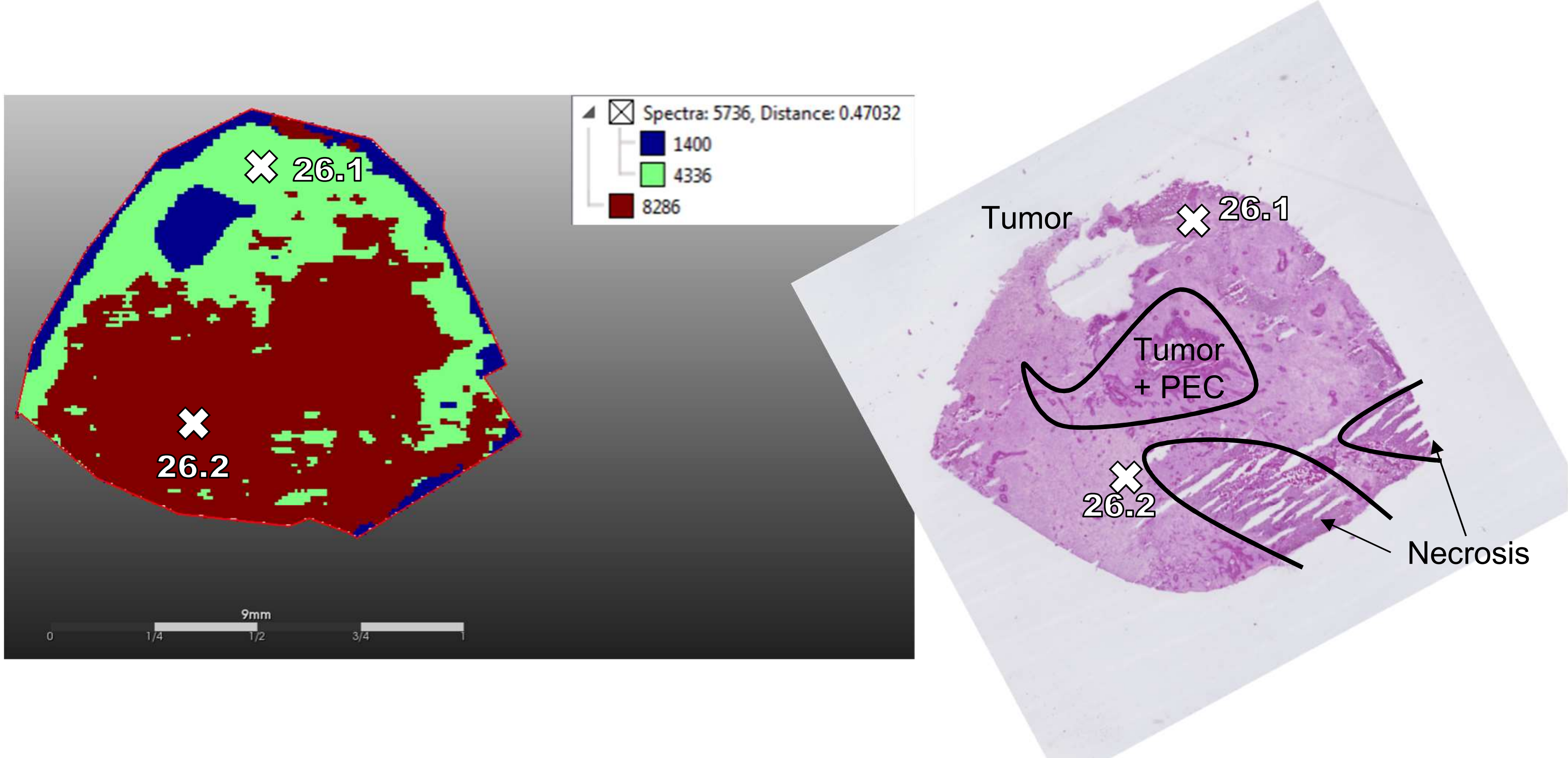
Spectra: 5736, Distance: 0.47032
1400
4336
8286
26.1
26.2
9mm
0
1/4
1/2
3/4
1
Tumor
26.1
Tumor
+ PEC
26.2
Necrosis

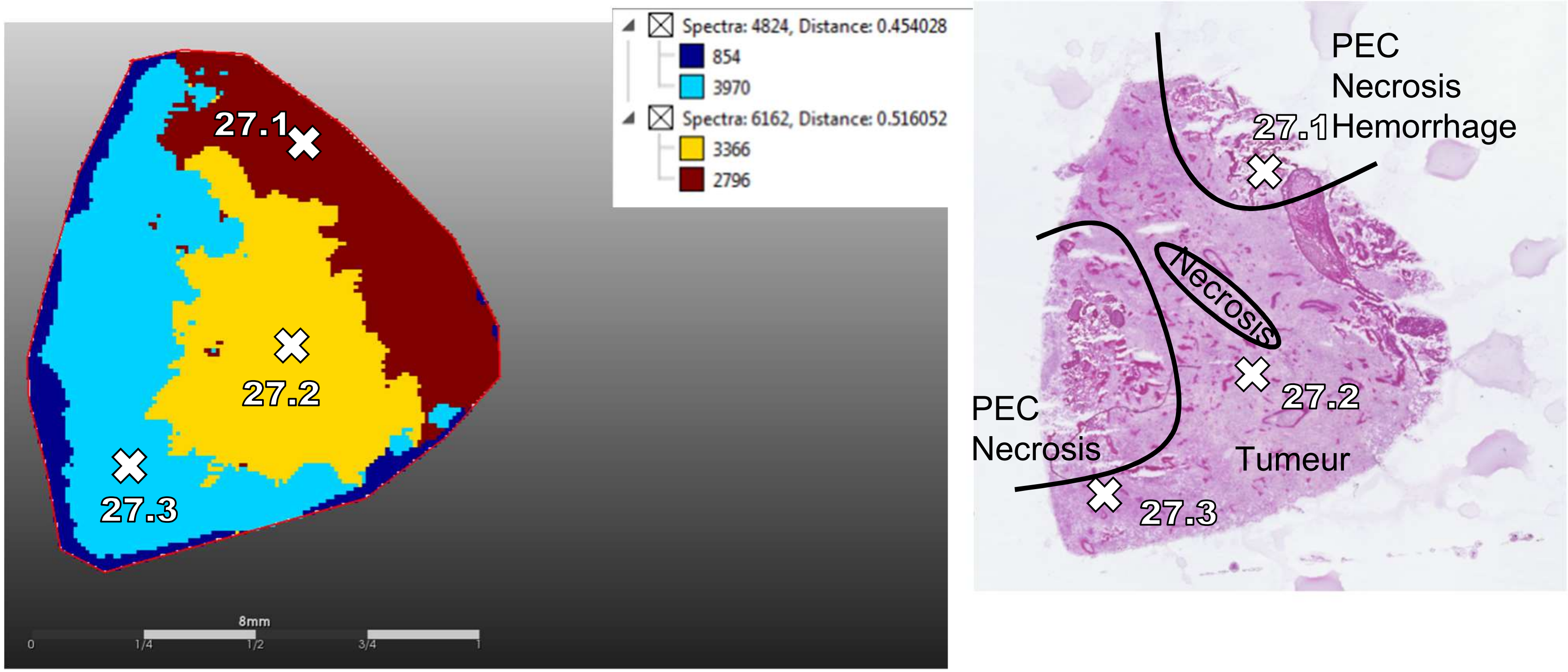
Spectra: 4824, Distance: 0.454028
854
3970
Spectra: 6162, Distance: 0.516052
3366
2796
27.1
27.2
27.3
8mm
0
1/4
1/2
3/4
1
PEC
Necrosis
Hemorrhage
Necrosis
PEC
Necrosis
Tumeur
27.1
27.2
27.3

23

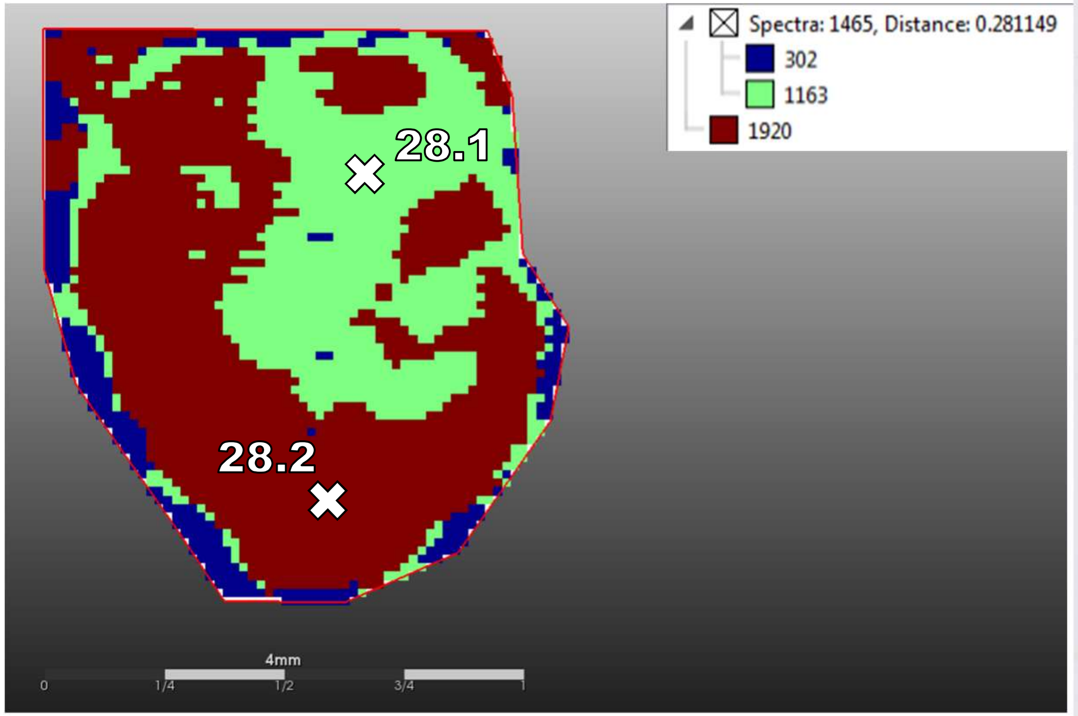
Spectra: 1465, Distance: 0.281149
302
1163
1920
28.1
28.2
4mm
0
1/4
1/2
3/4
1

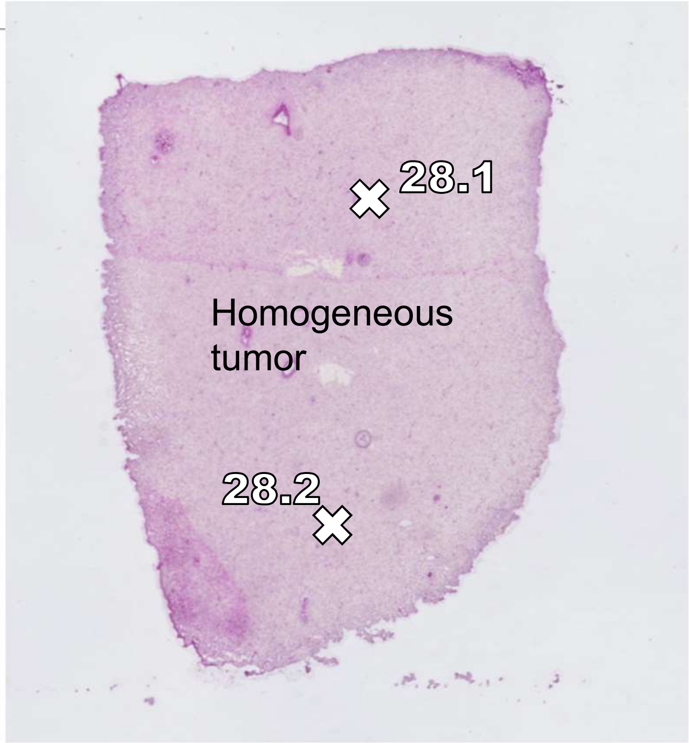
28.1
Homogeneous
tumor
28.2

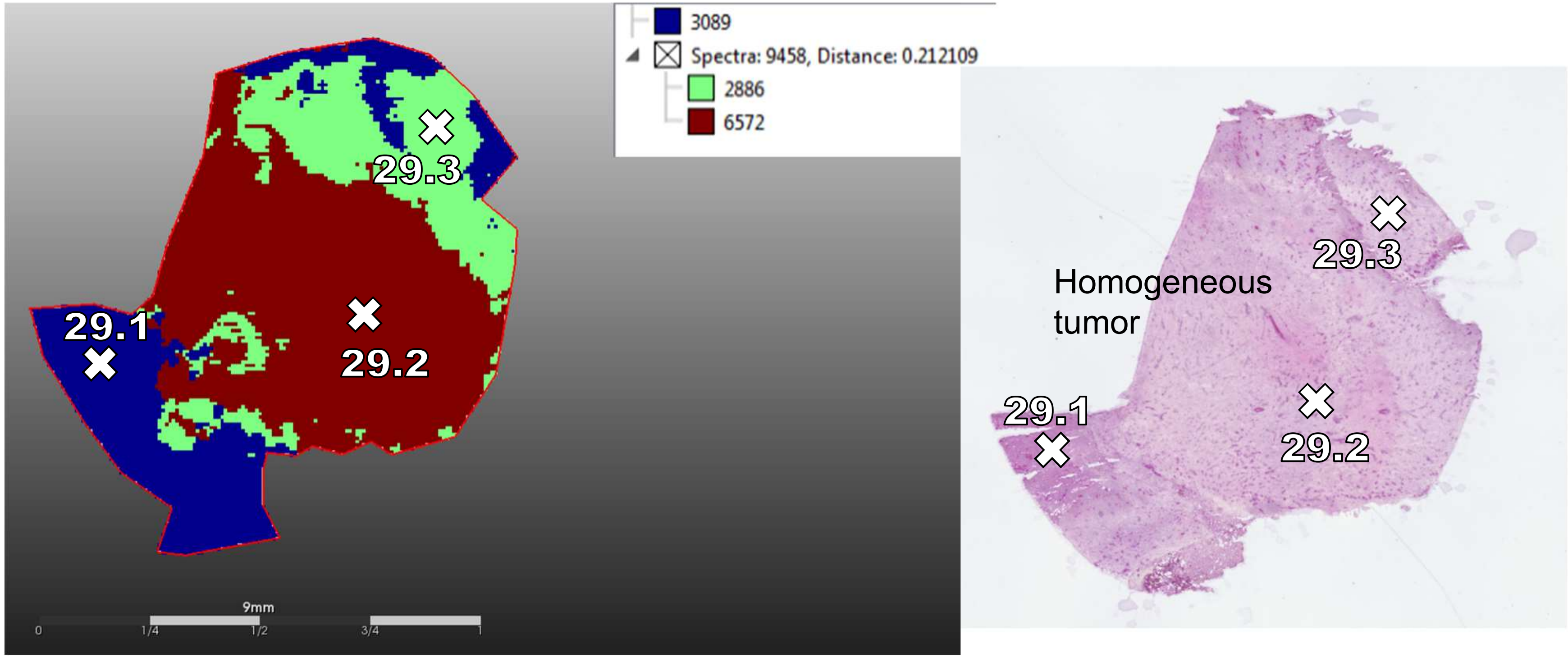
3089
Spectra: 9458, Distance: 0.212109
2886
6572
29.1
29.2
29.3
9mm
0
1/4
1/2
3/4
1
Homogeneous
tumor
29.1
29.2
29.3

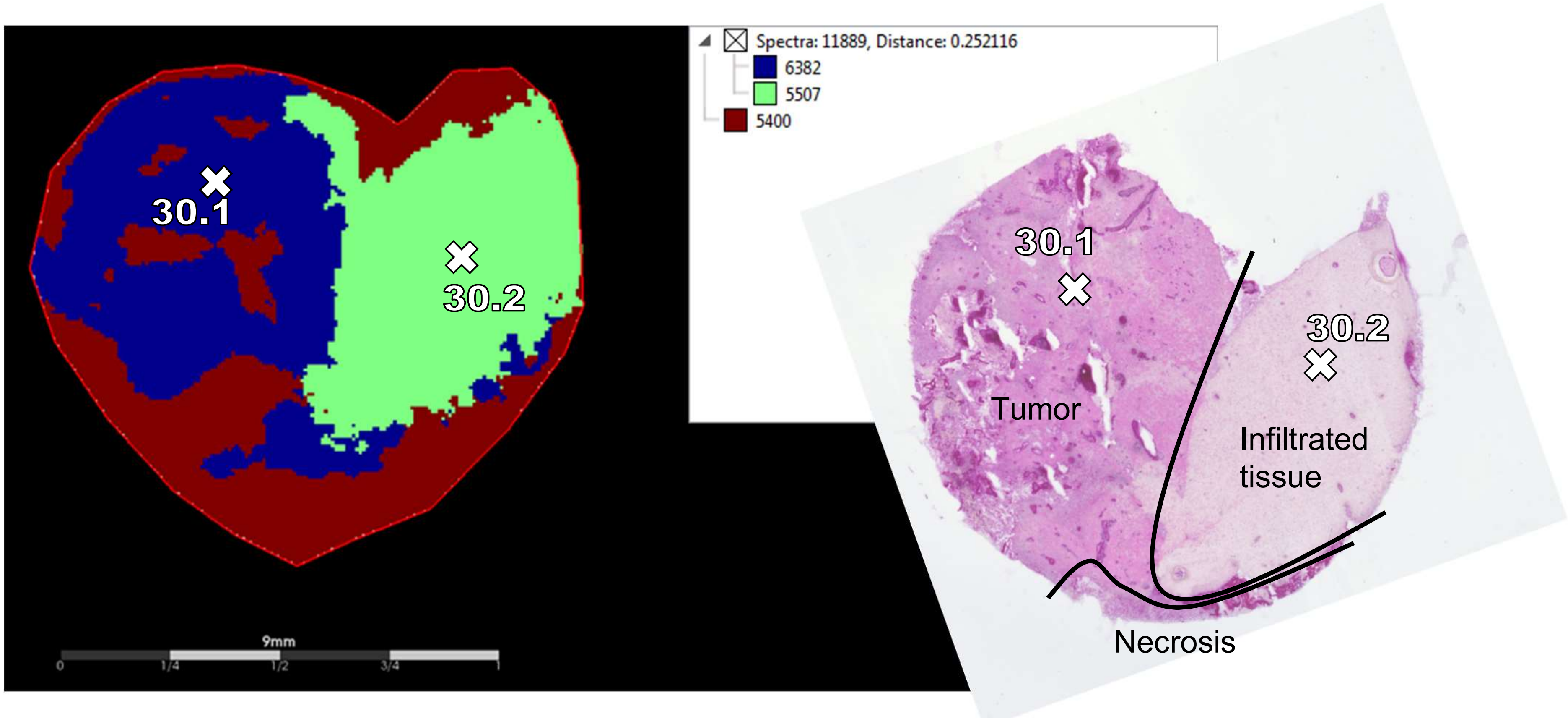
Spectra: 11889, Distance: 0.252116
6382
5507
5400
30.1
30.2
9mm
0
1/4
1/2
3/4
1
30.1
30.2
Tumor
Infiltrated tissue
Necrosis

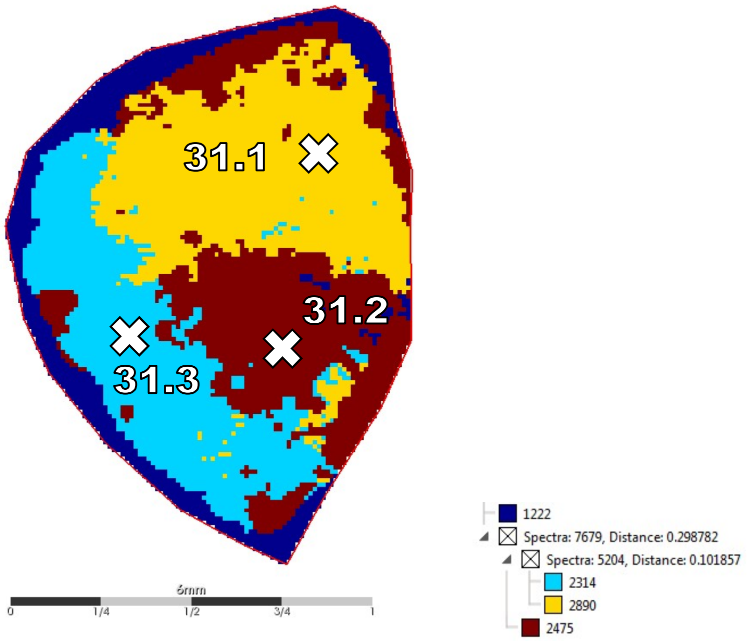

31.1
31.2
31.3
6mm
0
1/4
1/2
3/4
1
1222
Spectra: 7679, Distance: 0.298782
Spectra: 5204, Distance: 0.101857
2314
2890
2475


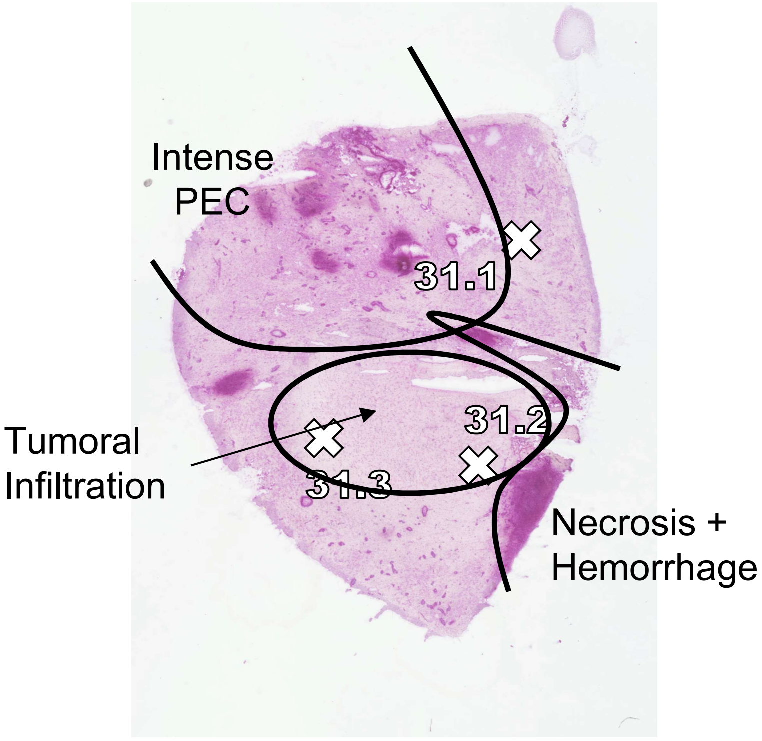

Intense PEC
31.1
31.2
31.3
Tumoral Infiltration
Necrosis + Hemorrhage

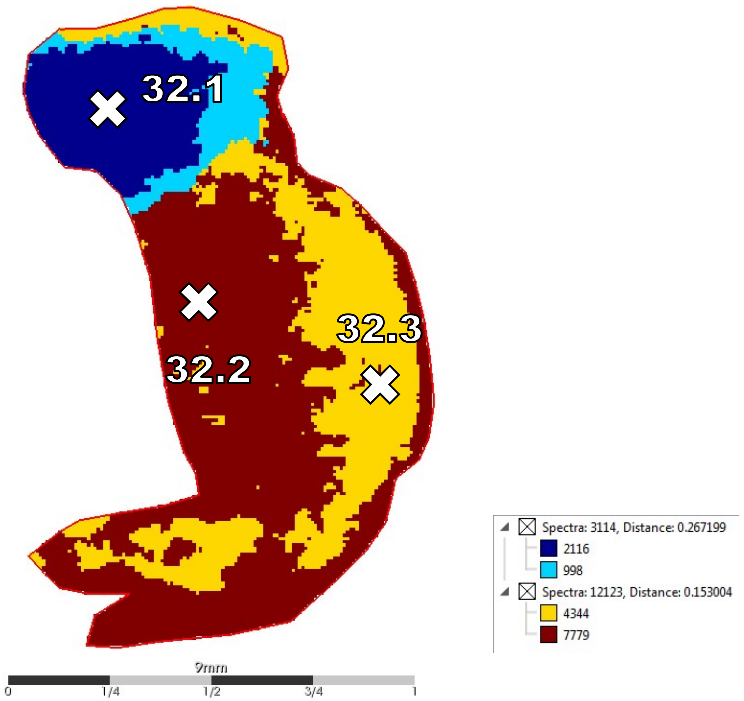
32.1
32.2
32.3
Spectra: 3114, Distance: 0.267199
2116
998
Spectra: 12123, Distance: 0.153004
4344
7779
9mm
0
1/4
1/2
3/4
1

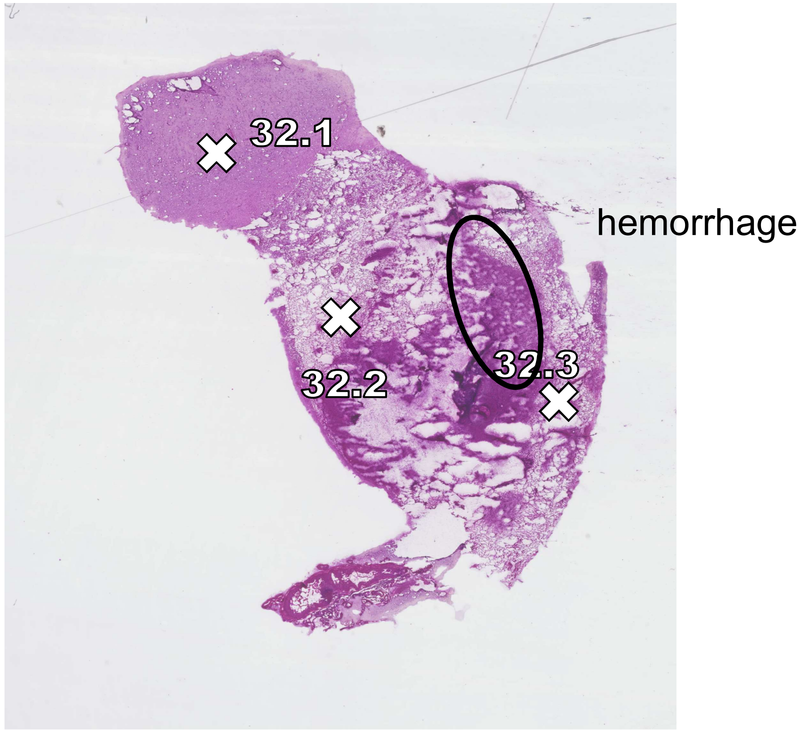
32.1
32.2
32.3
hemorrhage

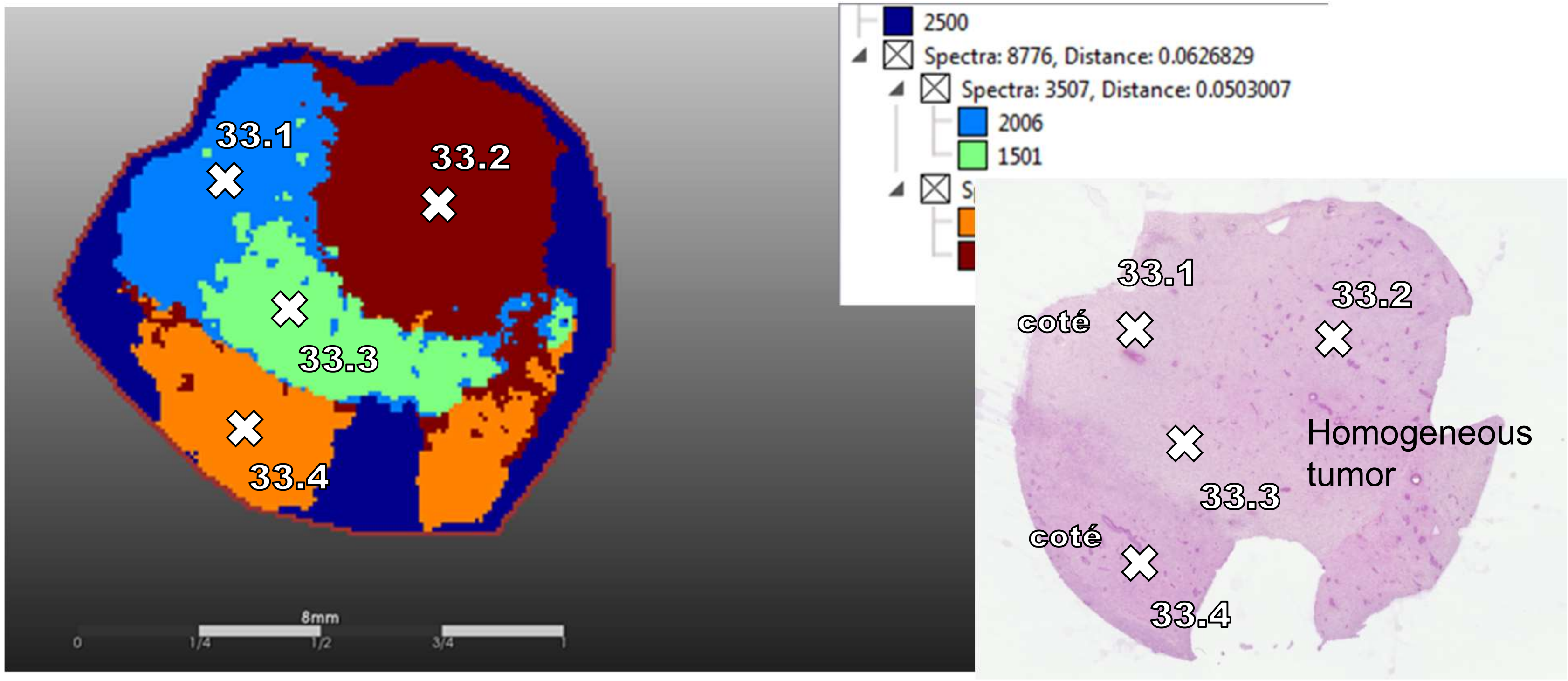

33.1
33.2
33.3
33.4
8mm
0
1/4
1/2
3/4
1
2500
Spectra: 8776, Distance: 0.0626829
Spectra: 3507, Distance: 0.0503007
2006
1501
33.1
33.2
coté
Homogeneous tumor
33.3
coté
33.4

29

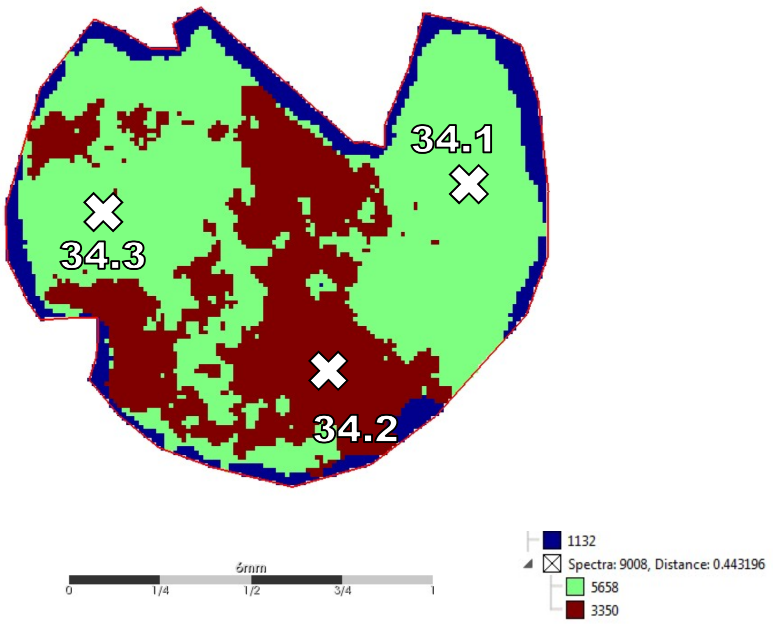
34.1
34.3
34.2
6mm
0
1/4
1/2
3/4
1
1132
Spectra: 9008, Distance: 0.443196
5658
3350

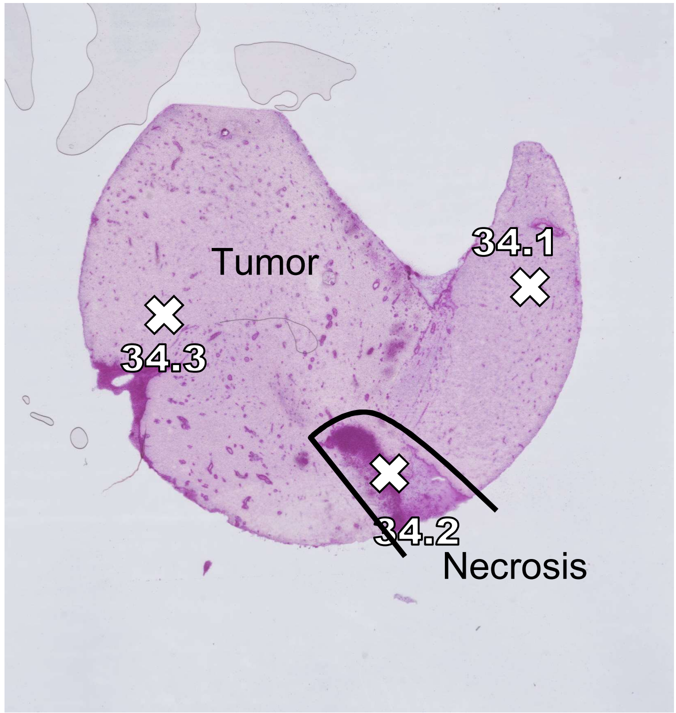
Tumor
34.1
34.3
34.2
Necrosis

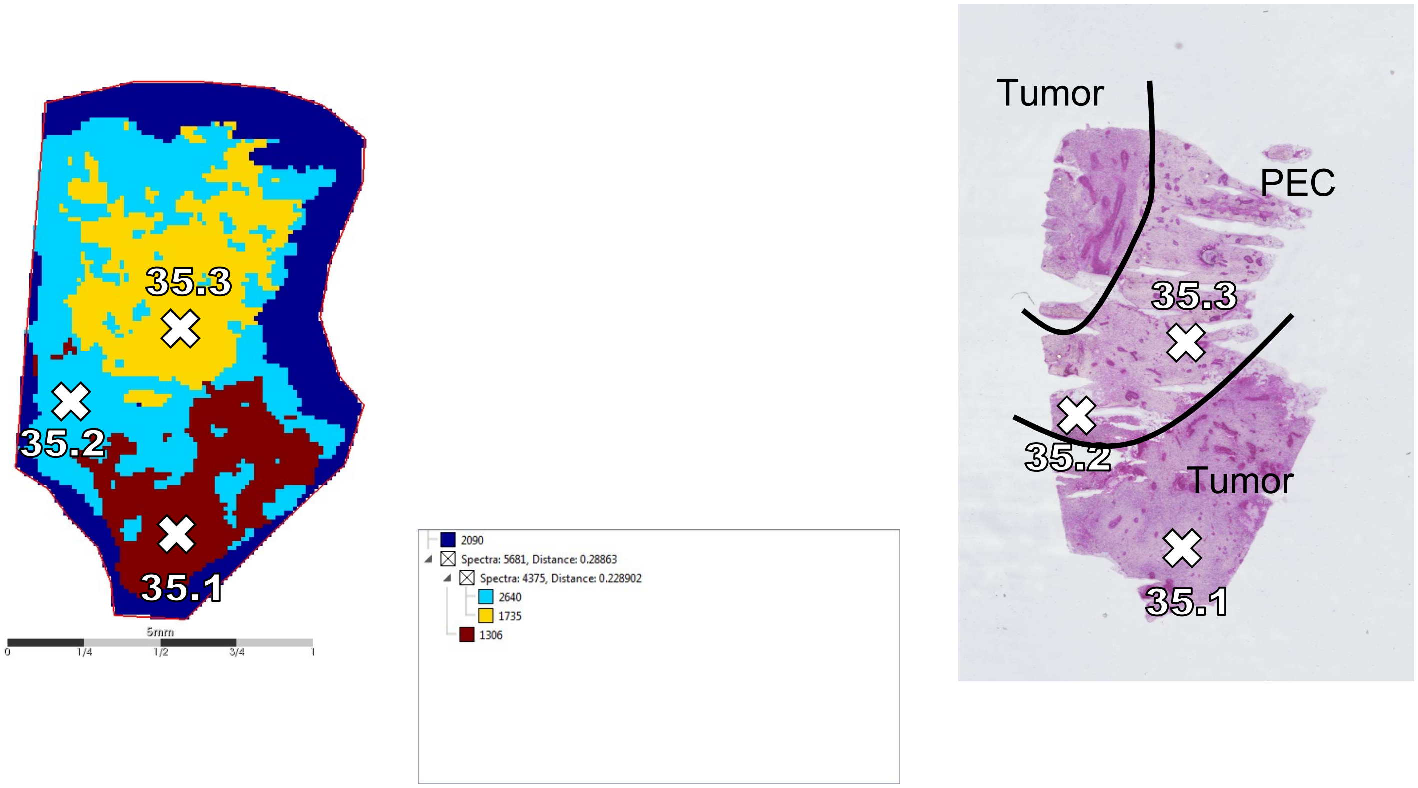
35.3
35.2
35.1
5mm
0
1/4
1/2
3/4
1
2090
Spectra: 5681, Distance: 0.28863
Spectra: 4375, Distance: 0.228902
2640
1735
1306
Tumor
PEC
35.3
35.2
Tumor
35.1

31

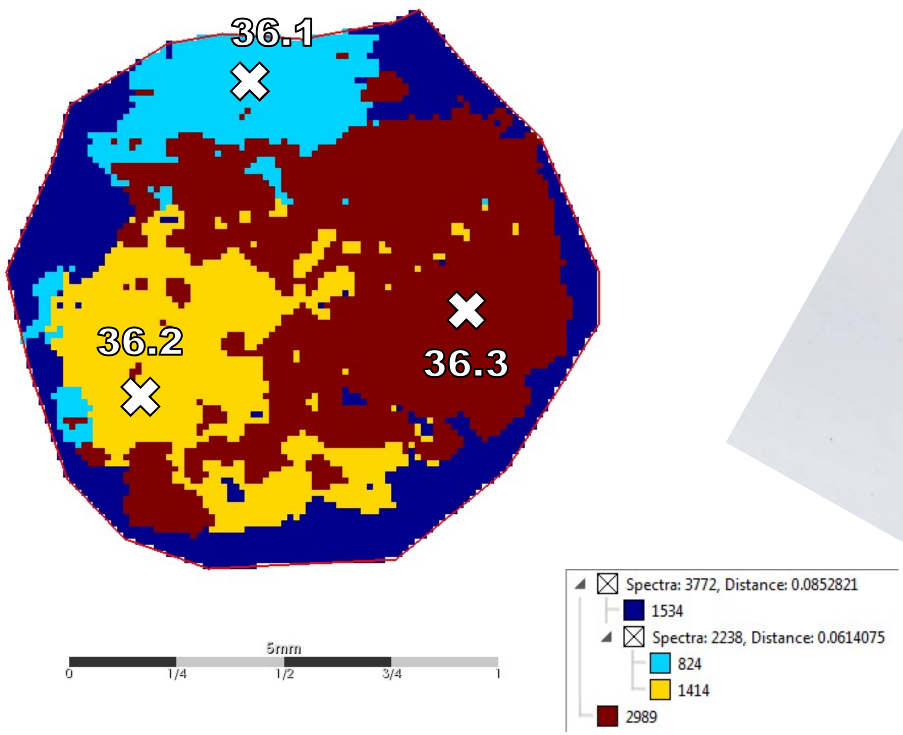
36.1
36.2
36.3
5mm
0
1/4
1/2
3/4
1
Spectra: 3772, Distance: 0.0852821
1534
Spectra: 2238, Distance: 0.0614075
824
1414
2989

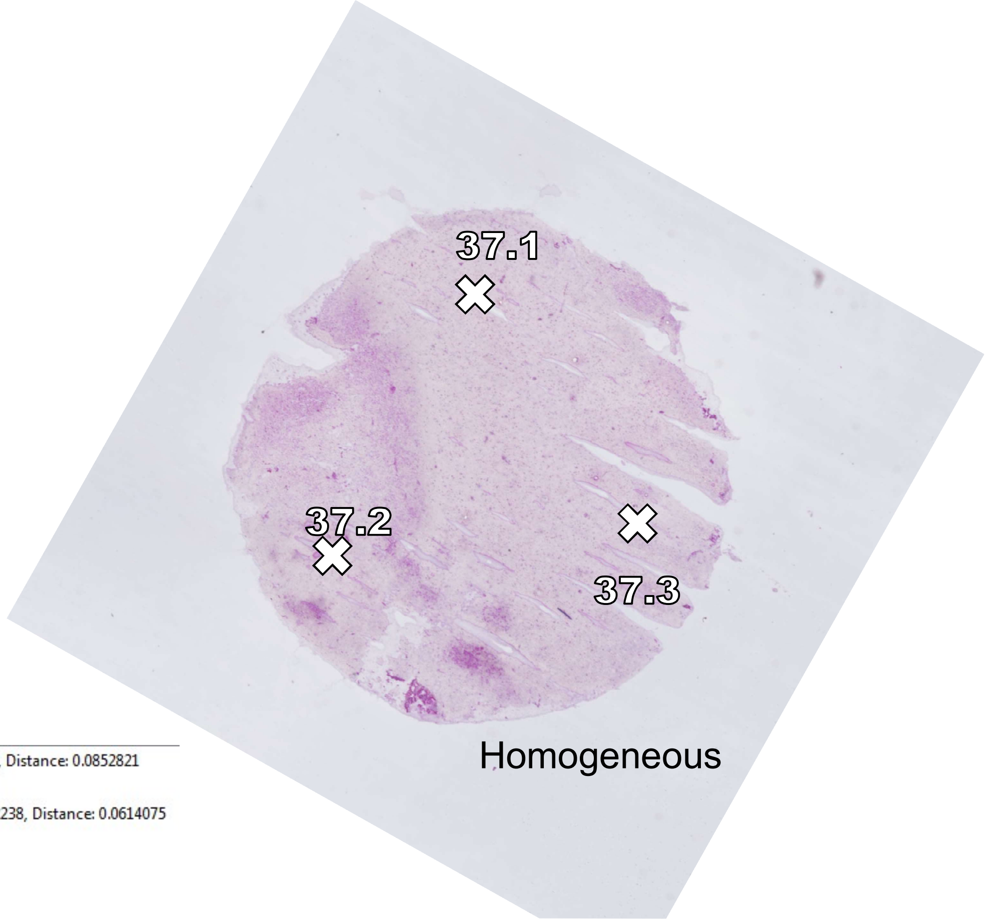
37.1
37.2
37.3
Homogeneous

32

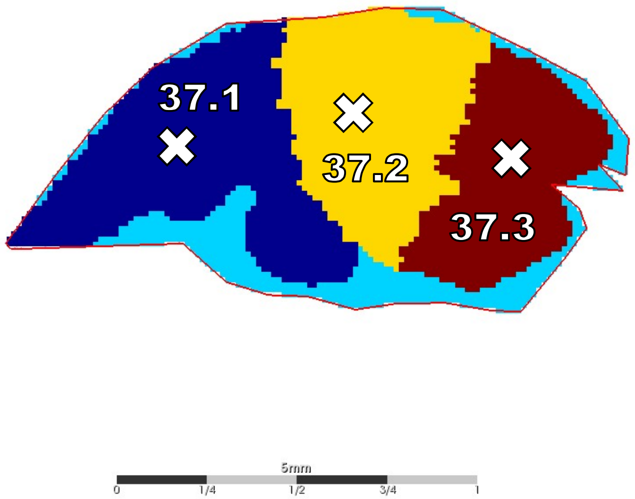
37.1
37.2
37.3
5mm
0
1/4
1/2
3/4
1

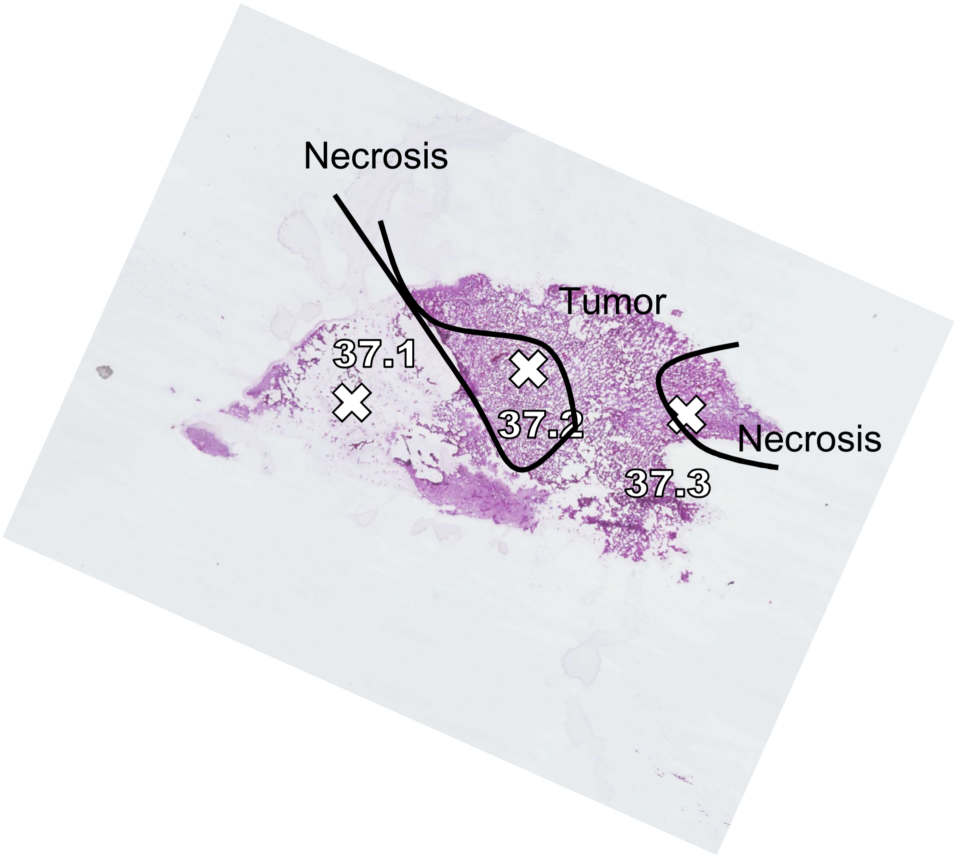
Necrosis
Tumor
37.1
37.2
37.3
Necrosis

33

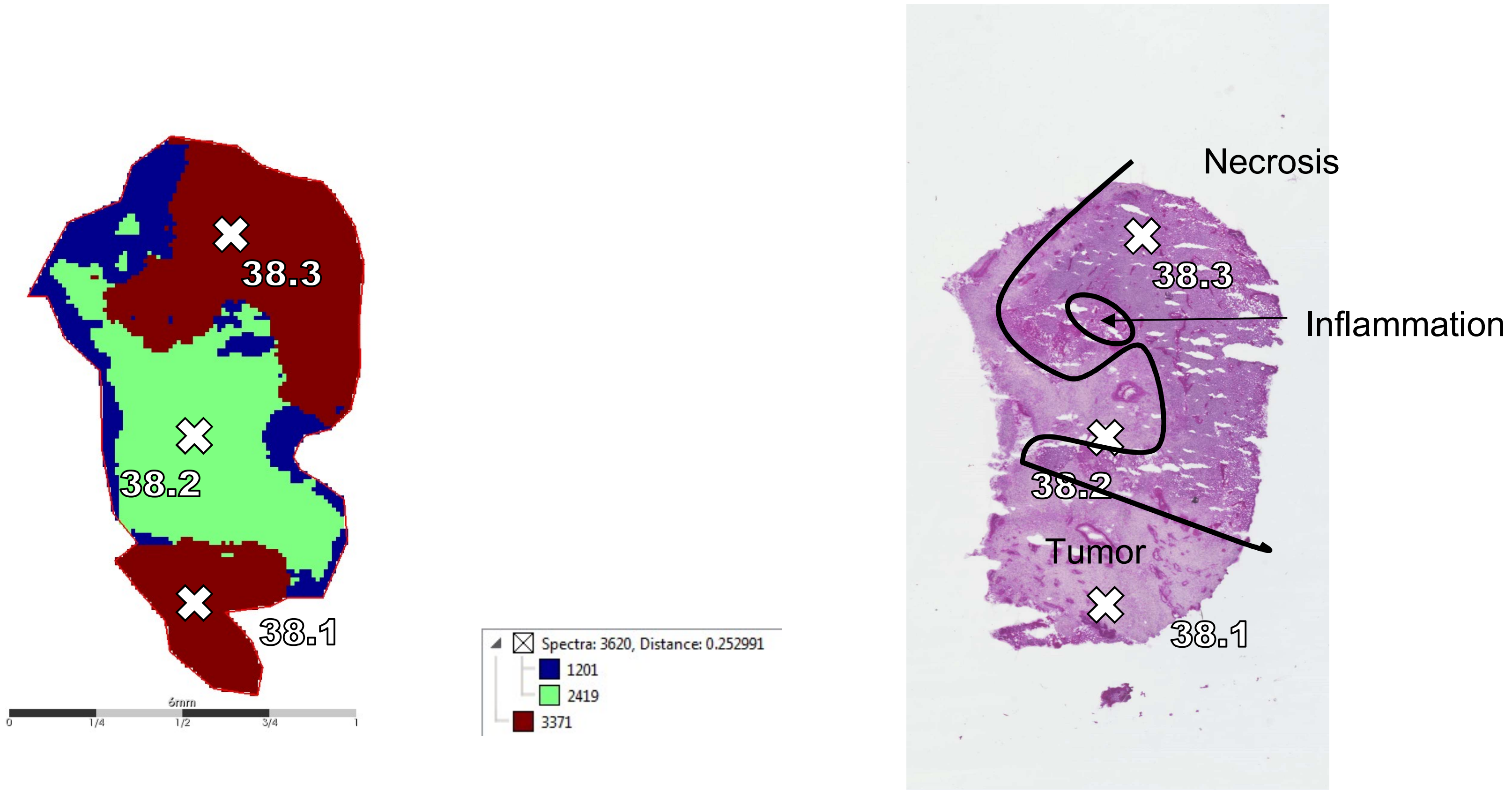
38.3
38.2
38.1
Spectra: 3620, Distance: 0.252991
1201
2419
3371
Necrosis
Inflammation
Tumor

34

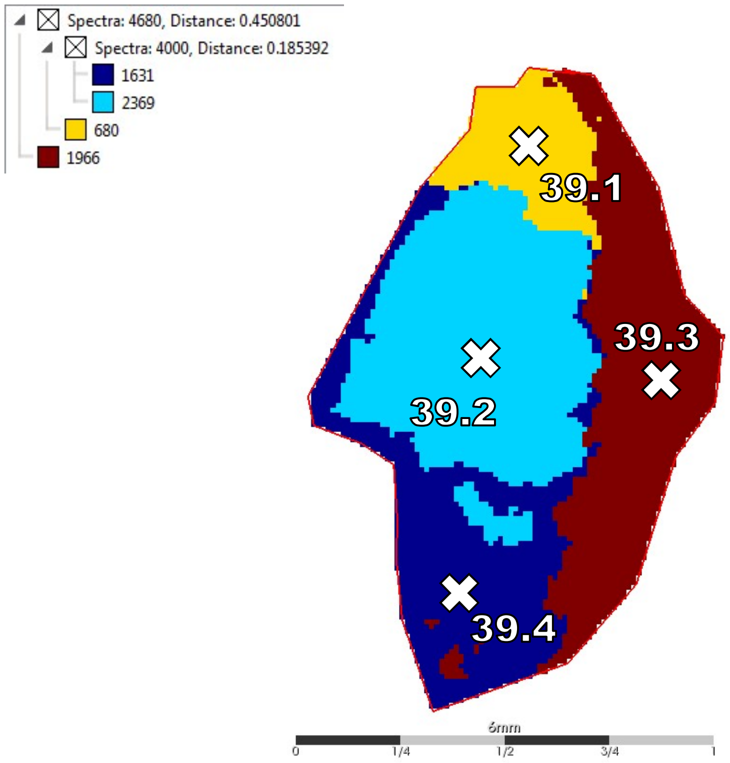
Spectra: 4680, Distance: 0.450801
Spectra: 4000, Distance: 0.185392
1631
2369
680
1966
39.1
39.2
39.3
39.4
6mm
0
1/4
1/2
3/4
1

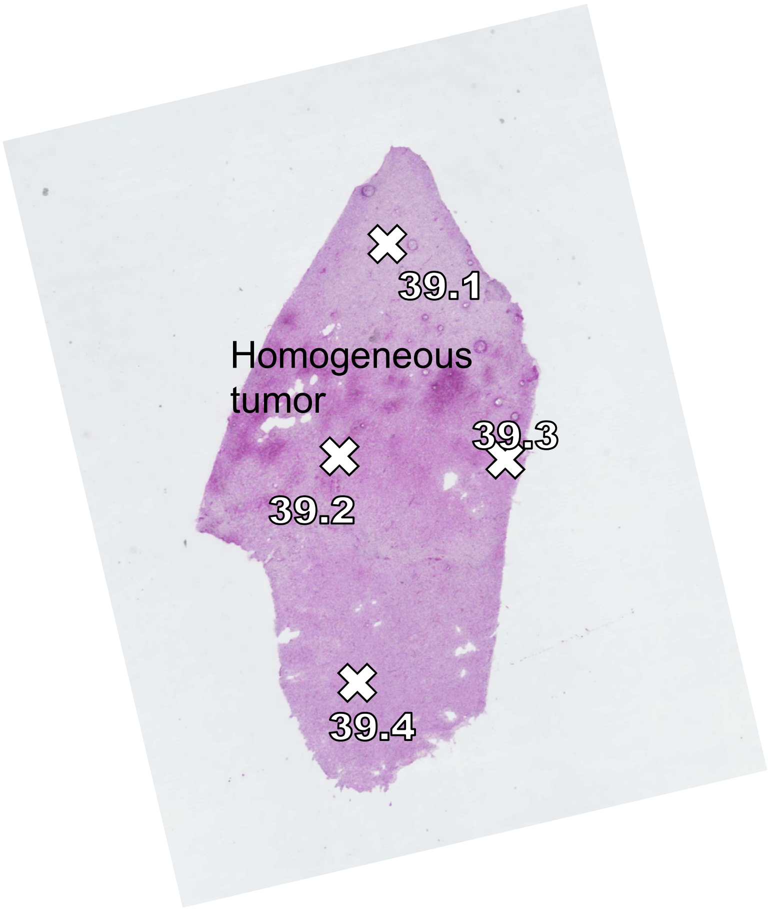
39.1
Homogeneous tumor
39.3
39.2
39.4

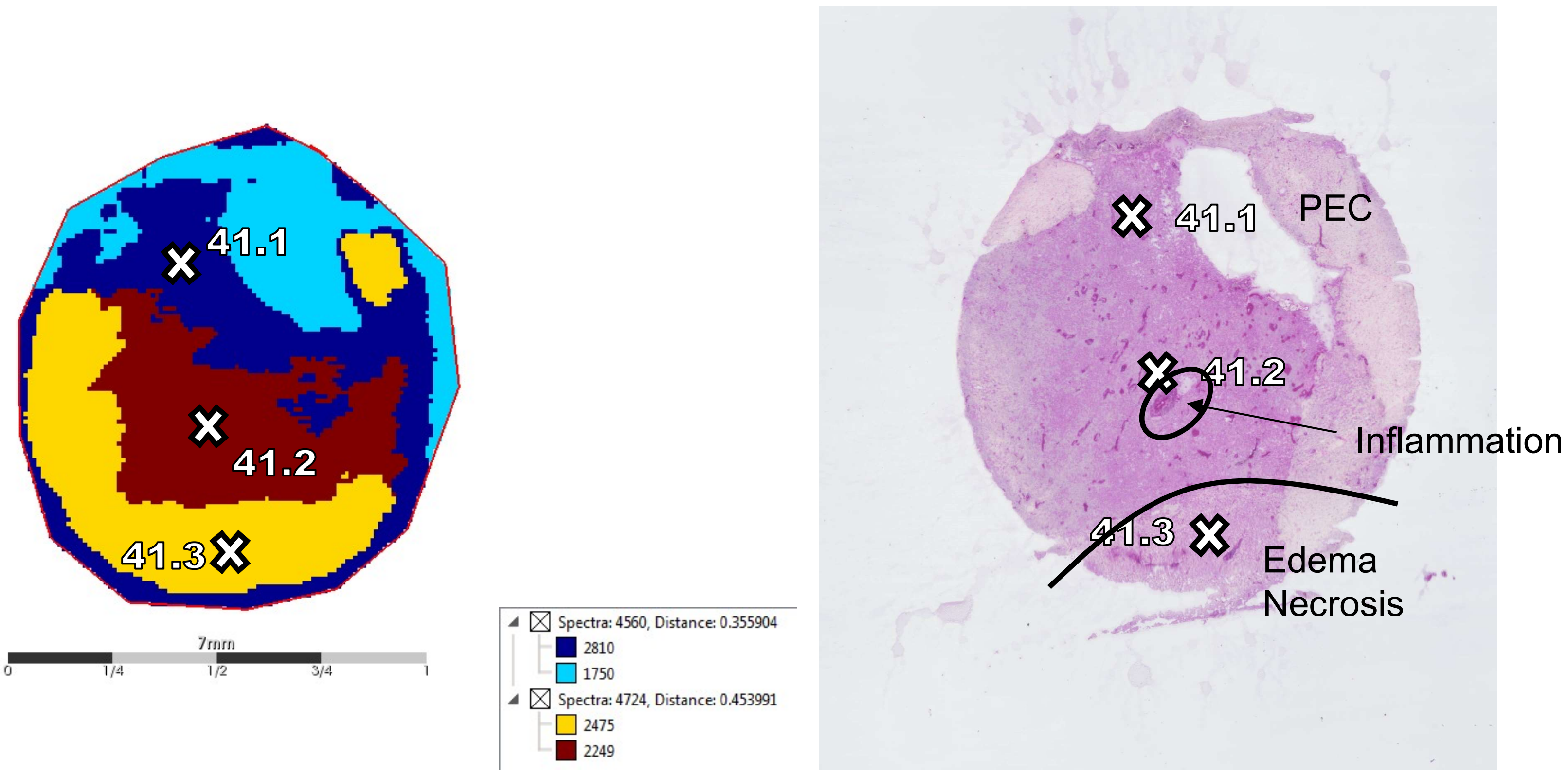
41.1
41.2
41.3
7mm
0
1/4
1/2
3/4
1
Spectra: 4560, Distance: 0.355904
2810
1750
Spectra: 4724, Distance: 0.453991
2475
2249
41.1
PEC
41.2
Inflammation
41.3
Edema
Necrosis

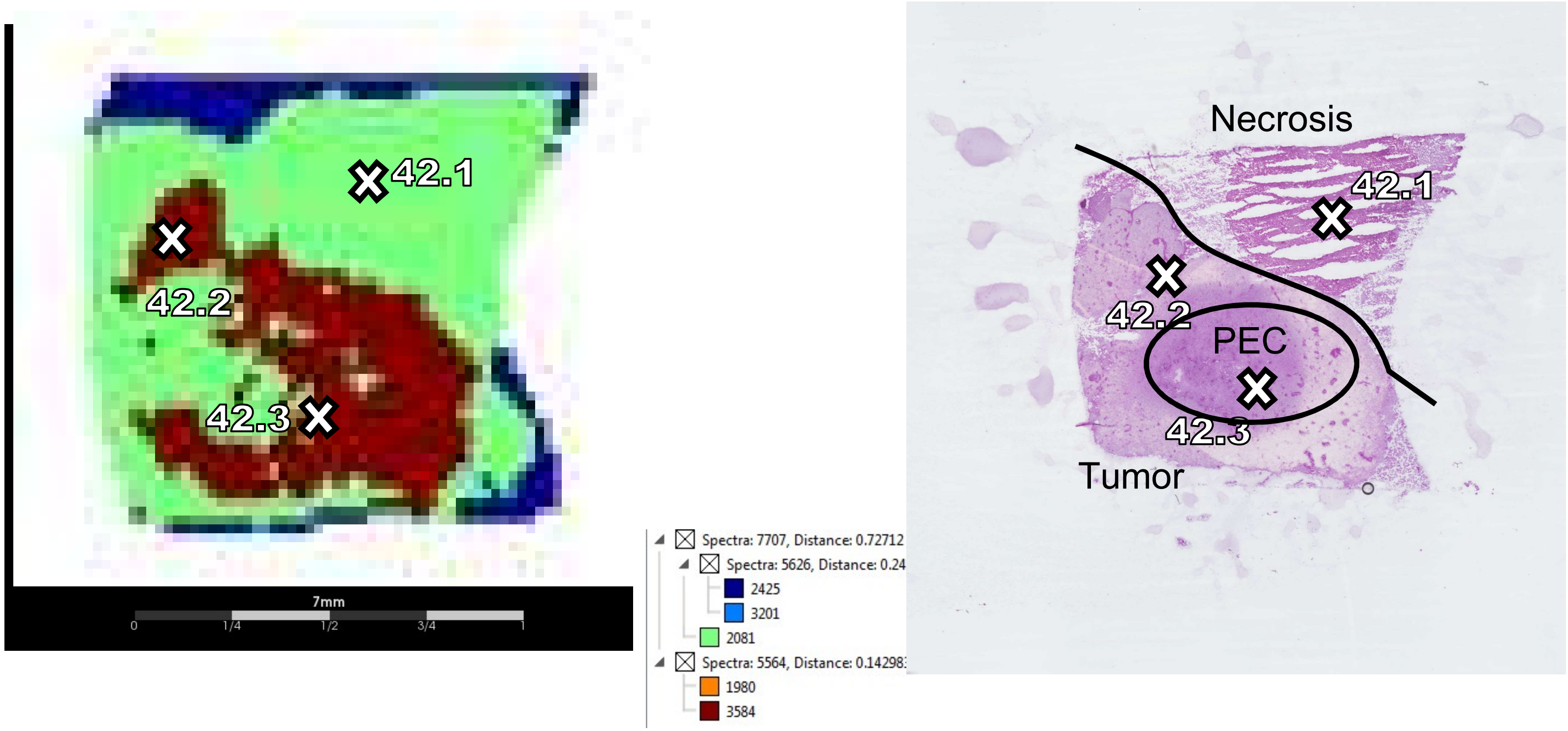
42.1
42.2
42.3
7mm
0
1/4
1/2
3/4
1
Spectra: 7707, Distance: 0.72712
Spectra: 5626, Distance: 0.24
2425
3201
2081
Spectra: 5564, Distance: 0.14298
1980
3584
Necrosis
42.1
42.2
PEC
42.3
Tumor

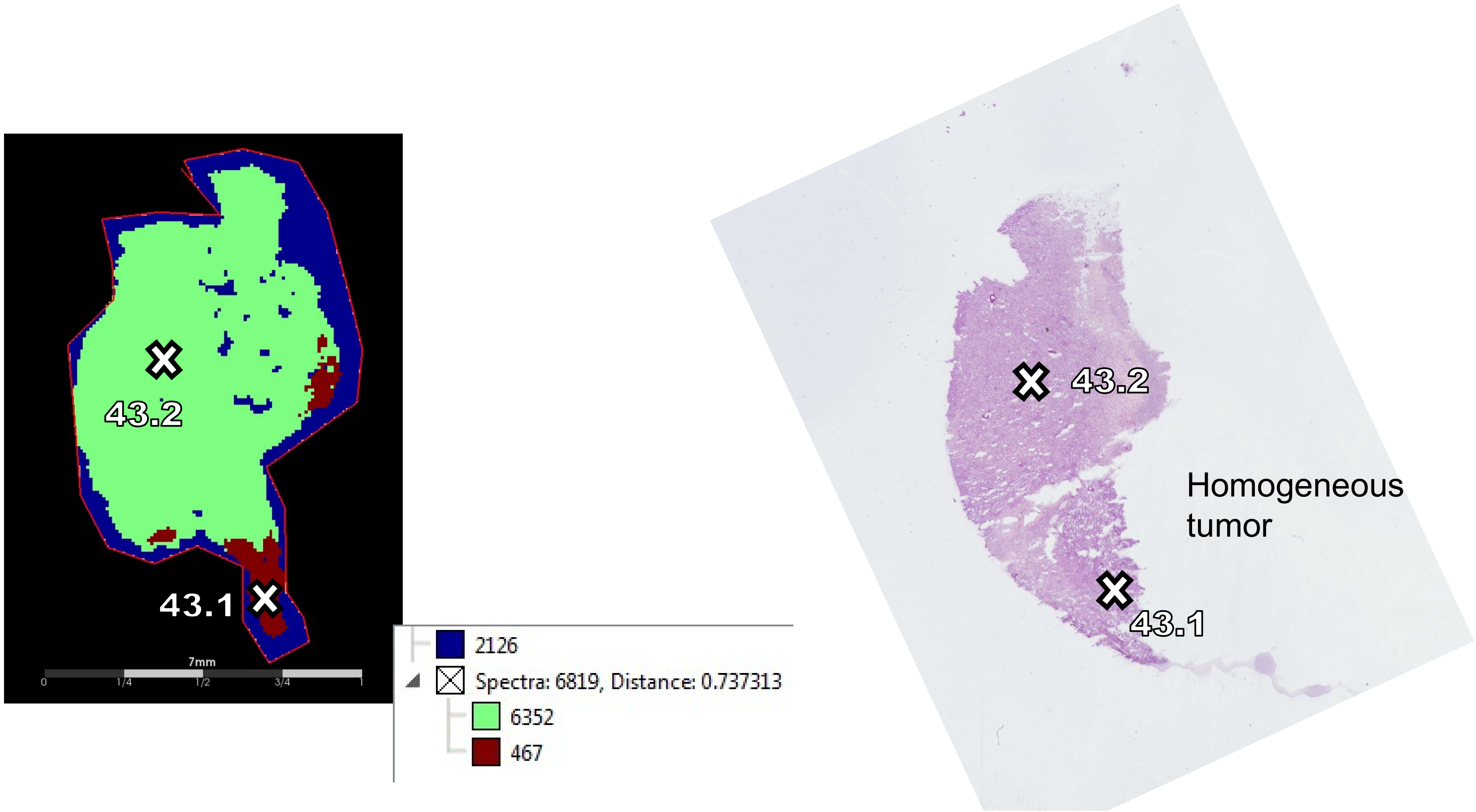
43.2
43.1
7mm
0
1/4
1/2
3/4
1
2126
Spectra: 6819, Distance: 0.737313
6352
467
43.2
Homogeneous tumor
43.1

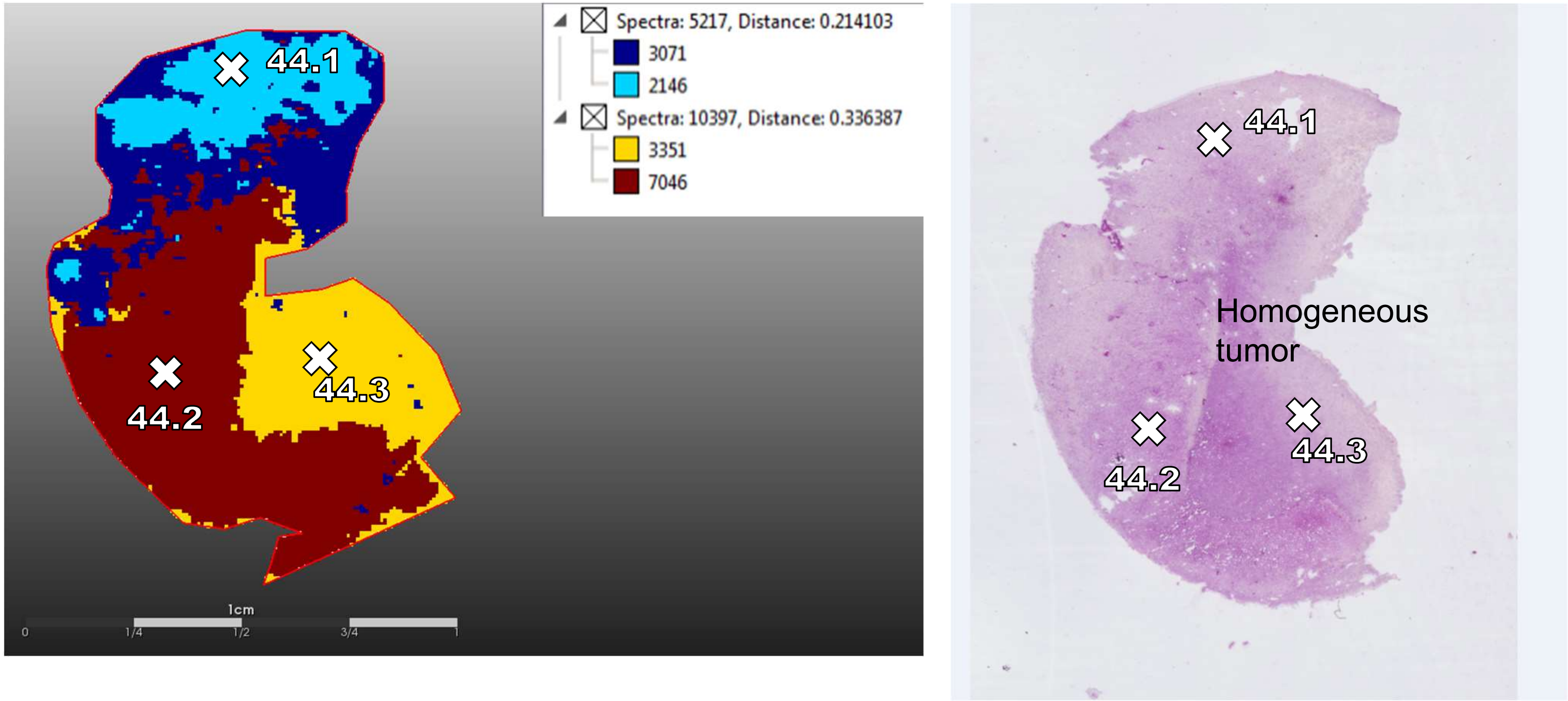
44.1
44.2
44.3
Spectra: 5217, Distance: 0.214103
3071
2146
Spectra: 10397, Distance: 0.336387
3351
7046
1cm
0
1/4
1/2
3/4
1
44.1
Homogeneous
tumor
44.2
44.3

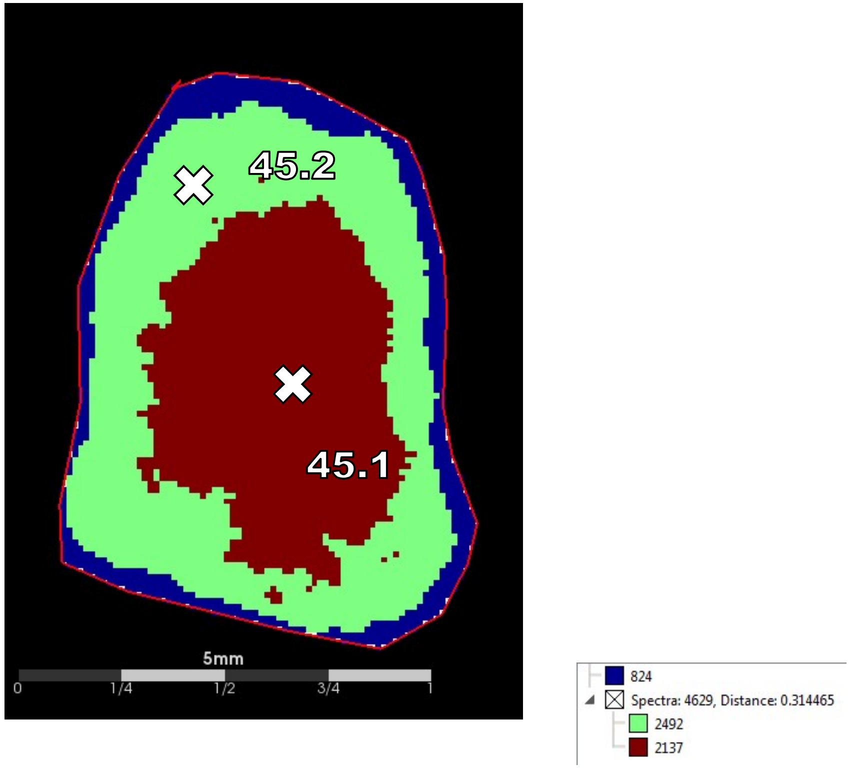
45.2
45.1
5mm
0
1/4
1/2
3/4
1
824
Spectra: 4629, Distance: 0.314465
2492
2137

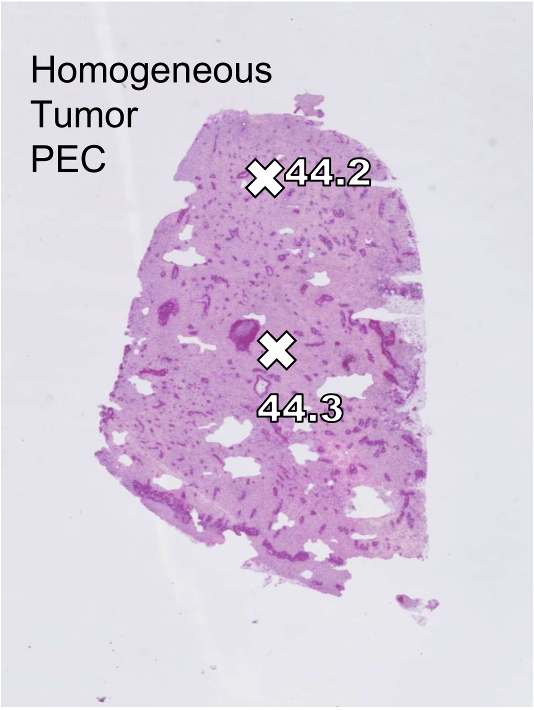
Homogeneous
Tumor
PEC
44.2
44.3

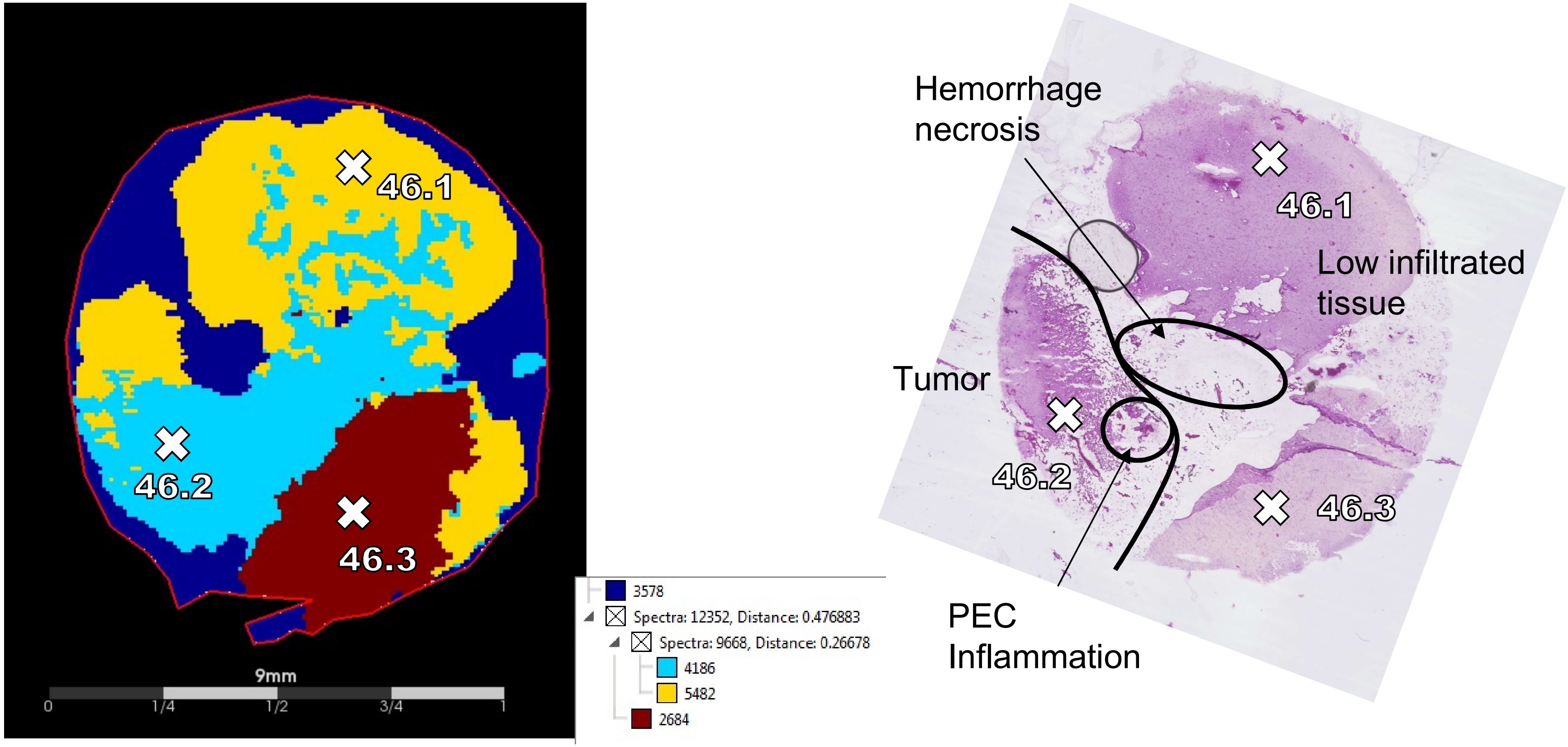
46.1
46.2
46.3
9mm
0
1/4
1/2
3/4
1
3578
Spectra: 12352, Distance: 0.476883
Spectra: 9668, Distance: 0.26678
4186
5482
2684
Hemorrhage
necrosis
46.1
Low infiltrated
tissue
Tumor
46.2
46.3
PEC
Inflammation

41

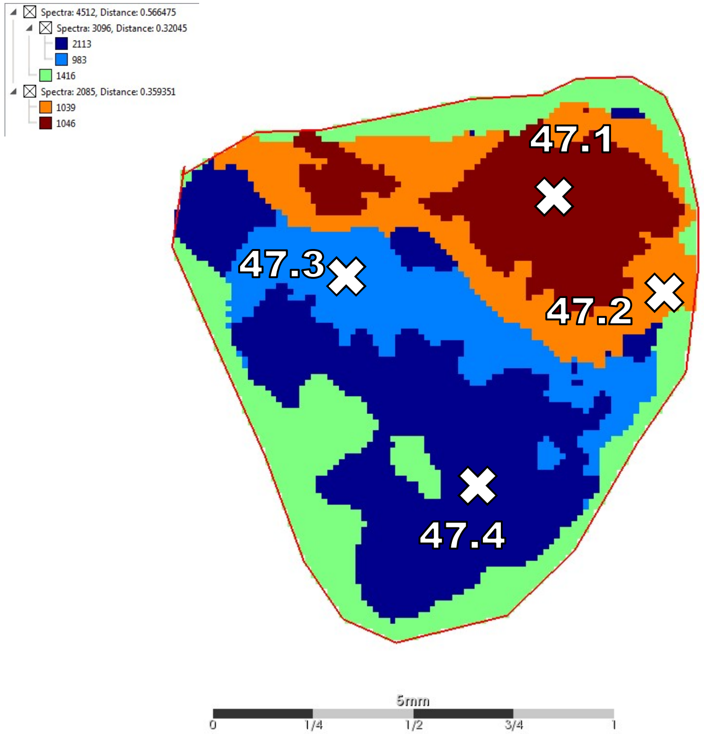
Spectra: 4512, Distance: 0.566475
Spectra: 3096, Distance: 0.32045
2113
983
1416
Spectra: 2085, Distance: 0.359351
1039
1046
47.1
47.3
47.2
47.4
5mm
0
1/4
1/2
3/4
1

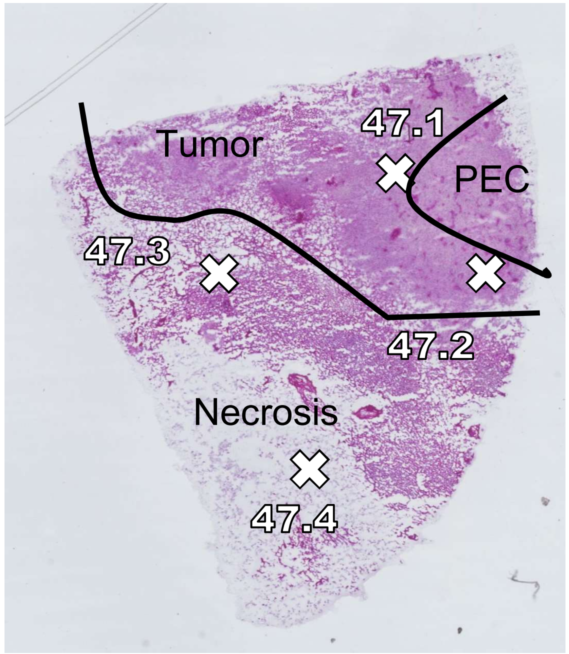
Tumor
47.1
PEC
47.3
47.2
Necrosis
47.4

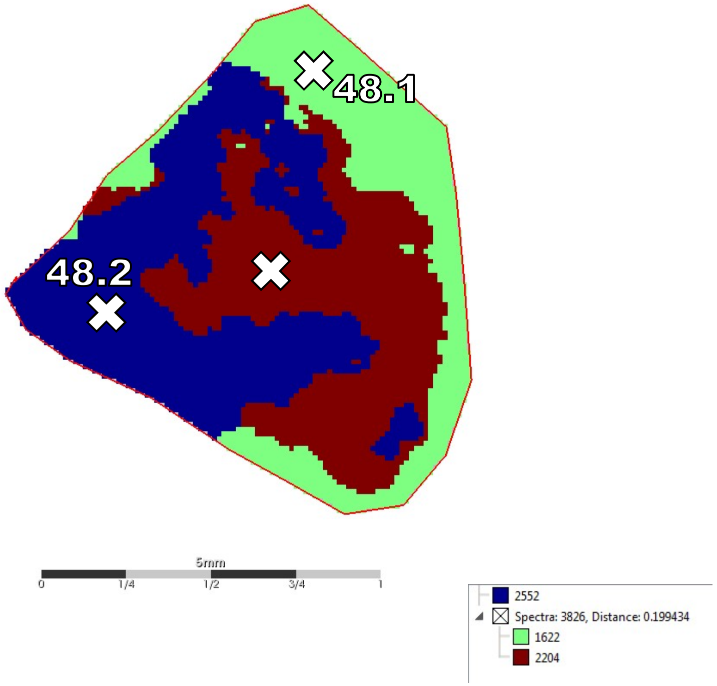
48.1
48.2
5mm
0
1/4
1/2
3/4
1
2552
Spectra: 3826, Distance: 0.199434
1622
2204

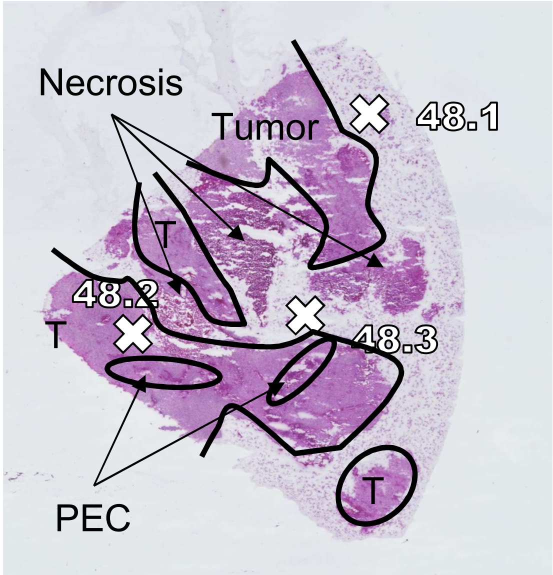
Necrosis
Tumor
48.1
T
48.2
T
48.3
T
PEC

43

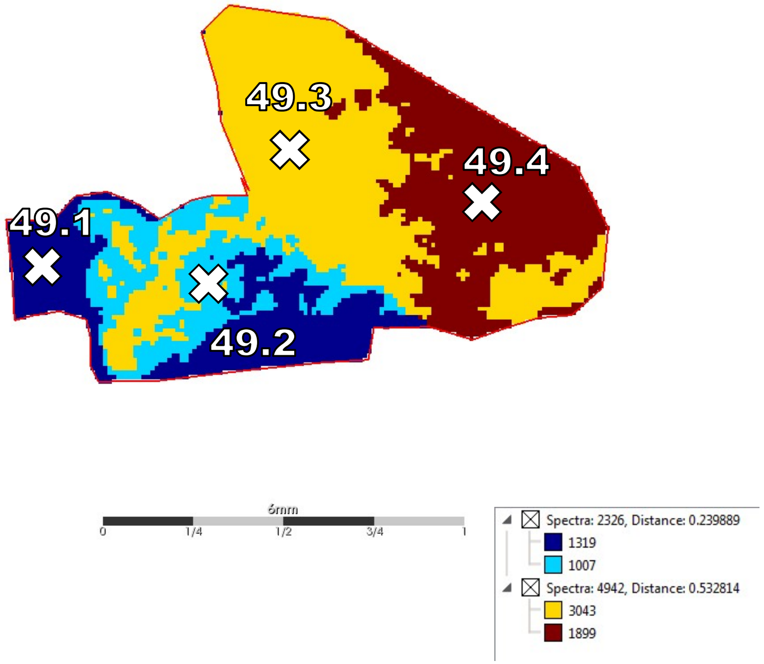
49.3
49.4
49.1
49.2
6mm
0
1/4
1/2
3/4
1
Spectra: 2326, Distance: 0.239889
1319
1007
Spectra: 4942, Distance: 0.532814
3043
1899

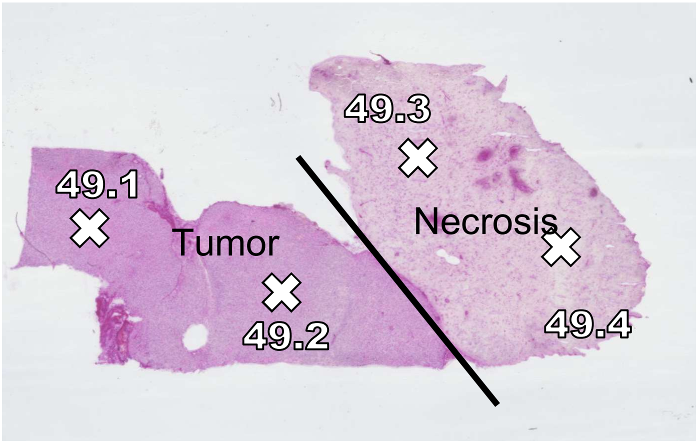
49.3
49.1
Tumor
Necrosis
49.2
49.4

44

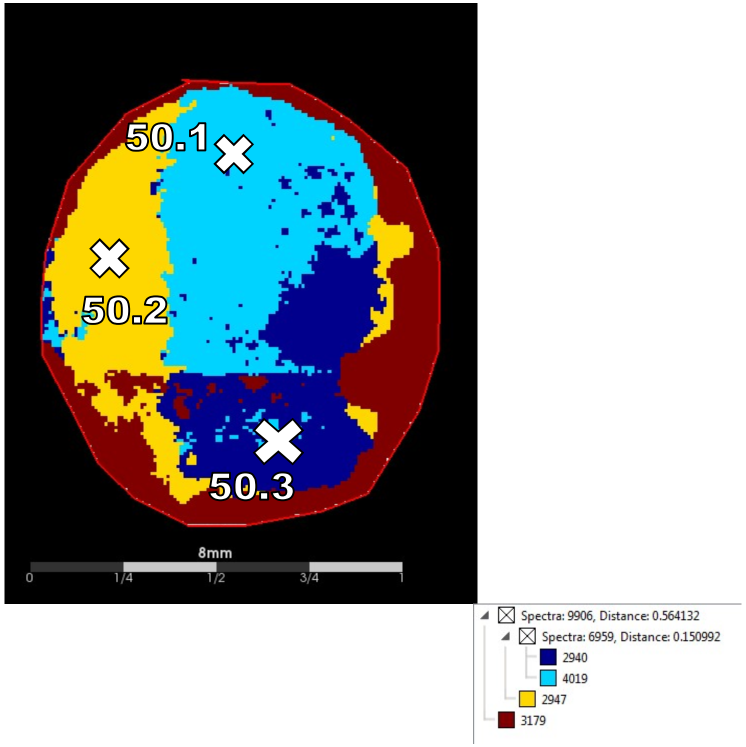
50.1
50.2
50.3
8mm
0
1/4
1/2
3/4
1
Spectra: 9906, Distance: 0.564132
Spectra: 6959, Distance: 0.150992
2940
4019
2947
3179

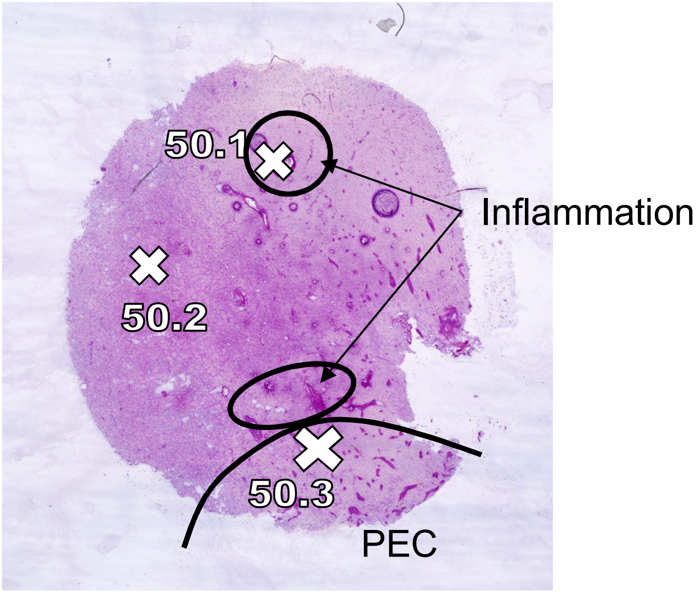
50.1
50.2
50.3
Inflammation
PEC

45

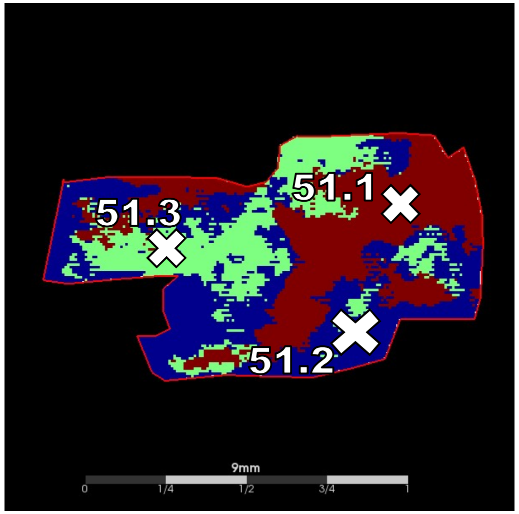
51.1
51.3
51.2
9mm
0
1/4
1/2
3/4
1

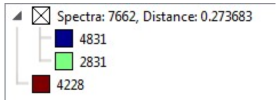
Spectra: 7662, Distance: 0.273683
4831
2831
4228

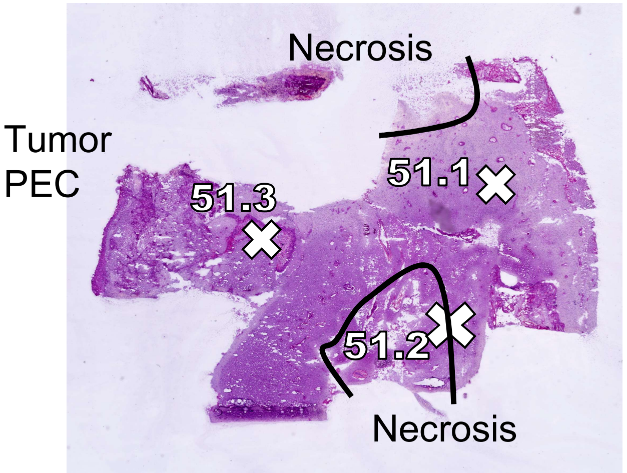
Necrosis
Tumor
PEC
51.1
51.3
51.2
Necrosis

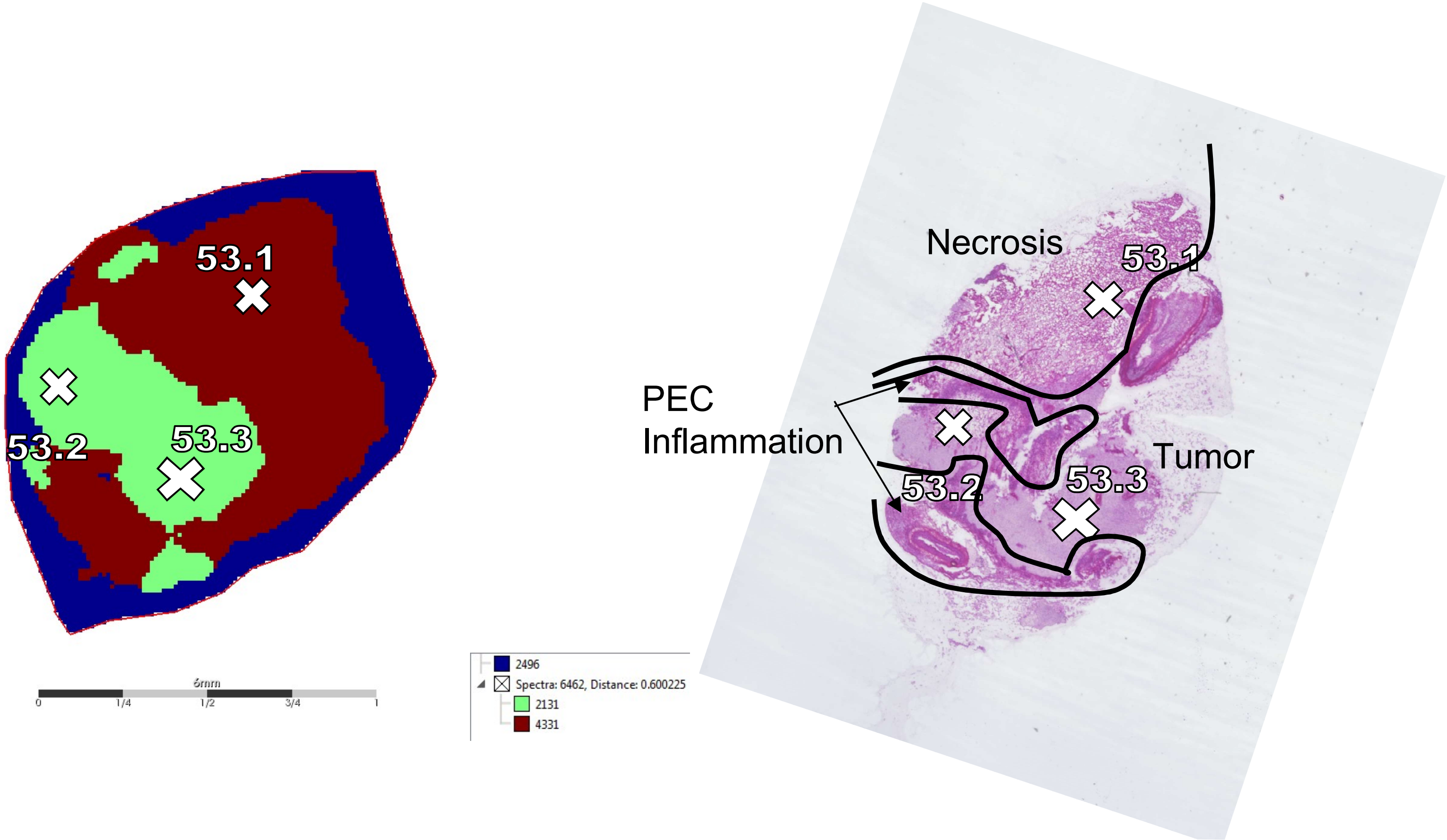
53.1
53.2
53.3
0
1/4
1/2
3/4
1
2496
Spectra: 6462, Distance: 0.600225
2131
4331
Necrosis
PEC
Inflammation
Tumor
53.1
53.2
53.3
